# CancerSTFormer enables multi-scale analysis of spot-resolution spatial transcriptomes and dissects gene and immune regulatory responses to targeted therapies

**DOI:** 10.64898/2025.12.22.696102

**Authors:** Benjamin Strope, Dana Varghese, William Bowie, Stacy Wang, Qian Zhu

## Abstract

The growing number of spot-resolution sequencing based spatial transcriptomic (ST) datasets provides an unprecedented opportunity to study multicellular spatial niches driving cancer transitions. However studying niche-level behavior of tumors remains challenging as it requires a multi-scale approach to modeling the spatial niches and the ability to predict possible effects of genetic perturbations on spatial niches. We propose CancerSTFormer, consisting of a pair of spatially aware transcriptomic foundation models to accommodate niche modeling at different length scales. These models, at the 50µm-Local and 250µm-Extended scales, possess unique capabilities to recover ligand-target gene relationships, niche-specific differentially expressed genes, and organ-specific metastasis associated genes in diverse cancer applications. CancerSTFormer can also reveal the regulatory effects of immune-checkpoint blockade therapies, and other targeted therapies, on patients’ tumors through perturbation analysis, and accurately recapitulated perturbation responses from a spatial Perturb-map experiment. By reusing existing spot-resolution ST studies at scale, this tool transforms the vast spot-resolution ST data into a resource for understanding how gene perturbation impact spatial niches in cancer, while also providing ST-driven, gene-based refinement of treatment-resistance and sensitivity signatures derived from existing bulk transcriptomic studies, enhancing signature interpretation.

## Introduction

Cancer progression and therapy resistance are inherently spatial processes, shaped by interactions between malignant populations, immune infiltrates, and stromal components within the tumor microenvironment^1^ (TME). Understanding these spatially organized signals is critical for uncovering molecular drivers of cancer transitions^2,3^, such as metastasis^4,5^, predicting patient responses to therapy^6^, and understand individual differences likely contributing to tumor formation^7^. Spatial transcriptomics (ST) technologies^8–13^ now provide a means of mapping the native spatial organization and molecular landscape of tumors. Among current sequencing-based ST technologies, including Visium^8^, Slide-seq^9,14^, DBiT-seq^10^, Stereoseq^15^, and others^16–18^, which are known for their untargeted ability to profile whole transcriptomes, spot-resolution is the most common variation of ST. Visium typically uses 50µm spots, while DBiT-seq offers 10µm and 50µm. Each 50µm spot captures 10–20 cells, representing a small tumor microenvironment. Being the most economical approach to ST, and with its widespread adoption across institutions, spot-resolution ST data have been growing at a rapid pace in number and diversity. Despite the data accumulation, this vast resource of publicly available ST data remains untapped for answering various cancer biology questions.

Foundation models^19,20^, such as GPT-based generative models^21,22^ and BERT models^23^, have displayed the promise of synthesizing new biological insights from exceedingly large knowledge bases or biological corpuses, going beyond query-based search systems^24^, and classification-focused deep learning models^25^. In single-cell genomics, its versatility has been demonstrated in the works of scGPT^26^, Geneformer^27^, scFoundation^28^, scBERT^29^, and others^30,31^, in applications ranging from zero-shot learning to transfer learning across a diverse range of tasks. Currently, single-cell RNAseq foundation models lack the important spatial information required to investigate the TME, and are thus limited in studying the cell-extrinsic effects of the tumors. A consequence of this is that single-cell foundation models are less capable of predicting tumor microenvironment effects induced by genetic perturbation, a computational problem known as *in silico* gene perturbations. To this date, there is no foundation model-based framework for **niche-level** *in silico* perturbation – addressing this problem would enable researchers to ask how molecularly targeted therapies alter the multicellular neighborhoods that shape tumor behavior. Although recent work Nicheformer^32^ builds a foundation model for ST data, it does not consider *in silico* gene perturbation, as it focuses on imaging ST with limited gene panels (300-500 genes), and excludes sequencing ST (**Figure 1a**). Implementing *in silico* gene perturbation requires whole-transcriptome sequencing ST like Visium, highlighting the need for a new tool.

**Figure 1:**
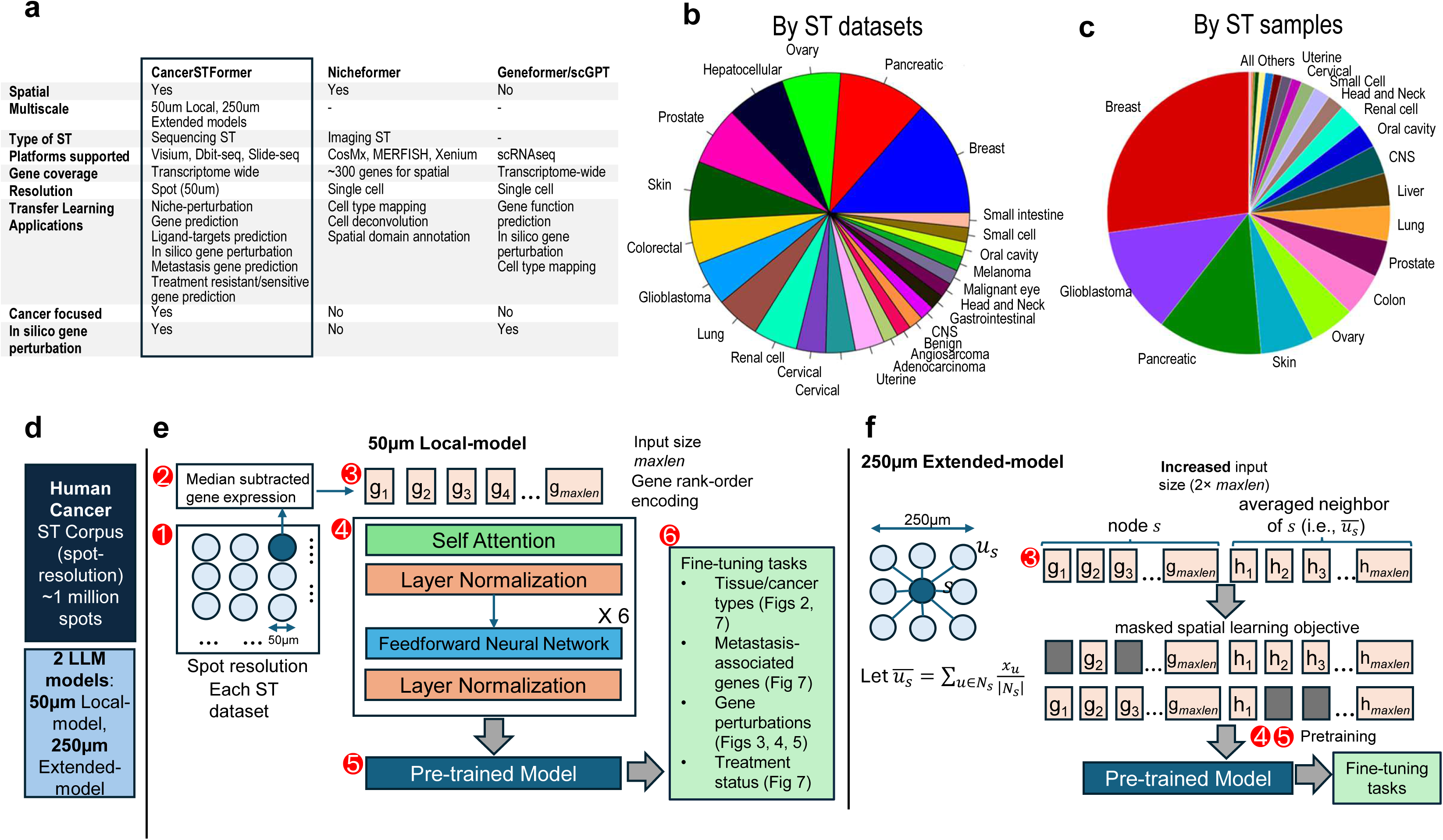
Diversity of cancer spatial transcriptomes and CancerSTFormer overview. **a**. Differences between CancerSTFormer, Nicheformer, and Geneformer/scGPT. **b-c**. Categorization of cancer spatial transcriptomes by the number of ST datasets and samples in each tissue type. **d.** Two spot-resolution spatially aware foundation models being built. **e.** The 50µm-Local Model, known for the spot (50µm diameter)-based modeling. The spot-by-gene matrices of all ST samples first undergo a global median normalization (steps 1-2). Each spot is tokenized to a rank-order encoding sequence, with input size *maxlen* (step 3). Then spots are fed into a large-language model for self-supervised pretraining (steps 4-5). Fine-tuning enables a variety of cancer objectives (step 6). **f.** The 250µm-Extended Model, which models the larger 250µm spot neighborhood. This incorporates the rank order encodings of the self-node and averaged neighbor node into an increased input size of 2x *maxlen*. Masked spatial learning objective then allows self-to-neighbor cross-learning. *Maxlen* is set to 2048.

Although there are non-foundation model-based perturbation tools, they are either focused on single-cell RNAseq (scGen^33^, GEARS^34^), limited to spatial proteomics (Morpheus^35^), or in spatial transcriptomics but not multi-scale, niche-aware, and has unclear relevance to patient-level perturbation responses (Celcomen^36^, River^37^). In contrast, foundation models are expected to produce more generalizable response gene predictions due to pretraining on a large diverse corpus and having the option to fine-tune to specific domains.

Modeling niche-level behavior is difficult as it requires thinking beyond individual cells. We reason that spots themselves are meaningful modeling units, representing multicellular niches where juxtacrine and paracrine signaling^38^ shape tumor phenotypes. Namely, we hypothesize that each 50µm ST spot represents a multicellular niche that provides a local snapshot of the TME, capturing short-range juxtacrine interactions. In contrast, paracrine signaling happens at longer spatial length scales, whereby larger neighborhood effects exist between adjacent spots and spots at a distance within a tumor. These requirements highlight the need for a framework that leverages the abundance of spot-resolution ST, preserves the full complexity of multicellular niches, and enables mechanistic, multiscale interrogation of the TME. Harnessing both local and neighborhood information requires spatially aware foundation models capable of learning across scales.

In this work, we propose to create a spatially aware molecular large-language model, CancerSTFormer, suitable for gene-centric, spot-centric predictions, and spatial *in silico* gene perturbation studies from spot-resolution cancer ST data. CancerSTFormer consists of two models, 50µm Local and 250µm Extended, to account for different length scales. Namely, the larger spatial-scale model (250µm Extended) will account for the neighborhood effect between adjacent spots in ST data, while the smaller 50µm Local model offers the intrinsic ability to model spatial gene expression variation resulting from short-range interactions. We evaluated the model’s ability to retrieve ligand-targets curated by existing database, spatially coexpressed genes and niche-specific DE genes derived from external single-cell resolution spatial Xenium ST data. These results provide the evidence that large-scale analysis of spot-resolution ST data can rival or exceed the analysis of few single-cell resolution spatial ST data, in ligand-target prediction and other tasks, reducing the need for strict reliance on single-cell spatial data. We demonstrate spatial *in silico* gene perturbation in the TME through perturbing *CD274*, *PDCD1*, and *CTLA4*, simulating the anti-PD1, PDL1, and CTLA4 therapies on human TNBC tumors. We show how this analysis can reveal novel immunosuppressive mechanisms that might be imposed by immunotherapy, exposing immunotherapy targets that can be further prioritized for future studies. We also illustrate the CancerSTFormer’s ability to retrieve organ-specific metastasis associated genes, classify ST spots as responders and nonresponders, in various fine-tuning applications. CancerSTFormer is thus a valuable computational tool highly complementary to the growing landscape of ST tools focused on cancer research.

## Results

CancerSTFormer is a sequencing-based, spot-resolution based spatial transcriptomic foundation model trained on over 1,000,000 spots from 50 human cancer studies and 511 samples (**Figure 1b-d**), covering a diverse range of human malignancies. The compendium also encompasses multiple sequence-based ST platforms, such as Visium, DBiT-seq, and Slide-seq (**Supplementary Data 1**). Consistent with previous works^27^, CancerSTFormer adopts a rank-based encoding scheme, in which the expression profile of each spot is tokenized into a series of up to *maxlen* gene tokens (*maxlen* is set to 2048), and adopts self-supervised masked learning during pretraining. Different from previous single-cell based works^27^, we created two modeling frameworks to capture spatial context at 50µm Local and 250µm Extended length scales (**Figure 1e-f**). In the former case, 50µm is the size of each spot, which summarizes both cell-intrinsic information and local neighborhood of 10-20 cells. Rank-based encodings of individual spots are fed into a BERT architecture that treats spots independently, thereby learning representations of diverse small-scale tumor niches. The 250µm Extended model, in contrast, incorporates large-scale spatial neighborhoods by integrating the tokenized vector of the central spot with that of its averaged neighbor in a longer 2×*maxlen* token vector. Such a spatial tokenization scheme, based on concatenated representations, echoes the approach used in DeepSpot^39^. During spatial masking, 15% of the spot’s tokens and 15% of the neighbor’s tokens are withheld, enabling cross-learning from spatial neighbors to individual spots and vice versa. Importantly, no cell type or tumor type annotations are provided during pretraining, allowing the model to infer structure directly from data.

### Zero-shot learning recovers major cancer types

Each model is pre-trained on the full corpus using an architecture incorporating self-attention layers and feedforward layers. Following pretraining, embeddings can be created for new datasets, onto which user-provided annotations (i.e., tumor type) can be overlaid for independent evaluation. To survey the global landscape of cancer spatial transcriptomes, we embedded 100,000 randomly selected spots with the CancerSTFormer-50µm Local model (**Figure 2**). Remarkably, the resulting embedding shows that spots belonging to the same or similar cancer types are highly clustered together, despite the absence of such labels during pretraining (**Figure 2, Supplementary Fig 1**). The embedding further revealed that spots from independent studies of the same cancer type clustered together, suggesting that pretraining can already overcome batch effects across studies within a platform (**Figure 2**). Besides cancer types, we also observed clustering by sample preservation method, with fresh/OCT frozen and FFPE samples forming partially distinct clusters. This sample preparation bias can be mitigated through balanced fine tuning or a batch correction method such as Combat to remove the covariate (i.e., preservation) while retaining cancer-type variation (**Supplementary Fig 2**).

**Figure 2:**
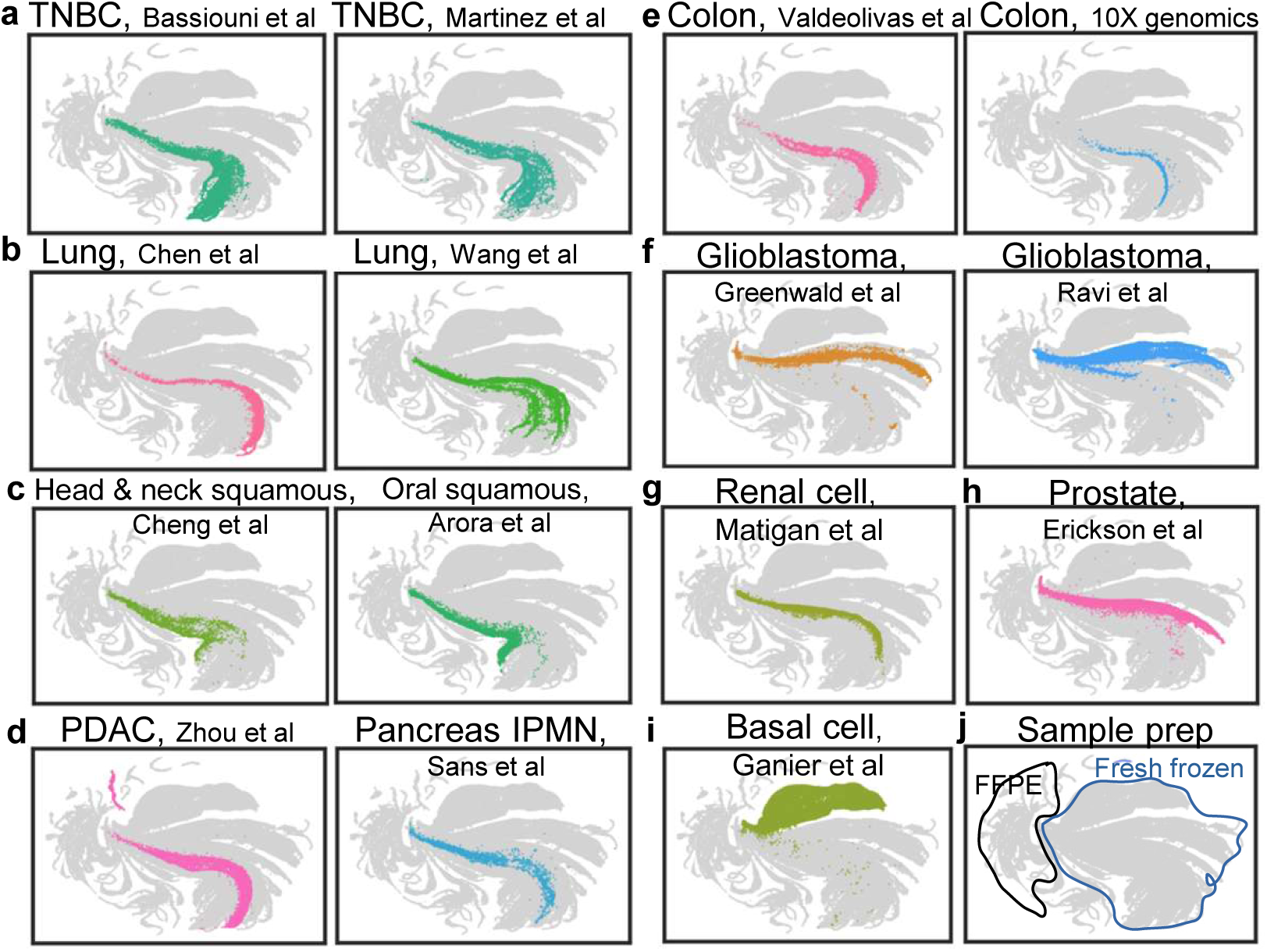
The embedding, pretrained without cancer-type labels, shows distinct clustering when cancer type is overlaid for validation. **a-f.** For each cancer type, spots from two representative studies were chosen to show their co-localization on the UMAP plot. **g-i.** Single representative studies showing renal cell carcinoma, prostate, and basal cell carcinoma spots clustering. CancerSTFormer thus enables meaningful cross-study, cross-tissue comparisons. **j.** Division of spots based on sample preparation.

### A systematic evaluation of retrieval of ligand-targets in the pretrained setting

The duo model set up is suitable for discovering both long-range ligand-target relationships and short-range spatially coexpressed genes. In the former case, a ligand is expressed in the sender, while target genes represent transcriptional targets that are activated in receivers as result of ligand-receptor binding. Targets are discovered by *in silico* perturbing the ligand in spatial ST data, with distinct perturbation implementations for the 50µm Local and 250µm Extended models (see **Methods**). **Figure 3a-b** shows a schematic of the *in silico* gene perturbation procedure for ligand-targets retrieval. Perturbation generates a ranked list of affected genes, or putative transcriptional targets, which we evaluated against curated ligand-target gene sets. To perform a systematic evaluation, we compiled over 1,000 ligand-target gene-sets curated in the NicheNet database^40^ as gold-standard. For each ligand, we perturbed the ligand and assessed the ability of the method to retrieve the associated targets of that ligand. Comparisons included CancerSTFormer (50µm and 250µm variations), Geneformer^27^, and retrieval using single-cell resolution datasets (Visium HD and Xenium-5K) (see **Methods**).

**Figure 3:**
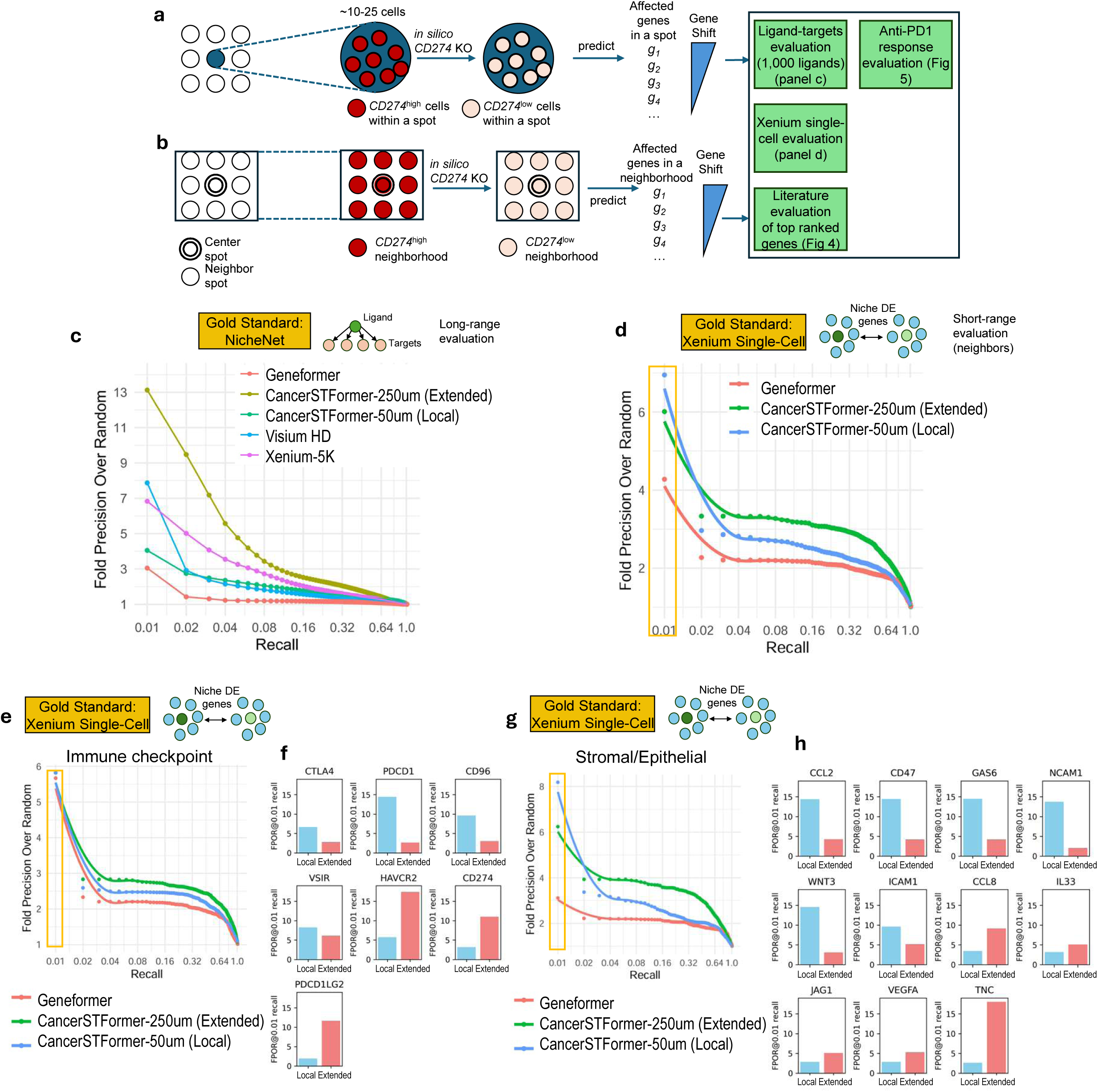
Systematic evaluation of CancerSTFormer in the ligand-targets retrieval task in the pretrained setting. **a-b.** Overview of perturbations in 50µm-Local and 250µm Extended models, with *CD274* (encoding PD-L1 protein) as example. *In silico* deletion removes *CD274* expression, turning *CD274*^high^ spot into a *CD274*^low^ spot. Affected genes are prioritized based on gene shift in the gene embedding space. Most affected genes represent the targets of that ligand and are evaluated. In the Extended Model, the ligand expression is deleted from the entire 250µm neighborhood. The targets are evaluated using NicheNet ligand-target matrix, single-cell spatial Xenium 5K dataset, and literature curation. **c.** NicheNet-based evaluation. Over 1,200 ligands are included in each precision-over-recall curve, with fold-precision-over-random used to account for differences in baseline precision between models. CancerSTFormer and Geneformer are LLM-based retrieval methods utilizing many spot-resolution spatial transcriptomes or single cell transcriptomes. Visium HD and Xenium-5K are simple retrievals using a few single-cell resolution spatial datasets. **d.** Evaluation of spatially coexpressed (niche-DE) gene retrieval defined from Xenium-5K. Using ligands as queries, niche-DE gene retrieval was evaluated and compared between models. Orange box: retrieval at top recall (top 5-10 genes in ranking), showing close performances of Local and Extended Models. **e-h.** Ligands are divided into immune checkpoint (**e-f**) and stroma/epithelia categories (**g-h**), with summary and individual ligand performances shown. These results illustrate variations between different ligands in terms of distance of influence on the target genes.

For the latter, since no transformer exists for Xenium-5K, we implemented a simple retrieval approach: differentially expressed genes were identified by comparing neighbors of ligand-hi versus ligand-lo cells. The comparison thus enables us to compare scRNAseq-based against ST-based transformers, and spot-resolution ST (CancerSTFormer) against single-cell resolution assays (Visium HD/Xenium-5K). The perturbation context uses TNBC ST or scRNAseq (in the case of Geneformer), which have abundant ligand expressed. In our evaluation results, the CancerSTFormer-250µm Extended model performs best, with Fold Precision Over Random (FPOR) of 13 at 1% recall, markedly outperforming Visium HD or Xenium-5K retrievals. This shows the collective advantage of large-scale LLM spatial modeling on many spot-resolution ST data, which rivals the biological predictions derived from few single-cell resolution spatial data like Visium HD/Xenium-5K. The lower performance could not have been due to the lower cell/bin count as Xenium-5K and Visium HD contain ∼700,000 cells and 120,000 bins under tissue respectively. Importantly, all the spatial data-derived models perform better than Geneformer, which was trained on single-cell RNAseq data with no spatial information.

We hypothesize that NicheNet database may be enriched for long-range interactions while placing less focus on short-range interaction-induced effect. To address this, we generated a separate gold standard, using single-cell resolution Xenium-5K spatial data, to derive niche-DE genes of receiver cells immediately surrounding ligand-high cells. These niche-DE genes may serve as short-range interaction induced genes for evaluation. Evaluation against this new standard of niche-DE genes revealed a much smaller performance gap (**Figure 3d**). Notably, at the recall of top 1% (see orange box **Figure 3d**), the Local model performs better than the Extended model. We further divided the ligands into Immune Checkpoint (IC) category (**Figure 3e**) and Stromal/Epithelial category (**Figure 3g**). In both cases, the short-range 50µm Local model has a higher FPOR at top recall (orange boxes **Figure 3e, g**) than the 250µm Extended model in retrieval of niche-DE genes, exemplified by *CTLA4*, *PDCD1*, *CD96* (in the IC category), and *CCL2*, *CD47*, *GAS6*, *NCAM1*, *WNT3* (in the Stroma/Epithelial category) (**Figure 3f, h**). Overall, the model performances reveal distinct preferences among ligands in mediating local versus extended effects.

### *In silico* gene perturbations simulating anti-PD1, anti-PDL1, and anti-CTLA4 therapy responses in the pretrained setting

Having established that CancerSTFormer retrieves biologically meaningful ligand-target relationships, both long-range and short-range, we next asked whether the same perturbation framework can be used to simulate therapeutic interventions. Because many cancer therapies act by blocking or activating ligand or receptor, such as PD1 and PDL1, *in silico* perturbation of these genes should reveal downstream transcriptional consequences within the spatial TME. This enables us to probe whether targeting a gene is predicted to trigger desirable responses such as activation of antitumor immunity. As proof of principle, we sought to perturb each of genes encoding PD-1, PD-L1, and CTLA-4 proteins, respectively *PDCD1*, *CD274*, and *CTLA4*, for which there exist approved therapies in solid tumors^41–44^ (**Figure 4**). We perturbed the genes using the CancerSTFormer-50µm Local and 250µm Extended models in a set of TNBC ST samples^45,46^. To evaluate the perturbation results, for each perturbation, we extracted the top 10 genes with the greatest shift, given that the FPOR at this low recall level (1%) is the highest. Using SEEK^47^, we next expanded on these 10 genes, then performed gene-set enrichment analysis (**Figure 4**). We find that *PDCD1*, *CD274*, and *CTLA4* deletion responsive genes were enriched in actin-filament processes (P=1×10^-2^), T cell receptor signaling pathway (P=1×10^-5^), and fatty acid catabolism (P=1×10^-2.4^) respectively (CancerSTFormer-50µm Local) (**Figure 4**). For the CancerSTFormer-250µm Extended model, the responsive genes were also enriched in the activation of immune response GO-term, albeit with much higher significance P=1×10^-10^ across all three immune checkpoint (IC) perturbations (**Figure 4**). Altogether, these results suggest that IC perturbations elicit a long-distance anti-tumor immune response (250µm Extended model), and a separate local response (50µm Local model) that is highly connected to immune function.

**Figure 4:**
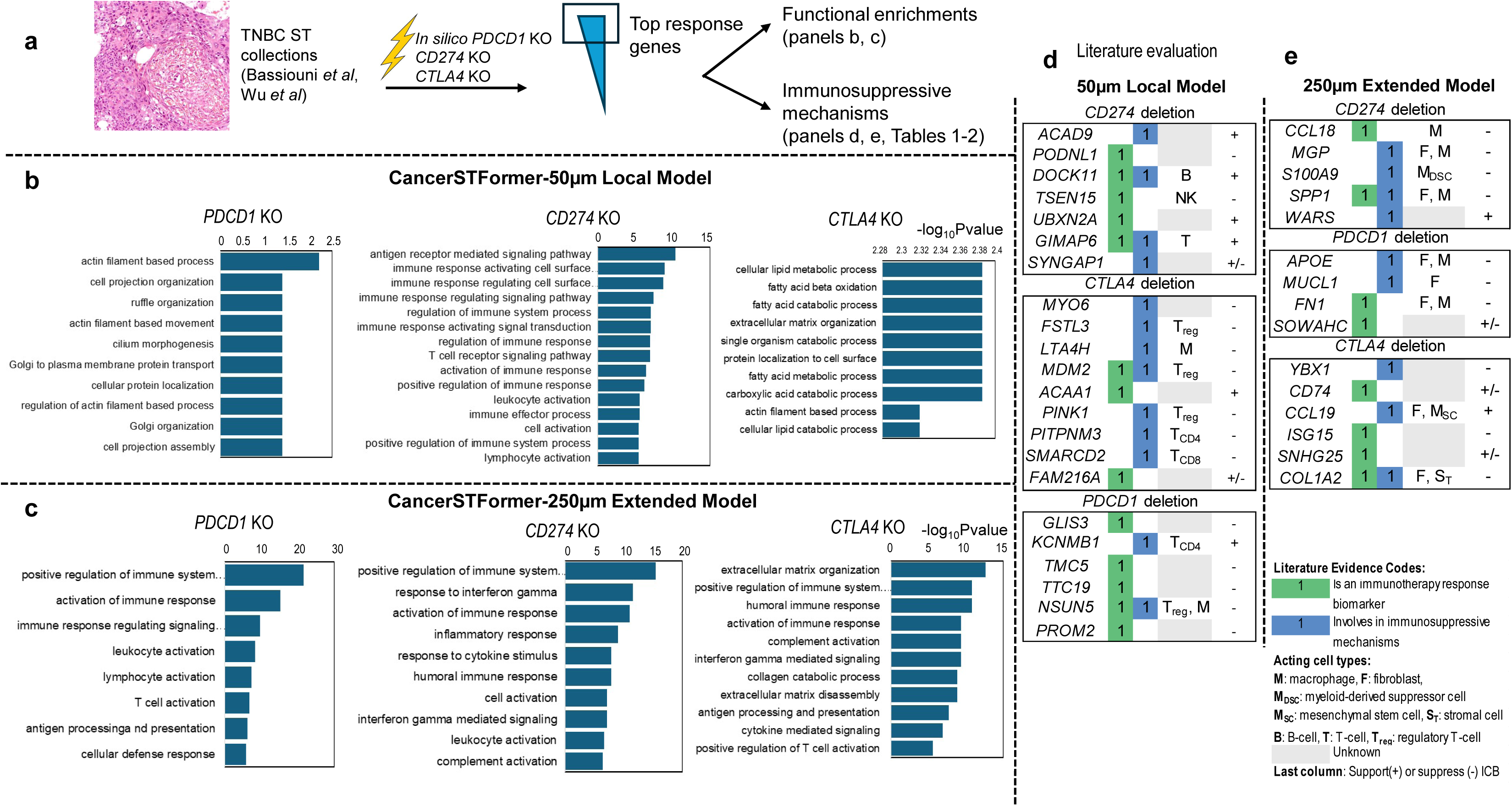
*In silico PDCD1*, *CD274*, and *CTLA4* perturbations reveal responsive genes to immune checkpoint inhibition. **a.** Spatial transcriptomes of TNBC were subject to each immune checkpoint molecule ablation, with most affected genes evaluated by GO-term functional enrichment and by connection with immunosuppression. **b-c.** GO-term enrichment results of top responsive genes to each perturbation (**b.** 50µm-Local model. **c.** 250µm-Extended model). The top 10 genes in each case were expanded using SEEK by coexpression. They were then subject to enrichment via hypergeometric test. In 50µm-Local model, previous research has reported a strong tie between actin cytoskeleton polymerization and T-cell antigen receptor engagement^101^. Fatty acid beta-oxidation is important for the development of CD8+ memory T cells^102^. In the 250µm-Extended model, the extracellular matrix disassembly in CTLA4 deletion has been reported in previous works. **d-e**: Literature evaluation. Each gene was searched for evidence of being an immunotherapy biomarker or being experimentally linked to immune responses to immunotherapies. Full literature search is presented in **Tables 1**–**2**. Acting cell types and the type of responses (boost or suppress) are noted. Pretrained model results were used for this figure.

### *PDCD1*, *CD274*, and *CTLA4* perturbations reveal distance-dependent immunosuppressive mechanisms and potential immunotherapy targets

A literature survey (**Figure 4d**, **Supplementary Table 1**) on the top 10 responsive genes to each IC deletion revealed biomarkers of immunotherapy response, such as *GIMAP6*, *TMC5*, *TSEN15*, *ACAA1*, *GLIS3*, *PROM2*, and *PODNL1* (**Figure 4d**, **Supplementary Table 1**, biomarker column). Expectedly, some genes, including *GIMAP6*, *ACAA1*, and *ACAD9*, are linked to the normal function of IC blockade and promote immune activation. Interestingly and importantly, many others were related to immunosuppression in connection with immunotherapies. Experimental deletion or knockdown of these response genes can reverse immunosuppression and increase immunotherapy efficacy, making these potential immunotherapy targets (**Figure 4d**, **Supplementary Table 1**, mechanistic column). Examples include *MYO6*, *FSTL3*, *LTA4H*, *MDM2*, *PINK1*, *PITPNM3*, *SMARCD2*, and *NSUN5* (**Figure 4c-d**, **Supplementary Table 1**). Other genes such as *DOCK11* and *MDM2* (**Figure 4d**, **Supplementary Table 1**) were related to immune-related adverse events (irAE), such as normocytic anemia and PD1 inhibitor-related hyperprogressive disease. In the predictions made by the CancerSTFormer-250µm Extended model (**Figure 4e, Supplementary Table 2**), some of the predicted response genes were shared across the three perturbations. Again, mechanisms of immunosuppression form a central theme. For example, *MGP* promotes CD8 T cell exhaustion and *S100A9* monocytes can orchestrate MDSC-mediated anti-PD1 immunotherapy resistance (**Figure 4e, Supplementary Table 2**). *SPP1*, a gene of intense interest previously associated with immunotherapy response, is also among the results of *CD274* perturbation (**Figure 4e**, **Supplementary Table 2**). Also, *CD74* is associated with the irAE pneumonitis (**Figure 4e**, **Supplementary Table 2**). Overall, in contrast to the conventional thinking that anti-PD1/PDL1 therapies activate one’s immune system, our *in silico* results showed that they also upregulate immunosuppressive genes that counteract the effectiveness of immunotherapy. CancerSTFormer can reveal such genes as possible immunotherapy targets.

We wonder whether the response genes of the Local and Extended model variants could indicate distinctive immunosuppressive cell types at play at different distances (**Figure 4d-e**, **Supplementary Tables 1-2**, cell-type column). Indeed, the 50µm Local model’s results are more enriched in regulatory T-cells, and other types of T-cells, such as CD4 and naive T cells, contributing to direct-contact mediated immunosuppression. In contrast, the 250µm Extended model’s results are more pointing to macrophages, cancer associated fibroblasts, and extracellular matrix, forming a theme of long-distance stroma-mediated immunosuppression. In summary, these results highlight the unique advantages of each of 50µm and 250µm CancerSTFormer models in capturing multiscale immunosuppressive mechanisms, with implications for predicting immunotherapy success and formulating combination co-targeting strategies.

### Finetuned CancerSTFormer refines bulk RNAseq-derived signatures of treatment-resistant and sensitive genes to achieve biomarker generalizability across patient cohorts

To enable more realistic simulations of immunotherapy response, we fine-tuned CancerSTFormer using gene signatures derived from human subjects who had undergone immunotherapy. The pre-trained model results (**Figures 3**–**4**) already retrieved biologically relevant PD1/PDL1/CTLA4 targets, but we hypothesized that incorporating clinically validated response genes would further improve accuracy of *in silico* gene perturbations. Thus, we downloaded RNA-seq profiles from Wolf *et al*^48^, comprising 60 TNBC patients categorized as pathological complete response (pCR) or no response after pembrolizumab treatment. From these groups, we derived PD1-responsive and PD1-resistant gene sets, which served as gold standards to train a gene-classifier within CancerSTFormer for predicting response-associated genes from ST data (**Figure 5b**). We then applied *in silico* perturbation of *PDCD1* using this fine-tuned model and assessed the results (**Figure 5b**). The same I-SPY2 trial^48^ also included response data for ganitumab (IGF1R inhibitor) and trebananib (Angiopoietin-1 or ANG1/*ANGPT1* inhibitor), permitting us to perform a broader evaluation across multiple treatment modalities.

**Figure 5:**
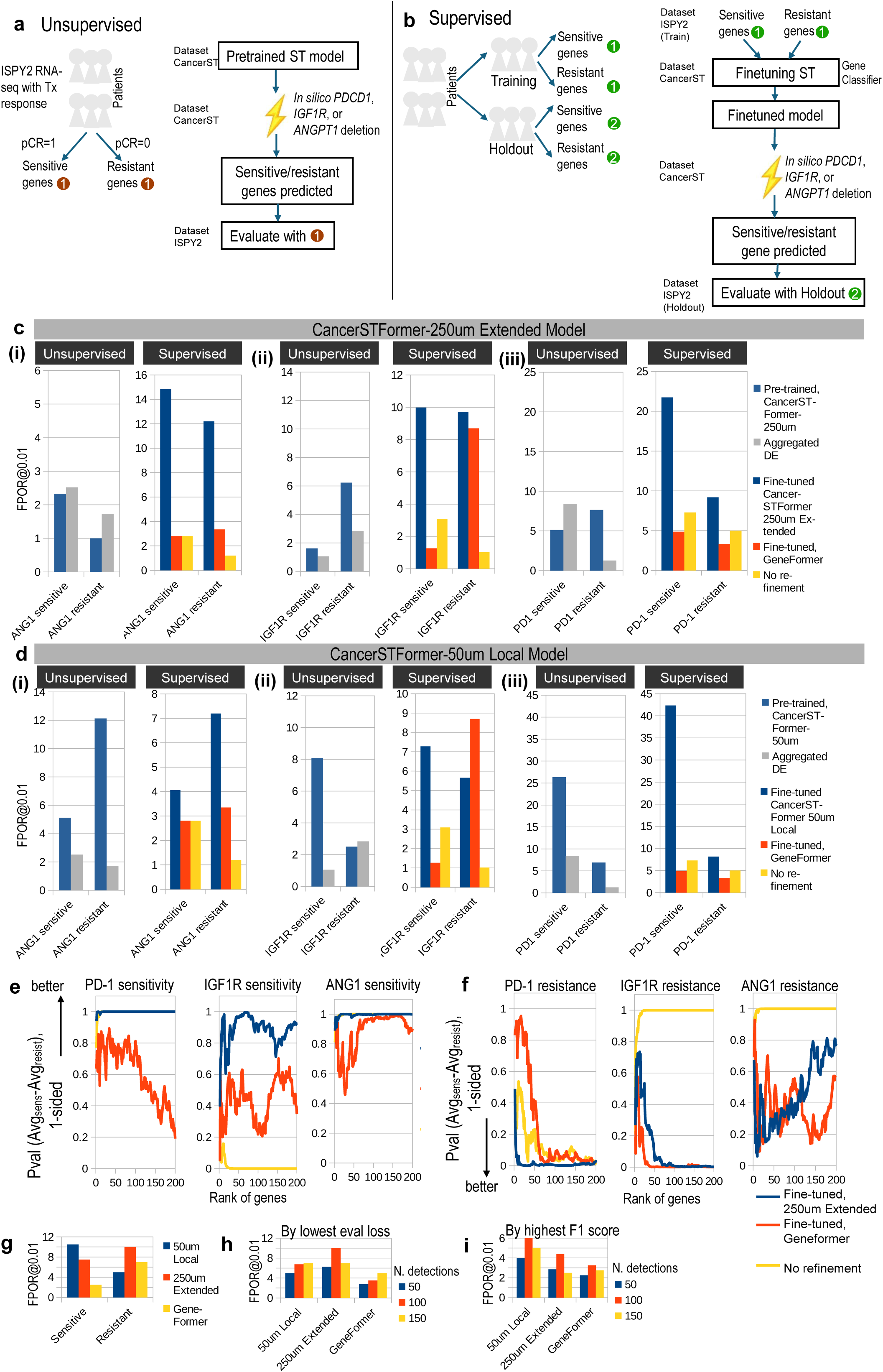
CancerSTFormer, in the finetuned setting, enables prediction of sensitive and resistant genes to targeted therapies, with finetuned model showing enhanced performance. **a-b.** Evaluation strategies. **Unsupervised**: evaluation of pretrained model. 3 targeted therapies investigated: anti-PD1 (pembrolizumab), anti-IGF1R (ganitumab), anti-ANG1 (trebanalib). The molecular gene of interest (*PDCD1*, *IGF1R* or *ANGPT1*) was *in silico* deleted in TNBC ST datasets. Responsive genes to deletions were computed per spot, and were aggregated across spots to form a ranking of top affected genes (according to cosine similarities). For evaluation, top affected genes were compared to treatment-sensitive genes or resistant genes defined on ISPY2 trial patients (based on their RNAseq profiles). <u>Note</u> that ST datasets (on which perturbations were performed) were a different cohort from ISPY2 transcriptomic data. We could still perform this evaluation as we believe in the generalizability of treatment-response gene signatures across patient cohorts. **Supervised**: Patients in the ISPY2 trial were split into Training and Holdout groups. Finetuning proceeds by using the sensitive/resistant genes derived from the Training patient group to train a gene classifier within CancerSTFormer model using its available ST datasets. After training, the fine-tuned model was used for *in silico PDCD1*, *IGF1R*, or *ANGPT1* perturbations. During evaluation, the gene results were compared to the sensitive/resistant genes derived from Holdout patients. **c-d.** Evaluation on the 250µm Extended model (**c**) and 50µm Local model (**d**). Treatments are *ANGPT1*(**i**), *IGF1R*(**ii**), and *PDCD1*(**iii**) inhibition. Aggregated DE is a baseline method compared to the pretrained model in the unsupervised setting. Geneformer in the finetuned case was also compared to the various finetuned models. No refinement is a control where Training and Holdout groups’ sensitive/resistant genes were directly compared against each other. **e-f**. A second evaluation method. For each rank in the perturbation response rank-list, the P-value of differential gene expression was computed up to genes in that rank using the Holdout patients. 1-sided P-values (Avg^resistant^ – Avg^sensitive^) shown. Thus, sensitivity gene prediction should have the highest 1-sided P-value, while resistance gene prediction should have the lowest 1-sided P-value. **g-i**. Robustness across finetuning settings and cutoffs. Aggregate performances are shown based on type of response (**g**), and fine-tuning objective (i.e., lowest eval loss or highest F1-score) (**h-i**). *N. detections* is a parameter related to the minimum number of spots (out of 1,000) that the gene is detected in order to be counted. In all finetuning, we ran parameter discovery with Ray using 60 trials – performances shown are the average of top 5 trials by lowest eval loss (**c-d, h**) or top 5 trials by highest F1-score (**i**). Each trial took 10 minutes (Local Model) or 30 minutes (Extended Model) on an NVIDIA RTX A6000 Ada GPU.

Model evaluation followed two complementary strategies (**Figure 5a-b**). In the **Unsupervised** strategy (**Figure 5a**), we evaluated the results of the pretrained CancerSTFormer directly on treatment-sensitive and -resistant genes derived from all ISPY2 patients. In the **Supervised** strategy (**Figure 5b**), we first fine-tuned the model using a subset of trial patients and evaluated performance on the remaining patient subset (i.e., holdout cohort). As shown in **Figure 5c-d (i, ii, iii, Unsupervised),** the pretrained 50µm Local model already outperformed the baseline Aggregated DE method (see **Methods** and later in this section) in unsupervised evaluations, achieving FPOR@0.01 recall = 25–8 across PD1 (**iii**), IGF1R (**ii**), and ANG1 (**i**) inhibitor treatments. Fine-tuning further enhanced the retrieval of PD1/PDL1-sensitive genes (**iii**) and produced similar gains for IGF1R (**ii**) and ANG1 (**i**) (**Figure 5c-d, i, ii, iii, Supervised**). Improvements over pretrained models were most pronounced for the 250µm Extended model—especially in predicting resistant genes—showing up to 5-fold performance increases (**Figure 5d**). Together, these analyses showcase the differential strengths of pretraining vs. fine-tuning and of Local vs. Extended models in identifying treatment-sensitive and -resistant genes.

Importantly, when compared against a fine-tuned Geneformer and a no-refinement control (**Figure 5c-d, i, ii, iii, Supervised**), both Local and Extended CancerSTFormer fine-tuned variants consistently achieved higher accuracy in predicting resistant and sensitive genes in holdout cohorts across all three treatments. This advantage stems from the closer correspondence between ST and bulk data than between single-cell RNA-seq and bulk profiles. In contrast, directly comparing resistant/sensitive genes between training and holdout patients yielded minimal gene overlap, meaning that the training patients’ resistant/sensitive genes, without refinement, have weak predictive power (“**no refinement”, Figure 5c-d**). Overall, these findings demonstrate the model’s ability to refine bulk-derived treatment-response signatures. The fine-tuned CancerSTFormer successfully aligned ST data with bulk RNA-seq–derived gene sets to generate refined predictive biomarkers that generalized to unseen patients.

Notably, the above results are robust to evaluation methods and parameter settings during fine-tuning. We evaluated the refined biomarker ranking by computing the differential gene expression change and P-value for each gene in the holdout cohort between pCR=1 and pCR=0 patient groups. The discriminatory power of biomarkers, evaluated by computing the significance of difference in expression means of sensitive and resistant populations in holdout, is maintained at the top ranks or throughout the gene ranking [**Figure 5e**, blue line (CancerSTFormer) vs. red line (GeneFormer), the higher the better]. Similarly for resistance prediction, the discriminatory power of genes in CancerSTFormer is superior to Geneformer and the no refinement control (**Figure 5f**, blue vs red line, the lower the better). We also varied the finetuning settings. Fine-tuning runs were discovered by optimizing an objective like minimizing evaluation loss or maximizing macro-F1 score, across 60 trial runs. Results have shown that regardless of the objective used to select fine-tuning runs, the superiority of CancerSTFormer over other foundation models were maintained, illustrating the robustness of our results (**Figure 5h-i**).

CancerSTFormer also outperforms a non-LLM baseline. Here, in a baseline method, termed Aggregated DE, each ST sample was queried by selecting ligand-high and ligand-low spots, followed by DE analysis (t-test) on their log-transformed expression values. DE results from all samples were then aggregated using Stouffer’s method to generate a master ranked list of genes for evaluation. In the ligand-retrieval task, CancerSTFormer substantially outperforms Aggregated DE in both mean FPOR and median FPOR across 1,000 ligand-target gene-sets, as shown by precision-recall curves (**Supplementary Fig 3a-b**). In the *in silico* perturbation task, CancerSTFormer performs better than Aggregated DE on average (**Supplementary Fig 3c-e**). Overall, these results confirm the advantage of transformer models compared to simple baseline models.

### Prediction of perturbation responses in Perturb-map experiments

Dhainaut *et al* used Perturb-map^49^, a technology that combines imaging and spatial transcriptomics from an in vivo CRISPR experiment, in a mouse model to identify regulators of lung cancer TME. Among these, Tgfbr2 knockout tumor clones were frequently appearing and the authors performed ST to understand the responses to Tgfbr2 deletion (**Figure 6a**). We can train a CancerSTFormer-Extended model to predict knockout responses across four clones (Tgfbr2_1, Tgfbr2_2, Tgfbr2_4-1, and Tgfbr2_4-2) by training on one clone and testing/evaluating on each of the remaining three. In each case, KP (KRAS^G12D^p53^null^) tumors without receiving Tgfbr2 CRISPR guides served as unperturbed controls. **Figure 6b** details the gene-based training and evaluation procedure that involves identifying DE up- and down-regulated training genes first. For testing, unperturbed KRAS tumors of a given sample were perturbed in silico and responses compared to Tgfbr2 KO clones from the ST profiles of the same sample. Results of our evaluation revealed that CancerSTFormer-Extended possesses the ability to predict Tgfbr2 KO up-regulated genes (FPOR of 10-13), and down-regulated genes (FPOR of 25) (**Figure 6c**). These results are vastly superior to Geneformer’s results, and are between 3-5 fold better compared to Geneformer (**Figure 6d**). Thus, we conclude that CancerSTFormer will be easily adaptable to train from spatial perturbation datasets.

**Figure 6:**
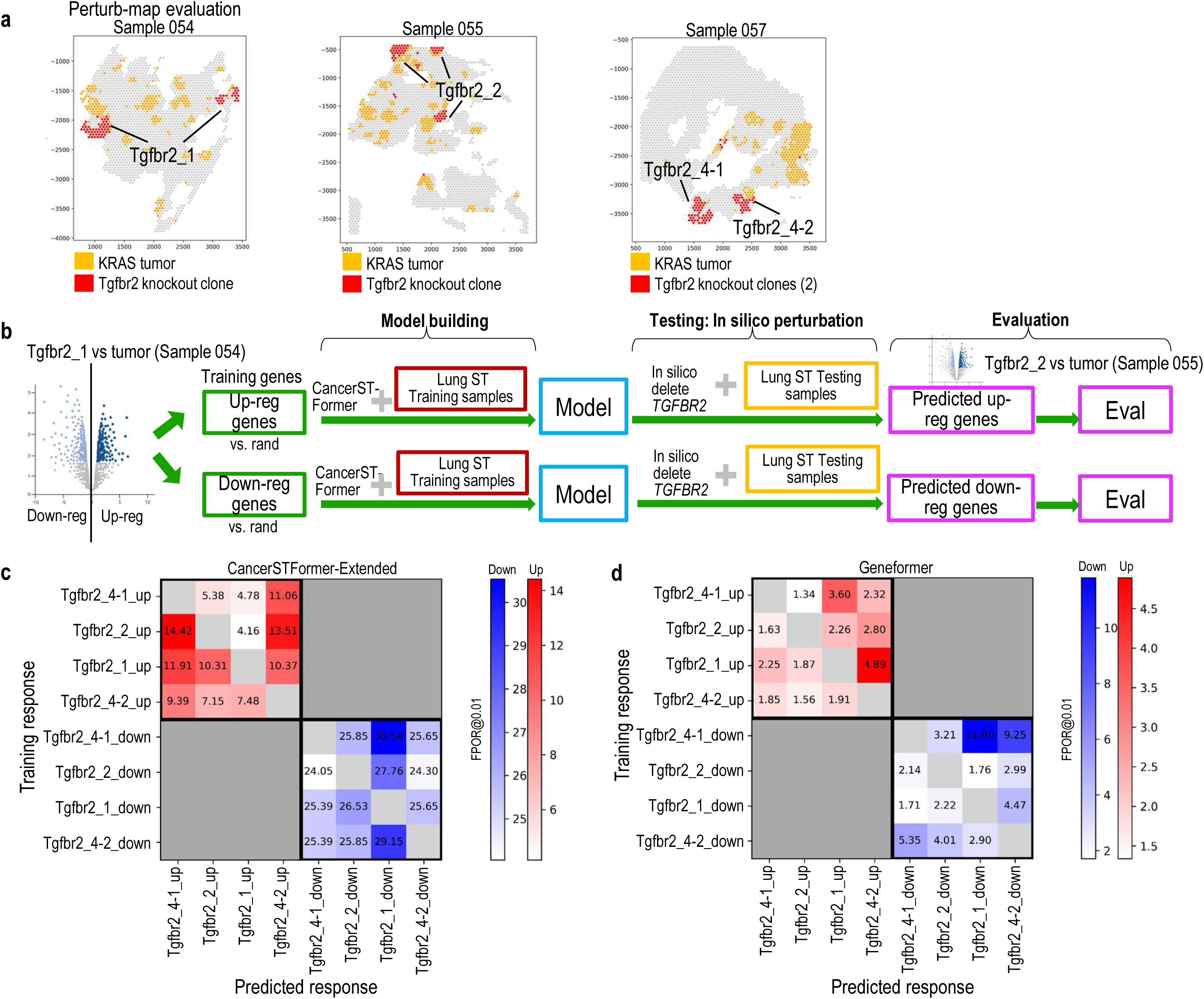
Spatial Perturb-map evaluation. **a.** Layout of 3 ST samples (054, 055, and 057) following Perturb-map experiments (GSE193460). These show the locations of 4 Tgfbr2 KO clones, and KRAS tumor clones (unperturbed). **b.** Overview of finetuning and testing procedure. For each KO clone, we derived up- and down-regulated genes, then used them to train a gene classifier within CancerSTFormer-Extended model with use of its available human lung ST samples. Then once the model is finetuned, during testing, given tumor (unperturbed) clones, we performed in silico gene perturbation of Tgfbr2 gene, using the trained model, to derive responsive genes. These predicted responsive genes were compared to actual response exhibited by Tgfbr2-KO clones. **c.** Performance results from cross-clone generalization evaluation (CancerSTFormer). To read this, for example, 5.38 is the performance (FPOR) from training on clone Tgfbr2_4-1_up, and predicting on clone Tgfbr2_2_up. **d.** Performance results for Geneformer.

### Spatial visualization of perturbation responses

We further provide visualization utilities to enable researchers to visualize the spatial response profiles following a perturbation. For example, given a CancerSTFormer model fine-tuned to discover ganitumab-sensitivity, we can perform *in silico* IGF1R deletion on a series of TNBC ST profiles to prioritize ST samples that would exhibit ganitumab (targeting IGF1R) sensitivity (**Figure 7a**). We can similarly prioritize ST samples based on resistance using a model trained to discover ganitumab-resistance (**Figure 7e**). In this case, we find that sample 1142243F preferentially exhibits sensitivity to ganitumab (**Figure 7a**), while samples GSM6433601-2 tend towards resistance (**Figure 7e**), illustrated by plotting resistance scores or sensitivity scores spatially. Response profiles can be pooled together across all IGF1R-expressed spots, then further clustered by UMAP (**Figure 7b**), or visualized with patient labels (**Figure 7c**). Finally, top markers defining ganitumab sensitivity or resistance can be obtained via differential analysis across response clusters (**Figure 7d, h**). These functions are readily accessible within the CancerSTFormer toolkit. When response predictions are aggregated over all ST profiles, they produce a master gene ranking of sensitivity/resistance that allows for separate evaluation on a hold-out cohort. The results so far pertain to the CancerSTFormer-50µm Local model but we also note similar prioritization of samples in the 250µm Extended model (**Supplementary Fig 4**). Overall, CancerSTFormer gives a per-ST perturbation prediction and enables a cohort-level evaluation of response signature genes.

**Figure 7:**
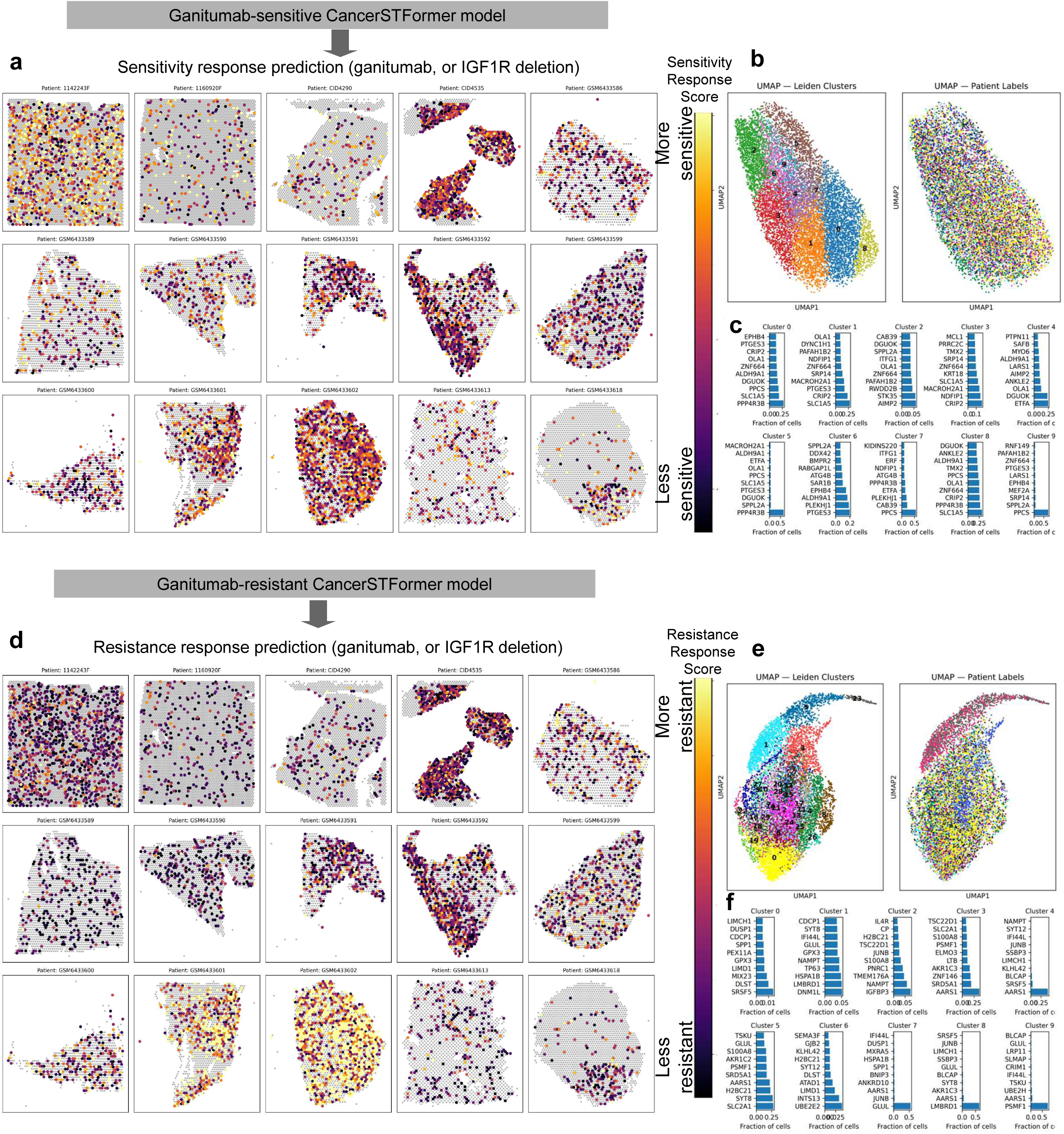
Spatial visualization of perturbation responses to ganitumab treatment. **a-d**. Given a fine-tuned ganitumab sensitive CancerSTFormer 50µm-Local model, TNBC ST samples were subject to IGF1R (which ganitumab targets) *in silico* deletion, and are ranked according to the overall sensitivity score (**a**). In each ST plot, non-gray spots are IGF1R-expressed, and color-gradient indicates degree of sensitivity (**a**). To derive the score (see Methods), gene shift was first computed for each gene in each spot forming a response profile, and then aggregated across the spots to derive master 100 sensitive genes. Per-spot enrichment of sensitive signature was then computed and plotted. UMAP plots show how spots cluster according to sensitivity response profiles (**b-c**). Panel **d** shows the most differential sensitivity markers of each spot cluster. **e-h.** Same as **a-d** but for ganitumab resistance signature.

### Finetuned CancerSTFormer enables the retrieval of organ-specific metastasis-associated genes, and responder/non-responder spot prediction to nivolumab therapy in hepatocellular cancer

To showcase the ability of CancerSTFormer to identify genes associated with organ-specific metastasis, we curated a set of experimentally validated genes shown in prior publications to be promoting/inhibiting/causing breast cancer metastasis to lung, bone, and brain (**Supplementary Data 3**). To prevent bias, none of these curated gene lists contained high-throughput RNAseq derived differentially expressed genes, which are correlative and unproven. Using a cross-validation set up, we next proceeded with evaluating the ability of CancerSTFormer to train and predict metastasis-associated genes based on the entire compendium of cancer ST data. Results have shown that our 50µm Local model can achieve AUCs of 0.78, 0.74, 0.72, for predicting genes associated with lung, bone, and brain metastasis respectively (**Figure 8a**). Furthermore, the 250µm Extended model considerably improves upon these AUCs to 0.85, 0.89, and 0.75, owing largely to the advantage of providing a larger spatial context (**Figure 8a**).

**Figure 8:**
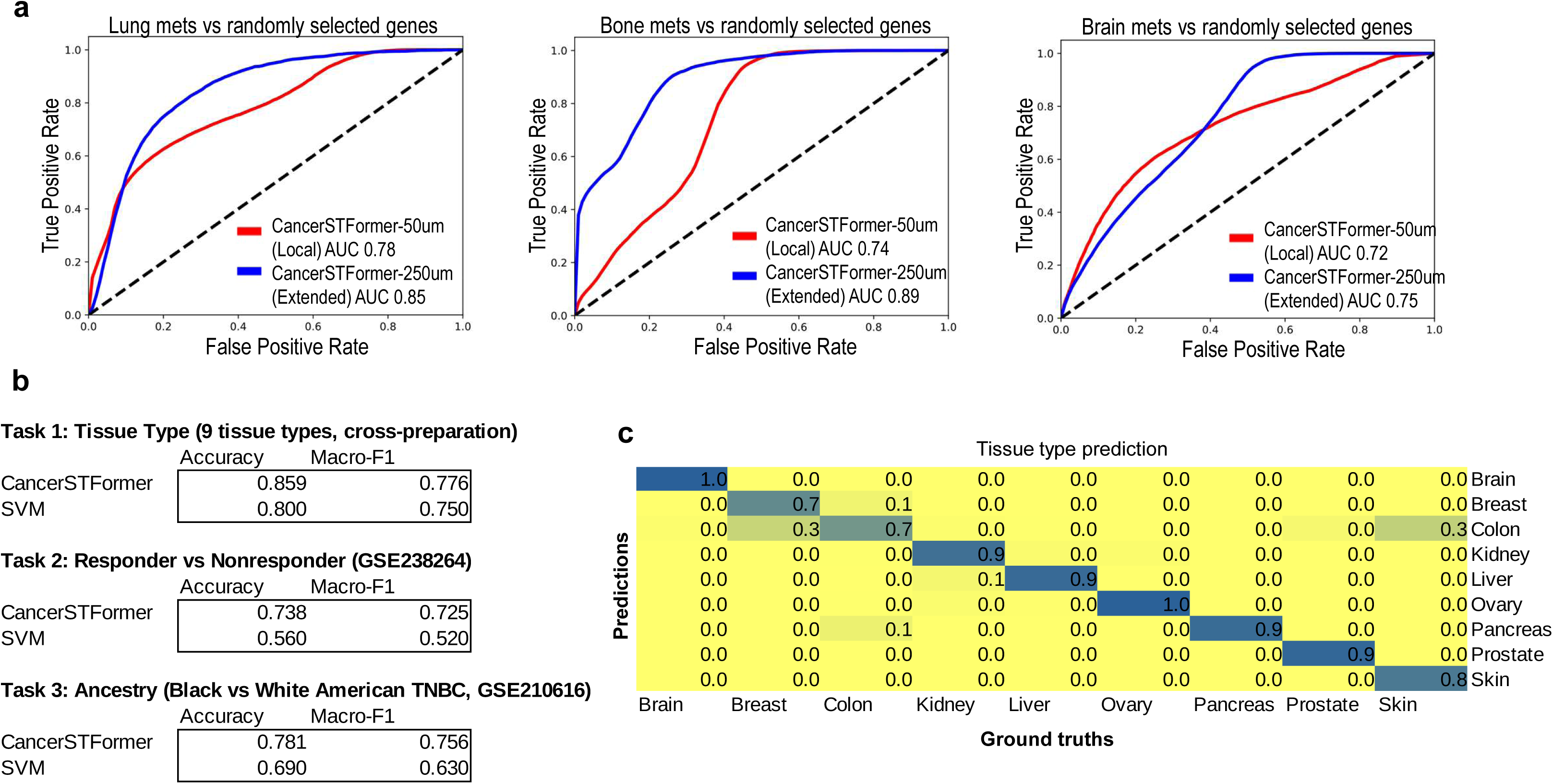
Prediction of metastasis-associated genes, tissue type, responder status, and ancestry from ST data. **a.** Prediction of breast cancer organ-specific metastasis associated genes. Manually curated, experimentally verified, literature supported metastasis-associated genes (∼60 or so) were made for each site of metastasis, defining the ground truths. These genes were split according to a 2-fold cross validation scheme to evaluate the ability of CancerSTFormer to predict such genes from the entire collection of ST data. A binary classifier was built to distinguish between each organ’s metastasis genes and randomly selected genes. Performances measured by AUROC on the holdout gene set, are indicated. **b.** Spot-based prediction of tissue-type, responder status (upon treatment with cabozantinib and nivolumab in HCC), and ancestry information. Patient-wide or sample-wide holdout was used. Accuracy and macro F1 scores were recorded. **c.** Details of the tissue-type prediction. Confusion matrix showing comparison between ground truths and prediction labels. Ochiai coefficients, as shown, measure the degree of overlap.

We finally benchmarked CancerSTFormer on a variety of spot-prediction tasks. In terms of cancer type prediction, the finetuned CancerSTFormer achieves an accuracy of 0.859 (macro F1 of 0.776) overall, which is superior to support-vector machine (SVM) (accuracy of 0.800 and macro F1 of 0.75) (**Figure 8b**). In terms of predicting responder and nonresponder status to nivolumab therapy in hepatocellular carcinoma patients^6^, CancerSTFormer has an accuracy of 0.738 in holdout patient samples, compared to 0.560 in SVM-baseline model, which is an improvement of 31% (**Figure 8b**). Finally, in ancestry prediction based on ST profiles^45^, CancerSTFormer is 13% better than SVM in accuracy, and 20% better in macro F1 (**Figure 8b**). It simultaneously classifies the majority of 9 cancer types correctly (see confusion matrix in **Figure 8c**). Along with the metastasis-gene prediction, these examples showcase the applicability of the foundation model across diverse downstream tasks.

## Discussion

In this study, we introduce CancerSTFormer, a spatially aware foundation model trained on 500 sequencing-based cancer spatial transcriptomic (ST) samples spanning diverse tumor types. By explicitly modeling neighborhood gene expression, CancerSTFormer addresses a major limitation of single-cell RNA-seq foundation models—which lack spatial context—and overcomes the constraints of Nicheformer, which excludes transcriptome-wide, spot-resolution ST. Supporting transcriptome-wide ST unlocks new applications, most notably *in silico* gene perturbation directly on spatial data, as demonstrated here.

Leveraging spot-resolution ST for foundation modeling is a deliberate strategic choice. Visium and related platforms represent the largest, most diverse, and truly transcriptome-wide ST resource available. Although imaging-based ST (Xenium^50^, MERSCOPE^51^, CosMx^52^) offers higher resolution, these platforms measure limited gene panels, are costly, and exist in far smaller numbers for large-scale modeling. We show that the reduced resolution of spot-based ST can be compensated—and often surpassed—by meta-analysis across large numbers of spot-resolution datasets, enabling accurate ligand–target retrieval. CancerSTFormer outperforms Geneformer and a few of single-cell ST (Visium HD, Xenium-5K) in ligand–target inference, demonstrating that modeling across many spatial niches compensates for lower resolution by offering greater diversity and statistical power, highlighting the importance of leveraging large models. This result encourages broad reuse of spot-resolution ST for modeling therapy response.

A central innovation of CancerSTFormer is its dual-scale spatial architecture (50μm Local and 250μm Extended), which captures short-range, contact-dependent interactions and long-range paracrine and stromal effects. We show that incorporating spatial context, particularly at the 250μm scale, substantially improves recovery of known ligand–target relationships and enhances prediction of in situ perturbation outcomes.

By recognizing spots as the fundamental modeling unit, CancerSTFormer naturally emphasizes the emerging concept of spatial niches that are echoed in other works: multicellular environments that drive tumor phenotypes more strongly than any single cells or single cell types^53–55^. Niche-level perturbations enable the discovery of immunotherapy targets that modulate the whole niche rather than any individual cell types. This differs from previous scRNAseq based *in silico* perturbation models, which investigate the perturbation effects on a single cell type, yet ignoring the contribution on the whole niche. Accordingly, we do not deconvolute spots into rigid cell-type proportions, an oversimplification requiring scRNAseq references that are not always available, but instead retain spot gene expression in full for pretraining the model.

Recent spatial foundation models highlight emerging interest. Single-cell spatial proteomic^56^ models link small marker panels to patient attributes; scGPT-spatial^57^ performs imputation but not spatial perturbations; stFormer^58^ uses spot-resolution but performs deconvolution as part of modeling which vastly simplifies the niche pattern and restricts cell-to-cell communication to ligand–receptor pairs. CancerSTFormer imposes neither restriction, and to our knowledge is the first full-transcriptome, sequencing-based spatial foundation model for human cancer with demonstrated perturbation capability.

We further show that CancerSTFormer can be fine-tuned on treatment resistance or sensitivity gene-sets derived from bulk studies. Fine-tuning prior to *in silico* gene perturbations enables the model to generalize resistant/sensitive signatures across unseen patients, without requiring paired ST and bulk datasets. This illustrates how spatially learned gene-regulatory structure in CancerSTFormer can enhance the transferability of predictive biomarkers across studies—a key challenge in bulk-only biomarker studies. This message is also consistent with prior work^30^.

Evaluating foundation models remains an active area of research^59–63^. Previous works in scRNAseq have shown the importance of comparing LLMs to a linear regression or correlation-based baseline method, for *in silico* perturbation studies^59,60^. Following best practices from these, we implemented a linear-baseline (Aggregated DE) and found CancerSTFormer substantially outperforms it. We adopted ranking-based metrics^63^ (precision-recall, fold-precision-over-random), to more specifically highlight the top predictions. We found that users usually need not examine beyond the top 10-20 genes in perturbation tasks. This small list limits biologist’s focus to the ones most likely to be meaningful. Other works have also used rank-based metrics for evaluation^63^, and similarly noted the accuracy of foundation models at top rank positions^59^.

At present, spatial Perturb-seq datasets remain very limited. Existing technologies have demonstrated pooled genetic screens in mouse liver and brain^64,65^, and emerging approaches such as spatial Perturb-Dbit^66^ are promising, but tumor-focused datasets are not widely available. We used a Perturb-map example to show how CancerSTFormer can be easily adaptable to training and predicting from spatial perturbation datasets. As spatial perturbation datasets come on the scene in future, these datasets will be highly valuable for fine-tuning CancerSTFormer. Our results indicate that while fine-tuning generally boosts performance – not all architectures benefit from it equally. The 250μm Extended model shows the greatest ability to predict resistant/sensitive gene-sets following fine-tuning. Despite its large pretraining corpus (10M), Geneformer, which lacks spatial modeling, in the fine-tuned case cannot match even our 50μm model, underscoring that spatial LLM architecture, not corpus size, is essential for representing niches. Simply feeding spatial labels or gene sets to a pretrained scRNAseq LLM during transfer learning step, in hopes of learning spatial niches, proved insufficient.

Overall, this work supports the view that local microenvironments are biologically meaningful units and that modeling them improves inference of ligand–target relationships and therapy response. CancerSTFormer’s *in silico* perturbation framework provides a powerful strategy for simulating therapeutic intervention and prioritizing immunotherapy targets. Perturbations of *PDCD1*, *CD274*, and *CTLA4* recapitulated known immunosuppressive programs, revealing contact-dependent suppression (captured by 50μm) and stromal/CAF-driven suppression (captured by 250μm), suggesting that disrupting immune evasion may require scale-specific therapeutic strategies.

In sum, CancerSTFormer distinguishes itself from existing spatial AI tools that identify spatial domains^67–70^ or other applications^32,71,72^ by its unique focus on modeling perturbations and predicting patient-level responses. Our results demonstrate that fine-tuning CancerSTFormer with targeted therapy response signatures enables robust generalization across studies. We believe that CancerSTFormer and models that leverage large amounts of spatial OMICS data through meta-analysis^47,73,74^ and representation learning are poised to become powerful tools for basic discovery and translational oncology.

## Methods

### Spatial transcriptomic data collection for pre-training CancerSTFormer

We collected over 511 publicly available spot-resolution, sequencing-based, whole-transcriptome spatial transcriptomic samples related to human cancer. In total, we collected 1.2 million spots from malignant tissues from 511 spatial transcriptomic samples across 50 studies. Transcriptomes and associated metadata were collected from Gene Expression Omnibus^75^ (GEO), Zenodo (https://zenodo.org/), ArrayExpress^76^, Datadryad (https://datadryad.org/), Human Tumor Atlas Network^2^, 10X Genomics, and Mendeley Data (https://data.mendeley.com/). Datasets’ original study information, associated publications, and metadata such as number of patients, samples, tissue type, technology, and other information were collected (**Supplementary Data 1**). For uniform processing and storage, we utilize Giotto^77^, Scanpy^78^, Numpy^79^, and Anndata^80^ with default settings. To keep high quality data, we removed spots that were detected as “out of tissue” and spots with no UMIs^81^ detected across all genes. We removed genes that had zero counts across cells in the dataset.

In building the foundational model, we required Ensembl gene IDs and number of counts stored. We used utilities like MyGene, org.Hs.eg.db^82^, and Ensembl^83^ to automatically convert gene symbols to Ensembl IDs. For samples that have raw gene counts, but no tissue coordinate file, we looked up the tissue coordinates based on the spot-barcodes, as we found that barcode-to-coordinates mapping to be conserved across studies. Sample level metadata such as cancer type, tissue preparation method, study ID, are recorded, and are stored into a separate database. These metadata are not used during the pre-training process, but rather they are used for evaluation and visualization purposes.

### Tokenization procedure and introduction of the 50µm Local and 250µm Extended scale models

The tokenization procedure builds upon the idea of rank-based gene encodings^27^, but also adds custom procedure for spatial data, including a “linearization” procedure to capture spatial neighborhoods in 1D sequence, and increased 2X “max length” to accommodate storage of both self- and neighbor-rank encoding information.

Before tokenization, we normalized gene counts within each spot using Giotto^77^ to enable computation of absolute gene medians and rankings. During tokenization, normalized expressions were stored as Tdigest^84,85^, which efficiently provide quantile medians for each gene across the full ST compendium, yielding global gene medians. To convert expression values for BERT training, we then ranked each spot’s genes relative to these global medians. For each spot, we identified the top-k genes (by median-normalized expression), returned their token IDs in descending order, and truncated to a user-defined “*max-length*”. This rank-based encoding is robust to variation in expression distributions across studies.

We describe two variants of CancerSTFormer, the 50µm Local and the 250µm Extended models, which differ by tokenization. In the 50µm Local mode, we performed gene ranking and tokenization through the process described above. In 250µm Extended mode, we calculated the neighborhood matrix through Delauney triangulation with SciPy^86^ and averaged gene rankings across all neighboring spots for each spot, which revealed the neighbor level tokenization. We then truncated this sequence to “*max-length*” and appended the sequence to the central spot’s token-sequence for a sequence of 2X “*max length*”. Such a spatial tokenization scheme, based on concatenated representations, echoes the approach used in DeepSpot^39^. This process effectively captures a neighborhood niche expression for each spot and gives context to gene expression. We utilized Hugging Face Tokenizer^87^ to store gene to ID vocabulary using a simple word level model. Finally, we saved ranked token IDs, neighbor token IDs, metadata, input IDs, and sequence length as a Hugging Face Dataset^87^ for pre-training. The 50µm Local model used the *max-length* of 2048, while the 250µm Extended model used 2X “*max length*” or 4096.

### Spatial masking during the 50µm Local and 250µm Extended model pretraining

Our pretraining class implements a masked language modeling^88^ (MLM) objective to pretrain a BERT-style transformer^89^. In the 50µm Local model pretraining, 15% of tokens are randomly masked, and the model learns to predict the masked genes based upon the unmasked context. This focuses the model on capturing gene-gene relationships and enables the model to build context-aware and biologically meaningful representations of gene activity. In the 250µm Extended case, due to the doubling of the allowed tokens accepted per sequence, equal number of masked tokens are selected from the center node and neighbor node portions of the token sequence. Masking in this Extended case thus permits center-to-neighbor and neighbor-to-center cross-neighbor spatial learning. Because each gene token is present at up to two times in the sequence in the 250µm Extended model, this model is capable of leveraging enhanced spatial context from longer max-length and repeated token presence. The masked pretraining optimizes the loss function:

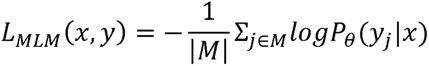

Where *M* is the set of masked tokens, *y* is the true token, and *x* is the observed tokens. Pretraining was performed using the Hugging Face Transformers library^87^, with masked language modeling implemented through PyTorch^90^. Datasets were prepared and loaded using Hugging Face Datasets^87^, and have the optional parameter for acceleration using DeepSpeed^91^.

### Model architecture during pretraining

CancerSTFormer uses six transformer units^92^, each with a self-attention layer and feed forward network layer. Additionally, each layer utilizes four attention heads which learns gene relationships in an unsupervised manner for downstream predictions. CancerSTFormer utilizes the following parameters to build models; max input 2048, relu activation, hidden dropout 0.02, attention dropout 0.02, learning rate 1e-3, linear scheduler, adamW optimizer, 10000 warmup steps, 3 epochs, weight decay 0.001, embedding dimensions 256, and initializer stander deviation of 0.02. Model architecture and parameters were preset based upon work done on single cell transformer models^27,92^. Model initialization was performed using Hugging Face Transformers library^87^.

### Adjustment of *in silico* gene perturbation for spatial models

The rank of genes is altered to model biological manipulations such as gene deletion.

#### The 50µm Local model

*In silico* deletion is performed by removing the token ID from the ranked gene encoding. We compute the cosine similarity between the original and perturbed gene embedding to predict the change on other gene token rankings from deletion of that gene. Specifically, the change in individual gene embeddings of the other genes, due to deletion of a particular gene, is computed as follows:

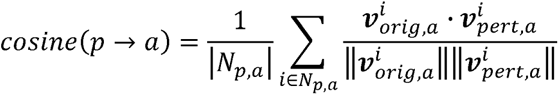

Where *p* is the perturbed gene, *a* is the affected gene, *N_p,a_* is the set of spots where *p* and *a* are expressed, *i* is the spot ID, 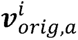 is the embedding vector for gene *a* in spot *i* before perturbation, and 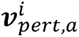 is the embedding vector for gene *a* in spot *i* after perturbation. Thus, *cosine*(*p* → *a*) captures the gene shift of the affected gene *a* from perturbing *p*. The 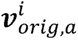 refers to the gene *a*’s embedding vector in the embedding matrix 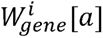 where 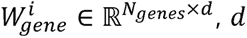 is the embedding dimension and *N_genes_* is the number of genes. Contextual embedding for each gene is created from the last-layer hidden states of the pretrained or finetuned neural network.

#### The 250µm Extended model

For *in silico* perturbation, we adjust the methodology because we have at most two tokens of the same gene in the rank encodings (i.e., one from the center spot and one from the averaged neighbor), making it possible to do **partial** (center or neighbor node) or **full** (both center and neighbor nodes) gene perturbations. We distinguish these two cases as the responses may be different.

#### Full perturbation

We first determine spots to perturb by collecting spots where the token to perturb is present in both spot and neighbor rank encodings. We therefore will delete two positions from the rank encodings and compute cosine similarity in the same way. This helps us determine the effect of simultaneous deletion of the gene in both center spot and neighboring spot to simulate the full effects of perturbation on spatial environments. After perturbation, the token sequence will be two tokens less across all examples. We used the *full perturbation* mode for ligand-targets, and PD1/IGF1R/ANG1 resistance/sensitivity predictions.

#### Partial perturbation

This covers the cases where the token to perturb is present in either the center node’s or averaged neighbor node’s token sequence, but not both. Token is deleted from each example, resulting in a new token sequence that is exactly 1 token less than the original state. The location of the “token to perturb”, i.e., whether in the center or neighbor portion, also matters and is distinguished by the model. This is because the Extended model uses separate positional encodings for center tokens [0…2047] and neighbor tokens [2048…4096] – the model can distinguish where the perturbed gene token originates. Removing a center token alters attention on all neighbor tokens as well as center tokens. Removing a neighbor token alters attention on all center tokens, and also other neighbor tokens. Thus, due to full self-attention, perturbing a token on either side ultimately influences both center and neighbor token’s attention. But the strongest perturbation effect is localized to the segment where the deletion occurs. Overall, both our *full perturbation* and *partial perturbation* modes for the 250µm Extended model capture **propagation effect of a gene perturbation** within a 250µm spatial niche.

#### Deriving the final affected master gene ranking

The above procedure computes a gene-shift (cosine similarity) per gene per spot. Genes with a lower cosine similarity are more likely to be affected by perturbation. For each gene, we further compute its average gene-shift among all spots (*AvgGeneShift*) and count the number of detected spots for the gene (*N. detections*). The final ranking is determined by the following pseudocode.

**Input:** AvgGeneShift: a rank-ordered list of tuples (gene, cosineSimValue); Ndetections: a dictionary mapping gene (key) to ndetect (value); detectMin: minimum cut off to count the gene

**Function** getRanking (AvgGeneShift, Ndetections, detectMin=-1):

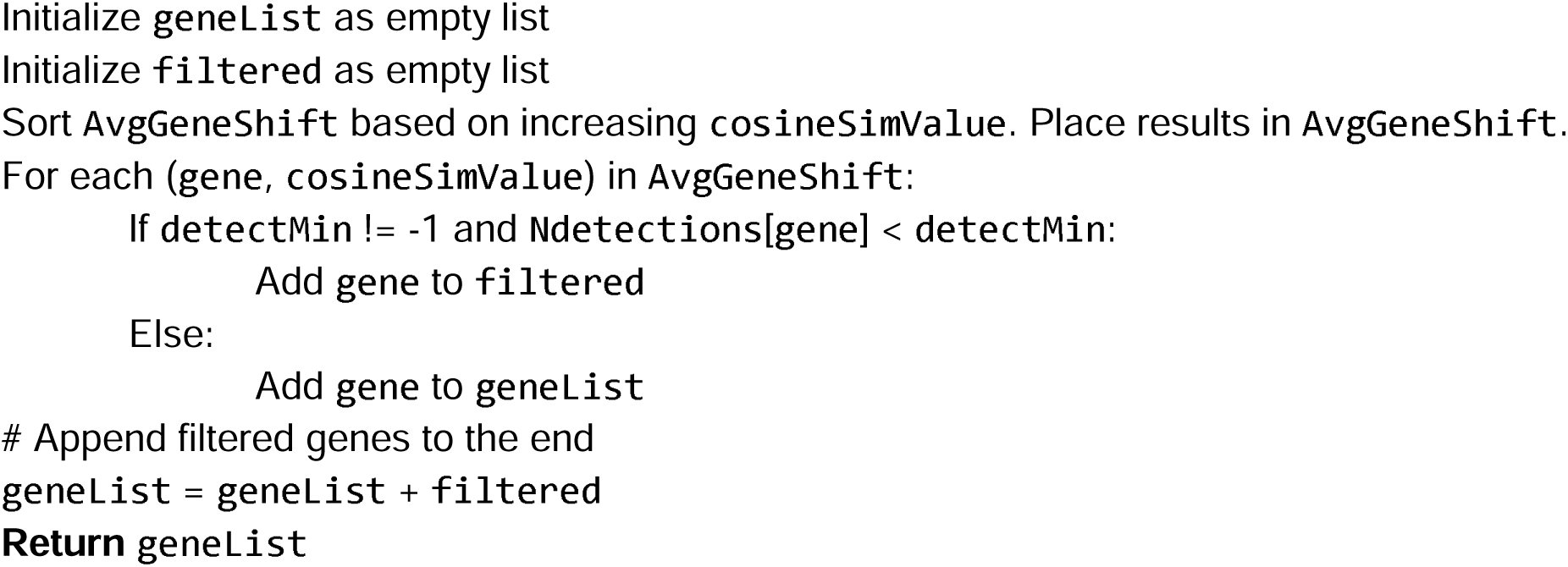

We set the detectMin to 100 in this paper, meaning 100 out of 1,000 max spots to perturb (10%) must contain the gene.

#### Choice of models for in silico perturbations

The pretrained CancerSTFormer models (50µm and 250µm variants), or finetuned models (for recognizing sensitivity and resistance genes) were used. Finetuning procedure is later described.

### Evaluation using NicheNet ligand-targets gene-sets

#### Gold standard construction

Ligand-target matrix was downloaded from the NicheNet^40^ Github page. Briefly, NicheNet compiled sets of ligand-target relationships for ∼1,200 ligands from experimental sources and integrative computational modeling^40^. These sources measure ligand-perturbation effects on target genes across a number of validation datasets and produce a weighted network, available as ligand-target matrix. For each ligand, top 500 target genes with highest weights in the matrix are retained, producing a target gene-set for each ligand. Likewise after repeating this procedure for all ligands, the complete ligand-target gene-sets are constructed and used as gold-standards during evaluation.

#### Methods compared

We perturbed each ligand using **CancerSTFormer** and **Geneformer**^27^ pretrained models and evaluated each system’s ability to retrieve ligand-targets in the affected gene list. In these cases, the perturbed gene is the ligand, and the perturbed datasets are either TNBC-related Visium ST datasets (for CancerSTFormer) (zenodo.4739739 and GSE210616), or single-cell RNAseq datasets (for Geneformer) (GSE176078). We also evaluated two retrieval methods made using single-cell spatial transcriptomic data, namely **Xenium-5K**^50,93,94^ and **Visium HD**^93,95^. Breast cancer related Xenium-5K and Visium HD datasets, each with over ∼300,000 single cells, were downloaded from the 10X Genomics website [selecting “Xenium-5K Human Breast Cancer with 5K Human Pan Tissue and Pathways Panel”, and “Visium HD Spatial Gene Expression Library, Human Breast Cancer (FFPE)]. To identify ligand-targets using Xenium-5K and Visium HD, we used a procedure described in our prior published work^53^, whereby given a ligand, we retrieve neighbors’ DE genes from comparing neighbors of ligand-high cells and neighbors of ligand-low cells in Xenium-5K and Visium HD datasets. These ligand-specific niche DE-genes, likely to be targets of that ligand, are evaluated.

#### Evaluation metric

Retrieval performance was expressed as fold precision-over-random, as we have used in our previous publication^47^. Here is the definition of fold precision-over-random (FPOR^47^) at each recall:

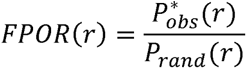

Where *r* is the recall level, 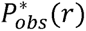 is the observed interpolated precision at recall *r*, *P_rand_* (*r*) is the expected precision from random guess. *P_rand_* (*r*) = *N_positive_*/*N_universe_* where *N_positive_* is the number of positive genes (i.e., targets), and *N_universe_* is the total genes in genome. *P_rand_* (*r*) is fixed at all recall levels. 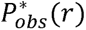 is the interpolated precision:

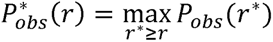

Where *P_obs_* is the precision at recall level *r.* 100-point precision-recall curve was constructed from *FPOR*(*r*) curve was constructed from each recall level among 0.01, 0.02, …, 0.99, 1.0.

### Evaluation using single-cell spatial ST datasets (Xenium-5K)

Separate from NicheNet^40^, we also evaluated CancerSTFormer and Geneformer using niche-DE genes determined from external single-cell spatial Xenium-5K^50^ ST dataset (10X Genomics, Human Breast Cancer with 5K Human Pan Tissue and Pathways Panel). This helps to assess the systems’ ability to retrieve ligand-induced genes that are spatially coexpressed within local spatial niches. We divided ligands into immune-checkpoint and stroma-epithelial categories - each category contained ligands previously reported^96^ to be important in TNBC. Similar to described previously^53^, we first constructed gold-standard ligand-targets by defining ligand niche-DE genes within Xenium-5K after comparing ligand-high and ligand-low niches^53^. CancerSTFormer and Geneformer’s perturbation results were next compared to this standard.

### Evaluation using literature guided search

Beyond large-scale evaluations using NicheNet^40^ and Xenium-5K^50^, we also evaluated the *PDCD1*, *CD274*, and *CTLA4 in silico* perturbation results using literature search. Top 10 perturbation-affected genes, sorted by having lowest cosine similarity to the perturbed ligand, were each assessed by looking for literature evidence of the affected gene linking to the immune-checkpoint blockade. If a publication describing such a link is discovered in some cancer types in mouse or human studies, it is included in the table of evidence. Evidence is further characterized by study type (immunotherapy biomarker study or mechanistic study), cell-type that it affects (if it is a mechanistic study), distance of regulation, and nature of regulation (boosts or suppresses immunity). Overall, literature guided search helps us to discover possible novel immunotherapy targets from the list of immunosuppressive genes that the CancerSTFormer model returns.

### Evaluating the prediction of treatment-resistant and sensitive genes made from *PDCD1*, *IGF1R*, and *ANGPT1 in silico* perturbations

To understand the biological processes associated with treatment resistance and sensitivity, we probed the most affected genes from *in silico* gene perturbation studies. We evaluated the model’s ability to retrieve treatment-resistant/sensitive genes given only the molecular targets (PD1, IGF1R, ANG1, represented by genes *PDCD1*, *IGF1R*, and *ANGPT1* respectively) and provided ST samples. Predictions were made along Unsupervised or Supervised pathway.

#### Gold-standard used for evaluation

The ISPY2 trial (GSE194040)^48,97^ enlisted TNBC patients who were treated with pembrolizumab (targeting PD1), ganitumab (targeting IGF1R), and trebanalib (targeting Angiopoietin-1 or *ANGPT1*). Pathological complete response data were provided for these patients. We downloaded the baseline (i.e., prior to treatment) RNAseq profiles of patient samples from this trial (GSE194040^48^). We computed PD1-sensitive genes and PD1-resistant genes by comparing the gene expression profiles of patient groups achieving pCR and with no response from treatment. Likewise, we computed sensitive and resistant gene-sets for anti-IGF1R and anti-ANG1 treatments.

#### Methods compared

Among **Unsupervised methods**, we have compared **CancerSTFormer pretrained models** and **Aggregated DE**.

- <u>CancerSTFormer pretrained models</u>: *PDCD1*, *IGF1R*, and *ANGPT1* were each *in silico* deleted in TNBC ST samples using the pretrained 50µm Local and 250µm Extended models. Resulting genes were compared against ISPY2 pembrolizumab, ganitumab, and trebanalib resistance/sensitivity gene lists.
- <u>Aggregated DE</u>: this is correlation baseline method. Here, given a gene to perturb (e.g. *PDCD1*), for each ST sample, we defined DE genes between *PDCD1*-high spots and *PDCD1*-low spots using t-test. High and low groups were defined by top 10% and bottom 10% of spots by expression. Then for each gene, we combined its P-value from DE test across ST samples using Stouffer’s z-score method, to generate a meta gene ranking. Let *Z_i_* = *ϕ*^-1^(*p_i_*), the meta-z score is 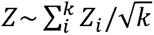 where *ϕ* is the standard normal cumulative distribution function, *p_i_* is the one-sided p-value, and *k* is the number of tests. Genes were sorted by descending *Z* and were evaluated.

Among **Supervised methods**, we have compared **CancerSTFormer finetuned models**, **GeneFormer finetuned model**, and **No Refinement control**.

- <u>CancerSTFormer finetuned models:</u> We first set up the data for evaluation. Briefly we divided the ISPY2 patients into Train and Test cohorts. The Train cohort was used to finetune the models by building a Gene Classifier that recognized resistant and sensitive genes defined on the Train cohort. The Test cohort was used for evaluating predictions of resistance/sensitivity genes. Details of the finetuning procedure are described in the next section. After model training, *PDCD1*, *IGF1R*, and *ANGPT1* were each *in silico* deleted in TNBC ST samples using the finetuned 50µm Local and 250µm Extended models.
- Geneformer finetuned model: Similar procedure to above, except that finetuning proceeded with TNBC single-cell RNAseq dataset.
- No Refinement control: resistance/sensitivity genes of Train cohort and those of the Test cohort were directly compared (without feeding them to any models).

#### Evaluation

These sensitive and resistant gene-sets were used as gold-standards to evaluate perturbation-affected genes derived from *in silico* gene perturbations of *PDCD1*, *IGF1R*, and *ANGPT1* genes, predicted by CancerSTFormer and applied to TNBC ST datasets. The detectMin parameter (in the pseudocode in previous section) is an important factor. We set detectMin to 100, maxSpotsToPerturb to 1000, for ligand-targets predictions (Figure 3), and *in silico* perturbation evaluations (Figure 5c-d). We set finetuning to optimize the objective of eval loss for all cases, except for *ANGPT1*-sensitivity and *PDCD1*-resistance which optimizes F1-scores, suggested by cross-validations.

### Fine-tuning the model for improved *in silico* gene perturbations

CancerSTFormer was fine-tuned by training the model to recognize the PD1-resistant/sensitive gene-sets. We first defined the resistant or sensitive gene sets as follows. For each treatment (PD1, IGF1R, or ANG1)^48^, we split the enrolled patients into training and test sets – within each set we made sure there are both pCR=1 and pCR=0 patients. Within the training set, the RNASeq profiles^48^ of patients were compared (based on response status) to derive PD1-responsive and resistant genes used for training the model. The test set was used only for evaluation.

During training, a gene classifier was built. We used a two-layer neural network to train this classifier on top of the existing pre-trained network, where the first 4 of 6 layers of the pretraining network are kept frozen and the next two layers are varied (i.e., the trainable layers). We used Ray (https://docs.ray.io) to assist in hyperparameter tuning. The best 5 models, discovered from 60 trials, were selected on the basis of having the lowest evaluation loss or highest macro F1 score based on cross-validations (see **Supplementary Data 2** for parameter setting). Average of the best 5 models were reported for perturbation tasks.

To evaluate the performance, the test patient set was used to determine similarly resistant/sensitive gene-set. This gene-set was treated as gold-standard to evaluate predictions. In this way, training and testing sets were independent.

#### Hyperparameter tuning during fine tuning

Training was managed through the Hugging Face Trainer^98^ interface. Key arguments included cosine learning rate scheduler, gradient accumulation, weight decay, and mixed precision to support large scale fine-tuning. We performed hyperparameter tuning through Ray Tune library^99^, using HyperOpt based search over model configuration, such as learning rate, epochs, weight decay, scheduler type, etc. This method evaluated important metrics across different hyperparameters and saves all checkpoint models as well as best models. Final models were saved in standardized Hugging Face format for reproducibility and downstream integration. One can optimize Ray Tuning based on evaluation metrics such as loss function, accuracy, macro F1-score, calculated on the training set.

In both the gene-classification and spot-level classification, the models sought to optimize the cross-entropy loss function, defined as:

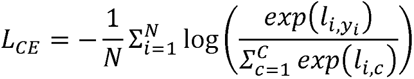

Where *N* is the number of training samples, *C* is the number of classes,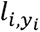 gives the logit (raw score) for sample *i* and true class label *y*, and *l_i,c_* gives the logit for sample *i* and class *c*. The logarithm gives the softmax probability that the model assigns to the true class.

### Perturb-map evaluations

We downloaded Perturb-map–associated 10x Visium spatial transcriptomics datasets (GSM5808054–GSM5808057) from the Gene Expression Omnibus. These data come from an in vivo CRISPR mouse lung cancer model. Spots were annotated by the original authors as Tgfbr2-knockout (KO) clones (Tgfbr2_1, Tgfbr2_2, Tgfbr2_4-1, and Tgfbr2_4-2), Kras tumor clones (KP clones; KP_1-1, KP_1-2, …), or normal epithelial cells. For each Tgfbr2 KO clone, we computed differentially up- and down-regulated gene signatures relative to the KP clones. We used these up- and down-genes to finetune CancerSTFormer, based on lung specific ST datasets in CancerSTFormer, to train its ability to predict Tgfbr2-KO responsive genes. We fine-tuned the 250 µm–Extended model because of its superior performance. We then performed a train-on-one-clone, test-on-another evaluation: for each Tgfbr2 KO clone, we trained on that clone and evaluated on each of the remaining KO clones. For testing, we performed in silico deletion of Tgfbr2 in KP (unperturbed) tumor spots within each sample and assessed whether the model could recapitulate the response genes observed in the corresponding Tgfbr2 KO clone from the same sample.

### Fine-tuning the model for organ-specific metastasis-associated gene predictions

#### Training gene-sets

We curated a short list of experimentally validated metastasis associated genes for breast cancer lung, brain, and bone metastases by using a literature guided approach. Specifically, we entered in Google a custom search query made of “breast cancer lung metastasis gene”, followed by a list of regulatory keywords such as “regulate inhibit promote activate initiate suppress”. This composite query is expected to return genes currently reported in literature to regulate a specific organ metastasis. Top 10 pages of Google search results were next reviewed by humans, filtered to those results that are referring to publication articles. For each hit, we clicked into the publication link, viewed the abstract and article contents, and checked to ensure that the article experimentally validates the gene’s role in that specific metastasis in animal models or human cells. Any genes derived from high-throughput RNAseq studies without further experimental validation are removed. This produced a high-confidence list of 60-70 metastasis-associated genes per organ (**Supplementary Data 3**).

#### Finetuning procedure and evaluation

We next built a CancerSTFormer model to predict metastasis-associated genes per organ using a 2-fold cross-validation set up. Input dataset used was the entire ST compendium. For each organ site of metastasis, training gene-set consisted of 50% of metastasis-associated genes of that organ site (positive class), and random genes of matched size selected from genome (negative class).

CancerSTFormer then trained a 2-layer neural network, on top of the existing pretrained model, to recognize metastasis-associated genes using ST data. After the model was learnt, predictions were next made on test gene-set. Hyperparameters are tuned by the Ray tuning library. To evaluate the predictions, we constructed a sensitivity vs (1-specificity) curve and computed the AUROC.

### Fine-tuning the model for spot-level predictions (i.e., cancer type, responder status, and ancestry)

GSE210616^45^ provided patient-level ancestry information and treatment status (treated vs untreated). GSE238264^6^ provided responder and nonresponder status for 4 patients with hepatocellular carcinoma. ST data were downloaded from both studies using Gene Expression Omnibus. We adopted patient-wise cross-validation scheme, where 1 patient of each group was held out for evaluation purpose, and N-2 patients were used for finetuning the CancerSTFormer model (from pretrained model). For cancer type prediction, we created a mini, balanced, multi-tissue ST dataset with 2-4 studies per tissue. We then trained the model using the Spot Classifier mode. Hyperparameters are tuned by the Ray tuning library. To evaluate the predictions, we constructed a sensitivity vs (1-specificity) curve and computed the AUROC.

### Spatial visualization of gene deletion responses

#### Computation of sensitivity/resistance score per spot

The finetuned model was used to perform *in silico* gene deletion simulating IGF1R, PDCD1, and ANGPT1 inhibition. We first obtained the top 100 master response genes, *G*_100_, from such perturbations (see previous section *“Deriving the final affected master gene ranking”*). Then for each spot, we sorted all genes based on gene-shift and converted them into gene ranks (1…|*G*| where *G* is the number of genes in genome). Then the response score is computed as:

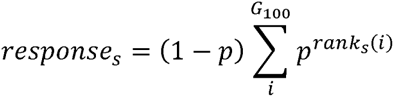

where *s* is the spot, *p* is a rank-weighting constant (inspired by rank-biased precision^100^, and was set to 0.99), and *rank_s_*(i) is the rank of gene*i*in spot *s*. The response score was visualized over spots’ spatial coordinates in ST samples.

#### Clustering of spots by response profiles

Let *M*(*s*, *j*) be the spot-by-gene response matrix, where *s* is spot, and *j* is top 100 genes (*G*_100_). Each element (*s*, *j*) of *M* is 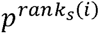 where *p* set to 0.99. We clustered *M* using Leiden clustering and visualized spot clusters using UMAP plot in Scanpy. Top 5 marker genes in each cluster were obtained and visualized using horizontal bar plots.

## Data Availability

We provide pre-trained and fine-tuned models generated on our large corpus described in the dataset section of methods and available on our hugging face model hub. The code, installation, and usage for CancersTFormer is made publicly available on Github at (https://github.com/bernard2012/CancerST). Models and datasets are available on Zenodo (https://zenodo.org/records/17807001). A website contains supplementary codes, tutorials, and visualization (https://qianzhulab.github.io/suppl/CancerST/). All data used for our modeling and analysis are publicly available. Datasets utilized to train our Local (50µm) and Extended (250µm) models can be found as part of Supplementary Data, where study names, dataset links, and metadata are provided.

## Acknowledgements

This work was supported by Cancer Prevention and Research Institute of Texas (CPRIT) grant RR220035, Marion R Wright Award from BCRF, and internal funding from Baylor College of Medicine. QZ is a CPRIT Scholar.

## Supplementary Material List

**Supplementary Table 1**: *In silico* gene perturbations (*CD274*, *CTLA4*, and *PDCD1* deletion) with the CancerSTFormer-50µm Local model

**Supplementary Table 2**: *In silico* gene perturbations (*CD274*, *CTLA4*, and *PDCD1* deletion) with the CancerSTFormer-250µm Extended model

**Supplementary Figure 1**: Evaluation of the pretrained embedding’s cluster coherence by measuring intra-cluster vs inter-cluster cosine similarity.

**Supplementary Figure 2**: Removal of sample preparation related batch effects on the pretrained embedding

**Supplementary Figure 3**: Comparison with a baseline method, Aggregated DE.

**Supplementary Figure 4:** Spatial visualization of ganitumab treatment responses using CancerSTFormer 250µm Extended model.

**Supplementary Data 1**: Spatial transcriptomic dataset summary

**Supplementary Data 2**: CancerSTFormer parameter tuning for various fine-tuned tasks

**Supplementary Data 3**: High-confidence list of breast cancer metastasis-associated genes per organ

## References

1. Hanahan, D. & Weinberg, R. A. Hallmarks of Cancer: The Next Generation. Cell 144, 646–674 (2011).

2. Rozenblatt-Rosen, O. et al. The Human Tumor Atlas Network: Charting Tumor Transitions across Space and Time at Single-Cell Resolution. Cell 181, 236–249 (2020).

3. Barkley, D. et al. Cancer cell states recur across tumor types and form specific interactions with the tumor microenvironment. Nature Genetics 2022 54:8 54, 1192–1201 (2022).

4. Khaliq, A. M. et al. Spatial transcriptomic analysis of primary and metastatic pancreatic cancers highlights tumor microenvironmental heterogeneity. Nat. Genet. 56, 2455–2465 (2024).

5. Klughammer, J. et al. A multi-modal single-cell and spatial expression map of metastatic breast cancer biopsies across clinicopathological features. Nat. Med. 30, 3236–3249 (2024).

6. Zhang, S. et al. Spatial transcriptomics analysis of neoadjuvant cabozantinib and nivolumab in advanced hepatocellular carcinoma identifies independent mechanisms of resistance and recurrence. Genome Med. 15, 72 (2023).

7. Arora, R. et al. Spatial transcriptomics reveals distinct and conserved tumor core and edge architectures that predict survival and targeted therapy response. Nat. Commun. 14, 5029 (2023).

8. Ståhl, P. L. et al. Visualization and analysis of gene expression in tissue sections by spatial transcriptomics. Science (1979). 353, 78–82 (2016).

9. Rodriques, S. G. et al. Slide-seq: A scalable technology for measuring genome-wide expression at high spatial resolution. Science (1979). 363, 1463–1467 (2019).

10. Liu, Y. et al. High-Spatial-Resolution Multi-Omics Sequencing via Deterministic Barcoding in Tissue. Cell 183, 1665–1681.e18 (2020).

11. Eng, C. H. L. et al. Transcriptome-scale super-resolved imaging in tissues by RNA seqFISH+. Nature 2019 568:7751 568, 235–239 (2019).

12. Xia, C., Fan, J., Emanuel, G., Hao, J. & Zhuang, X. Spatial transcriptome profiling by MERFISH reveals subcellular RNA compartmentalization and cell cycle-dependent gene expression. Proc. Natl. Acad. Sci. U. S. A. 116, 19490–19499 (2019).

13. Wang, X. et al. Three-dimensional intact-tissue sequencing of single-cell transcriptional states. Science 361, (2018).

14. Stickels, R. R. et al. Highly sensitive spatial transcriptomics at near-cellular resolution with Slide-seqV2. Nat. Biotechnol. 39, 313–319 (2021).

15. Chen, A. et al. Spatiotemporal transcriptomic atlas of mouse organogenesis using DNA nanoball-patterned arrays. Cell 185, 1777–1792.e21 (2022).

16. Vickovic, S. et al. High-definition spatial transcriptomics for in situ tissue profiling. Nat. Methods 16, 987–990 (2019).

17. Cho, C.-S. et al. Microscopic examination of spatial transcriptome using Seq-Scope. Cell 184, 3559–3572.e22 (2021).

18. Schott, M. et al. Open-ST: High-resolution spatial transcriptomics in 3D. Cell 187, 3953–3972.e26 (2024).

19. Bommasani, R., et al. On the Opportunities and Risks of Foundation Models. (2022).

20. Vaswani, A., et al. Attention Is All You Need. (2023).

21. Radford, A. et al. Language Models are Unsupervised Multitask Learners. in (2019).

22. Brown, T. B. et al. Language Models are Few-Shot Learners. (2020).

23. Devlin, J., Chang, M.-W., Lee, K. & Toutanova, K. BERT: Pre-training of Deep Bidirectional Transformers for Language Understanding. (2019).

24. Brin, S. & Page, L. The anatomy of a large-scale hypertextual Web search engine. Computer Networks and ISDN Systems 30, 107–117 (1998).

25. Krizhevsky, A., Sutskever, I. & Hinton, G. E. ImageNet classification with deep convolutional neural networks. Commun. ACM 60, 84–90 (2017).

26. Cui, H. et al. scGPT: toward building a foundation model for single-cell multi-omics using generative AI. Nat. Methods 21, 1470–1480 (2024).

27. Theodoris, C. V. et al. Transfer learning enables predictions in network biology. Nature 618, 616–624 (2023).

28. Zhang, Y., Chen, L., Wang, S. & others. scFoundation: Foundation models for single-cell biology. bioRxiv https://doi.org/10.1101/2023.05.29.542705 (2023) doi:10.1101/2023.05.29.542705.

29. Yang, F. et al. scBERT as a large-scale pretrained deep language model for cell type annotation of single-cell RNA-seq data. *Nat*. Mach. Intell. 4, 852–866 (2022).

30. Kalfon, J., Samaran, J., Peyré, G. & Cantini, L. scPRINT: pre-training on 50 million cells allows robust gene network predictions. Nat. Commun. 16, 3607 (2025).

31. Zeng, Y. et al. CellFM: a large-scale foundation model pre-trained on transcriptomics of 100 million human cells. Nat. Commun. 16, 4679 (2025).

32. Schaar, A. C., et al. Nicheformer: A Foundation Model for Single-Cell and Spatial Omics. Preprint at 10.2139/ssrn.4803291 (2024).

33. Lotfollahi, M., Wolf, F. A. & Theis, F. J. scGen predicts single-cell perturbation responses. Nat. Methods 16, 715–721 (2019).

34. Roohani, Y., Huang, K. & Leskovec, J. Predicting transcriptional outcomes of novel multigene perturbations with GEARS. Nat. Biotechnol. 42, 927–935 (2024).

35. Wang, Z. J. et al. Identifying perturbations that boost T-cell infiltration into tumours via counterfactual learning of their spatial proteomic profiles. *Nat*. Biomed. Eng. 9, 390–404 (2025).

36. Megas, S., et al. Celcomen: spatial causal disentanglement for single-cell and tissue perturbation modeling. (2024).

37. Cui, Y. & Yuan, Z. Prioritizing perturbation-responsive gene patterns using interpretable deep learning. Nat. Commun. 16, 6095 (2025).

38. Jin, S. et al. Inference and analysis of cell-cell communication using CellChat. Nature Communications 2021 12:1 12, 1–20 (2021).

39. Nonchev, K. et al. DeepSpot: Leveraging Spatial Context for Enhanced Spatial Transcriptomics Prediction from H&E Images. Preprint at 10.1101/2025.02.09.25321567 (2025).

40. Browaeys, R., Saelens, W. & Saeys, Y. NicheNet: modeling intercellular communication by linking ligands to target genes. Nature Methods 2019 17:2 17, 159–162 (2019).

41. André, T. et al. Nivolumab plus Ipilimumab in Microsatellite-Instability–High Metastatic Colorectal Cancer. New England Journal of Medicine 391, 2014–2026 (2024).

42. Rohaan, M. W. et al. Tumor-Infiltrating Lymphocyte Therapy or Ipilimumab in Advanced Melanoma. New England Journal of Medicine 387, 2113–2125 (2022).

43. Cortes, J. et al. Pembrolizumab plus Chemotherapy in Advanced Triple-Negative Breast Cancer. New England Journal of Medicine 387, 217–226 (2022).

44. Schmid, P. et al. Atezolizumab and Nab-Paclitaxel in Advanced Triple-Negative Breast Cancer. New England Journal of Medicine 379, 2108–2121 (2018).

45. Bassiouni, R. et al. Spatial Transcriptomic Analysis of a Diverse Patient Cohort Reveals a Conserved Architecture in Triple-Negative Breast Cancer. Cancer Res. 83, 34–48 (2023).

46. Wu, S. Z. et al. A single-cell and spatially resolved atlas of human breast cancers. Nat. Genet. 53, 1334–1347 (2021).

47. Zhu, Q. et al. Targeted exploration and analysis of large cross-platform human transcriptomic compendia. Nat. Methods 12, 211–214 (2015).

48. Wolf, D. M. et al. Redefining breast cancer subtypes to guide treatment prioritization and maximize response: Predictive biomarkers across 10 cancer therapies. Cancer Cell 40, 609–623.e6 (2022).

49. Dhainaut, M. et al. Spatial CRISPR genomics identifies regulators of the tumor microenvironment. Cell 185, 1223–1239.e20 (2022).

50. Janesick, A. et al. High resolution mapping of the tumor microenvironment using integrated single-cell, spatial and in situ analysis. Nat. Commun. 14, 8353 (2023).

51. Moffitt, J. R. et al. Molecular, spatial, and functional single-cell profiling of the hypothalamic preoptic region. Science (1979). 362, (2018).

52. He, S. et al. High-plex imaging of RNA and proteins at subcellular resolution in fixed tissue by spatial molecular imaging. Nature Biotechnology 2022 40:12 40, 1794–1806 (2022).

53. Zhu, Q. et al. Integrative spatial omics reveals distinct tumor-promoting multicellular niches and immunosuppressive mechanisms in Black American and White American patients with TNBC. Nat. Commun. 16, 6584 (2025).

54. Jackson, H. W. et al. The single-cell pathology landscape of breast cancer. Nature 2020 578:7796 578, 615–620 (2020).

55. Wang, X. Q. et al. Spatial predictors of immunotherapy response in triple-negative breast cancer. Nature 2023 621:7980 621, 868–876 (2023).

56. Wenckstern, J., et al. AI-powered virtual tissues from spatial proteomics for clinical diagnostics and biomedical discovery. (2025).

57. Wang, C. et al. scGPT-spatial: Continual Pretraining of Single-Cell Foundation Model for Spatial Transcriptomics. Preprint at 10.1101/2025.02.05.636714 (2025).

58. Cao, S. et al. stFormer: a foundation model for spatial transcriptomics. Preprint at 10.1101/2024.09.27.615337 (2024).

59. Li, C. et al. Benchmarking AI Models for *In Silico* Gene Perturbation of Cells. Preprint at 10.1101/2024.12.20.629581 (2024).

60. Ahlmann-Eltze, C., Huber, W. & Anders, S. Deep-learning-based gene perturbation effect prediction does not yet outperform simple linear baselines. Nat. Methods 22, 1657–1661 (2025).

61. Han, C. et al. Reusability report: Exploring the transferability of self-supervised learning models from single-cell to spatial transcriptomics. *Nat*. Mach. Intell. 7, 1414–1428 (2025).

62. Boylan, J. et al. Single Cell Foundation Models Evaluation (scFME) for In-Silico Perturbation. Preprint at 10.1101/2025.09.22.677811 (2025).

63. Liu, X. et al. Evaluating Foundation Models for *In-Silico* Perturbation. Preprint at 10.1101/2025.05.11.653338 (2025).

64. Binan, L. et al. Simultaneous CRISPR screening and spatial transcriptomics reveal intracellular, intercellular, and functional transcriptional circuits. Cell 188, 2141–2158.e18 (2025).

65. Saunders, R. A. et al. Perturb-Multimodal: A platform for pooled genetic screens with imaging and sequencing in intact mammalian tissue. Cell 188, 4790–4809.e22 (2025).

66. Fan, R., et al. Spatially Resolved Panoramic in vivo CRISPR Screen via Perturb-DBiT. Preprint at 10.21203/rs.3.rs-6481967/v1 (2025).

67. Hu, J. et al. SpaGCN: Integrating gene expression, spatial location and histology to identify spatial domains and spatially variable genes by graph convolutional network. Nat. Methods 18, 1342–1351 (2021).

68. Long, Y. et al. Spatially informed clustering, integration, and deconvolution of spatial transcriptomics with GraphST. Nat. Commun. 14, 1155 (2023).

69. Zhu, Q., Shah, S., Dries, R., Cai, L. & Yuan, G. C. Identification of spatially associated subpopulations by combining scRNAseq and sequential fluorescence in situ hybridization data. Nature Biotechnology 2018 36:12 36, 1183–1190 (2018).

70. Dong, K. & Zhang, S. Deciphering spatial domains from spatially resolved transcriptomics with an adaptive graph attention auto-encoder. Nat. Commun. 13, 1739 (2022).

71. Chidester, B., Zhou, T., Alam, S. & Ma, J. SpiceMix enables integrative single-cell spatial modeling of cell identity. Nat. Genet. 55, 78–88 (2023).

72. Ma, Y. & Zhou, X. Spatially informed cell-type deconvolution for spatial transcriptomics. Nat. Biotechnol. 40, 1349–1359 (2022).

73. Adler, P. et al. Mining for coexpression across hundreds of datasets using novel rank aggregation and visualization methods. Genome Biol. 10, R139 (2009).

74. Lachmann, A. et al. Massive mining of publicly available RNA-seq data from human and mouse. Nat. Commun. 9, 1366 (2018).

75. Clough, E. & Barrett, T. The Gene Expression Omnibus Database. in 93–110 (2016). doi:10.1007/978-1-4939-3578-9_5.

76. Athar, A., et al. ArrayExpress update – from bulk to single-cell expression data. Nucleic Acids Res. 47, D711–D715 (2019).

77. Dries, R. et al. Giotto: a toolbox for integrative analysis and visualization of spatial expression data. Genome Biol. 22, 1–31 (2021).

78. Wolf, F. A., Angerer, P. & Theis, F. J. SCANPY: Large-scale single-cell gene expression data analysis. Genome Biol. 19, 1–5 (2018).

79. Harris, C. R. et al. Array programming with NumPy. Nature 585, 357–362 (2020).

80. Virshup, I., Rybakov, S., Theis, F. J., Angerer, P. & Wolf, F. A. anndata: Access and store annotated data matrices. J. Open Source Softw. 9, 4371 (2024).

81. Kivioja, T. et al. Counting absolute numbers of molecules using unique molecular identifiers. Nat. Methods 9, 72–74 (2012).

82. Gentleman, R. C. et al. Bioconductor: open software development for computational biology and bioinformatics. Genome Biol. 5, R80 (2004).

83. Aken, B. L. et al. The Ensembl gene annotation system. Database 2016, baw093 (2016).

84. Cameron Davidson-Pilon. tdigest: computing accurate estimates from streaming data or distributed data. Github (2023).

85. Dunning, T. & Ertl, O. Computing Extremely Accurate Quantiles Using t-Digests. (2019).

86. Virtanen, P. et al. SciPy 1.0: fundamental algorithms for scientific computing in Python. Nat. Methods 17, 261–272 (2020).

87. Wolf, T. et al. HuggingFace’s Transformers: State-of-the-art Natural Language Processing. (2020).

88. Sinha, K. et al. Masked Language Modeling and the Distributional Hypothesis: Order Word Matters Pre-training for Little. (2021).

89. Devlin, J., Chang, M.-W., Lee, K. & Toutanova, K. BERT: Pre-training of Deep Bidirectional Transformers for Language Understanding. (2018).

90. Ansel, J. et al. PyTorch 2: Faster Machine Learning Through Dynamic Python Bytecode Transformation and Graph Compilation. in 29th ACM International Conference on Architectural Support for Programming Languages and Operating Systems, Volume 2 (ASPLOS ’24) (ACM, 2024). doi:10.1145/3620665.3640366.

91. Rasley, J., Rajbhandari, S., Ruwase, O. & He, Y. DeepSpeed: System Optimizations Enable Training Deep Learning Models with Over 100 Billion Parameters. in Proceedings of the 26th ACM SIGKDD International Conference on Knowledge Discovery & Data Mining 3505–3506 (Association for Computing Machinery, New York, NY, USA, 2020). doi:10.1145/3394486.3406703.

92. Vaswani, A., et al. Attention Is All You Need. (2017).

93. Long, M. et al. Comparing Xenium 5K and Visium HD data from identical tissue slide at a pathological perspective. Journal of Experimental & Clinical Cancer Research 44, 219 (2025).

94. Ren, P. et al. Systematic Benchmarking of High-Throughput Subcellular Spatial Transcriptomics Platforms. Preprint at 10.1101/2024.12.23.630033 (2024).

95. Oliveira, M. F. de et al. High-definition spatial transcriptomic profiling of immune cell populations in colorectal cancer. Nat. Genet. 57, 1512–1523 (2025).

96. Strope, B. S., Pendleton, K. E., Bowie, W. Z., Echeverria, G. V & Zhu, Q. Xenomake: a pipeline for processing and sorting xenograft reads from spatial transcriptomic experiments. Bioinformatics 40, (2024).

97. Pusztai, L. et al. Durvalumab with olaparib and paclitaxel for high-risk HER2-negative stage II/III breast cancer: Results from the adaptively randomized I-SPY2 trial. Cancer Cell 39, 989–998.e5 (2021).

98. Wolf, T. et al. Transformers: State-of-the-Art Natural Language Processing. in Proceedings of the 2020 Conference on Empirical Methods in Natural Language Processing: System Demonstrations (eds. Liu, Q. & Schlangen, D.) 38–45 (Association for Computational Linguistics, Online, 2020). doi:10.18653/v1/2020.emnlp-demos.6.

99. Liaw, R., Liang, E., Nishihara Robert and Moritz, P., Gonzalez, J. E. & Stoica, I. Tune: A Research Platform for Distributed Model Selection and Training. arXiv preprint arXiv:1807.05118 (2018).

100. Moffat, A. & Zobel, J. Rank-biased precision for measurement of retrieval effectiveness. ACM Trans. Inf. Syst. 27, 1–27 (2008).

101. Tsopoulidis, N. et al. T cell receptor–triggered nuclear actin network formation drives CD4 ^+^ T cell effector functions. Sci. Immunol. 4, (2019).

102. Lochner, M., Berod, L. & Sparwasser, T. Fatty acid metabolism in the regulation of T cell function. Trends Immunol. 36, 81–91 (2015).

