## Supplementary Data 1 for "CancerSTFormer enables multi-scale analysis of spot-resolution spatial transcriptomes and dissects gene and immune regulatory responses to targeted therapies"

**Supplementary Data 1 (Visium)**

| Dataset | Num of Patients | Patient Description Table | Num of Samples | Clinal Outcome (Large Sample | Disease Type | Data Size | Technology Type | Predictive or Prognostic | Repository | Link To Data |
|---|---|---|---|---|---|---|---|---|---|---|
| *Bassiouni et al. Spatial Transcriptomic Analysis of a Diverse Patient Cohort Reveals a Conserved Architecture in Triple-Negative Breast Cancer. Cancer Research. 2023* | 15 AA 7 White | TRUE | 42 | Survival Reported | TNBC | 35 GB (tar) | Visium | Prognostic | GEO GSE210616 | https://www.ncbi.nlm.nih.gov/geo/query/acc.cgi?acc=GSE210616 |
| *Runmin Wei et al. Spatial charting of single-cell transcriptomes in tissues. Nature Biotechnology. 2022* | 2 | FALSE | 2 | Non-applicable | DCIS | 34.2 MB (zipped) | Visium | Non-applicable | GEO GSE181254 | https://www.ncbi.nlm.nih.gov/geo/query/acc.cgi?acc=GSE181254 |
| *Dalia Barkley. Cancer cell states recur across tumor types and form specific interactions with the tumor microenvironment. Nature Genetics. 2022.* | 10 | TRUE (only cancer type) | 10 | Non-applicable | carcinoma of the ovary (OVCA), endometrium (UCEC), breast (BRCA), prostate (PRAD), kidney | 22 MB(.gz) | Visium | Non-applicable | GEO GSE203612 | https://www.ncbi.nlm.nih.gov/geo/query/acc.cgi?acc=GSE203612 |
| *Wang, Y., Chen, D., Liu, Y. et al. Multidirectional characterization of cellular composition and spatial architecture in human multiple primary lung cancers. Cell Death Dis 14, 462 (2023)* | 3 | TRUE | 12 | Non-applicable | Multiple primary lung cancers (one squamous carcinoma and three adenocarcinomas) | 283.5 Mb (.tar) | Visium | Non-applicable | GEO GSE200916 | https://www.ncbi.nlm.nih.gov/geo/query/acc.cgi |
| *Davidson G, Helleux A, Vano YA, Lindner V, Fattori A, Cerciat M, Elaidi RT, Verkarre V, Sun CM, Chevreau C, Bennamoun M, Lang H, Tricard T, Fridman WH, Sautes-Fridman C, Su X, Plassard D, Keime C, Thibault-Carpentier C, Barthelemy P, Oudard SM, Davidson I, Malouf GG. Mesenchymal-like Tumor Cells and* | 2 | FALSE | 2 | Non-applicable | Renal Cell Carcinoma | 116.9 MB (.tar) | Visium | Non-applicable | GEO GSE210041 | https://www.ncbi.nlm.nih.gov/geo/query/acc.cgi?acc=GSE210041 |
| *Loh JW, Lee JY, Lim AH, et al. Spatial transcriptomics reveal topological immune landscapes of Asian head and neck angiosarcoma. Commun Biol. 2023* | 4 | TRUE | 4 | Non-applicable | Angiosarcomas head and neck region (AS-HN) | 568.8 Mb (tar) | Visium | Non-applicable | GEO GSE227469 | https://www.ncbi.nlm.nih.gov/geo/query/acc.cgi?acc=GSE227469 |
| *Brooke E. Sanders, Rebecca Wolsky, Elizabeth S. Doughty, Kristen L. Wells, Debashis Ghosh, Lisa Ku, Joseph G. Pressey, Benjamin G. Bitler, Lindsay W. Brubaker. Small cell carcinoma of the ovary* | 1 | FALSE | 8 | Recovered | novel case of small cell carcinoma of the ovary hypercalcemic type | 337.7 Mb (.tar) | Visium | Non-applicable | GEO GSE213699 | https://www.ncbi.nlm.nih.gov/geo/query/acc.cgi?acc=GSE213699 |
| *Liu T, Liu C, Yan M, et al. Single cell profiling of primary and paired metastatic lymph node tumors in breast cancer patients. Nat Commun. 2022* | 4 | FALSE | 4 | Non-applicable | lymph node metastasized tumors (breast cancer) | 405.3 Mb (.tar) | Visium | Non-applicable | GEO GSE190811 | https://www.ncbi.nlm.nih.gov/geo/query/acc.cgi?acc=GSE190811 |
| *Spatiotemporal Profiling Unveiling the Cellular Organization Patterns and Local Protumoral Immune Microenvironment Remodeling in Early Lung Adenocarcinoma Progression* | 7 | N/A | 7 | Non-applicable | Lung Adenocarcinoma | 175 Mb (zip) | Visium | Non-applicable | Zenodo | https://zenodo.org/records/8417887 |
| *RNA Spatial Sequencing of Colorectal Liver Metastases regarding their Histopathological Growth Patterns* | 6 | N/A | 6 | Non-applicable | Colorectal Cancer Liver Metastases | Unsure | Visium | Non-applicable | ArrayExpress E-MTAB-12043 | https://www.ebi.ac.uk/biostudies/arrayexpress/studies/E-MTAB-12043?query=spatial%20transcriptomics |
| *Ravi et al. Spatially resolved multi-omics deciphers bidirectional tumor-host interdependence in glioblastoma. 2022* | 20 | TRUE | 28 | N/A | Glioblastomas | 7.87 GB (zipped) | Visium | Non-applicable | Datadryad | https://doi.org/10.5061/dryad.h70rxwdmj |
| *Wu et al. A single-cell and spatially resolved atlas of human breast cancers. Nature Genetics. 2021* | 6 | TRUE | 6 | N/A | 11 ER+, 5 HER2+ and 10 TNBCs | 1.2 GB (tar.gz) | Visium | Non-applicable | Zenodo | https://zenodo.org/records/4739739 |
| *Moncada et al. Integrating microarray-based spatial transcriptomics and single-cell RNA-seq reveals tissue architecture in pancreatic ductal adenocarcinomas. Nature Biotechnology. 2020* | 2 | FALSE | 10 | Non-applicable | pancreatic ductal adenocarcinomas | 1 GB (.gz) | Visium | Non-applicable | GEO GSE111672 | https://www.ncbi.nlm.nih.gov/geo/query/acc.cgi?acc=GSE111672 |
| *Emelie Berglund et al. Spatial maps of prostate cancer transcriptomes reveal an unexplored landscape of heterogeneity. Nature Communications. 2018.* | 1 | FALSE | 12 | Non-applicable | prostate cancer | N/A | Visium | Non-applicable | Spatial DB | http://www.spatialomics.org/SpatialDB/st_29925878_browse.php |

| | | | | | | | | | | |
|---|---|---|---|---|---|---|---|---|---|---|
| *Amani V, Riemondy KA, Fu R, et al. Integration of single-nuclei RNA-sequencing, spatial transcriptomics and histochemistry defines the complex microenvironment of NF1-associated plexiform neurofibromas. Acta* | 4 | FALSE | 4 | Non-applicable | Human NF1-associated plexiform neurofibroma | 210.8 MB (.tar) | Visium | Non-applicable | GEO GSE232766 | https://www.ncbi.nlm.nih.gov/geo/query/acc.cgi?acc=GSE232766 |
| *Arora, R., Cao, C., Kumar, M. et al. Spatial transcriptomics reveals distinct and conserved tumor core and edge architectures that predict survival and targeted therapy response. Nat Commun 14, 5029 (2023)* | 10 | FALSE | 12 | Non-applicable | HPV-negative oral squamous cell carcinoma | 153.4 Mb (.tar) | Visium | Non-applicable | GEO GSE208253 | https://www.ncbi.nlm.nih.gov/geo/query/acc.cgi?acc=GSE208253 |
| *Visium Spatial transcriptomics analysis of human primary colorectal cancer* | 4 | TRUE | 4 | Non-applicable | colorectal cancer | 41.2 Gb | Visium | Non-applicable | GEO GSE226997 | https://www.ncbi.nlm.nih.gov/geo/query/acc.cgi?acc=GSE226997 |
| *Sans M, Makino Y, Min J, Rajapakshe KI, Yip-Schneider M, Schmidt CM, Hurd MW, Burks JK, Gomez JA, Thege FI, Fahrmann JF, Wolff RA, Kim MP, Guerrero PA, Maitra A. Spatial Transcriptomics of Intraductal Papillary* | 13 | FALSE | 13 | N/A | Intraductal papillary mucinous neoplasms (IPMN) of the pancreas | 216. Mb (.tar) | Visium | Non-applicable | GEO GSE233293 | https://0-www-ncbi-nlm-nih-gov.brum.beds.ac.uk/geo/query/acc.cgi?acc=GSE233293 |
| *Ji AL, Rubin AJ, Thrane K, et al. Multimodal Analysis of Composition and Spatial Architecture in Human Squamous Cell Carcinoma [published correction appears in Cell. 2020* | 4 | FALSE | 12 | Non-applicable | cutaneous squamous cell carcinoma | 170 Kb (.tar) estimate | Visium | Non-applicable | GEO GSE144240 | https://www.ncbi.nlm.nih.gov/geo/query/acc.cgi?acc=GSE144240 |
| *Glioma spatialomics dataset* | 19 | N/A | 18 | Non-applicable | Glioblastoma | ~3 Gb (zip) | Visium | Non-applicable | Zenodo | https://zenodo.org/records/8105467 |
| *Spatial transcriptomic analysis of Sonic Hedgehog Medulloblastoma identified that loss of heterogeneity and promotion of differentiation underlies the response to CDK4/6 inhibition* | 4 | N/A | 4 | Non-applicable | Sonic Hedgehog Medulloblastoma | Unsure | Visium | Non-applicable | ArrayExpress E-MTAB-11720 | https://www.ebi.ac.uk/biostudies/arrayexpress/studies/E-MTAB-11720?query=spatial%20transcriptomics |
| *Lisa J Sudmeier et al. Distinct phenotypic states and spatial distribution of CD8+ T cell clonotypes in human brain metastases. Cell Report Medicine. 2022* | 6 | FALSE | 6 | Non-applicable | Brain Metastasis with various primary sites [ 6 lung, 6 breast, 4 Melanoma, 7 other ] | 3 GB (zipped) | Visium | Non-applicable | GEO GSE179572 | https://www.ncbi.nlm.nih.gov/geo/query/acc.cgi?acc=GSE179572 |
| *Andersson et al. Spatial deconvolution of HER2-positive breast cancer delineates tumor-associated cell type interactions. Nature Communications. 2021* | 8 | FALSE | 36 | Non-applicable | HER2-positive | 2 GB (zipped) Processed Count | Visium | Non-applicable | Zenodo | https://zenodo.org/records/3957257 |
| *Daniel Cuio Zhou et al. Spatially restricted drivers and transitional cell populations cooperate with the microenvironment in untreated and chemo-resistant pancreatic cancer. Nature Genetics. 2022* | 12 | TRUE | 12 | Outcome reported for 4 patients | pancreatic ductal adenocarcinomas | Unsure | Visium | Non-applicable | Human Tumor Atlas Network (HTAN) | https://www.ncbi.nlm.nih.gov/projects/gap/cgi-bin/study.cgi?study_id=phs002371.v1.p1 |
| *Erickson et al. Spatially resolved clonal copy number alterations in benign and malignant tissue. Nature 2022* | 2 | FALSE | 21 | Non-applicable | prostate cancer | 5 GB(zipped) | Visium | Non-applicable | Mendeley | https://data.mendeley.com/datasets/svw96g68dv/1 |
| *Zhang S, Yuan L, Danilova L, et al. Spatial transcriptomics analysis of neoadjuvant cabozantinib and nivolumab in advanced hepatocellular carcinoma identifies independent mechanisms of resistance and recurrence. Preprint. BioRxiv. 2023* | 7 ( 4 Responder 3 Non-responder ) | FALSE | 7 | Non-applicable | hepatocellular carcinoma (HCC, treated) | 241.3 Mb (.tar) | Visium | Predictive | GEO GSE238264 | https://0-www-ncbi-nlm-nih-gov.brum.beds.ac.uk/geo/query/acc.cgi?acc=GSE238264 |
| *Watanabe R, Miura N, Kurata M, Kitazawa R, Kikugawa T, Saika T. Spatial Gene Expression Analysis Reveals Characteristic Gene Expression Patterns of De Novo Neuroendocrine Prostate Cancer Coexisting with Androgen Receptor Pathway Prostate Cancer. Int J Mol Sci. 2023* | 1 | FALSE | 1 | Non-applicable | Neuroendocrine prostate carcinoma with androgen receptor pathway-positive adenocarcinoma of the prostate | 129.9 mb (.tar) | Visium | Non-applicable | GEO GSE230282 | https://www.ncbi.nlm.nih.gov/geo/query/acc.cgi?acc=GSE230282 |
| *Pavel I, Irina L, Tatiana G, et al. Comparison of the Illumina NextSeq 2000 and GeneMind Genolab M sequencing platforms for spatial transcriptomics. BMC Genomics. 2023* | N/A | FALSE | 6 | Non-applicable | ovarian cancer | 88.2 Mb (.tar) | Visium | Non-applicable | GEO GSE227019 | https://www.ncbi.nlm.nih.gov/geo/query/acc.cgi?acc=GSE227019 |
| *Cassier PA, Navaridas R, Bellina M, et al. Netrin-1 blockade inhibits tumour growth and EMT features in endometrial cancer. Nature. 2023* | 2 | TRUE | 4 | Translational Study | human endometrial carcinomas (ECs) treated with anti-netrin-1 antibody (NP137) | 571.0 Mb | Visium | Predictive | GEO GSE225690 | https://www.ncbi.nlm.nih.gov/geo/query/acc.cgi?acc=GSE225690 |

| | | | | | | | | | | |
|---|---|---|---|---|---|---|---|---|---|---|
| *Spatial transcriptomimcs in HCC* | 2 | N/A | 2 | Non-applicable | hepatocellular carcinoma (two HCC and normal adjacent tissues) | 52 Mb (zip) | Visium | Non-applicable | Zenodo | https://zenodo.org/records/7785709 |
| *Spatial transcriptomics reveals ovarian cancer subclones with different microenvironments* | 8 | N/A | 8 | Non-applicable | ovarian carcinoma (HGSOC) | 283 Mb | Visium | Non-applicable | GEO GSE211956 | https://www.ncbi.nlm.nih.gov/geo/query/acc.cgi?acc=GSE211956 |
| *A robust experimental and computational framework analysis at multiple resolutions, modalities and coverages* | 3 | N/A | 5 | Non-applicable | basal cell carcinoma (BCC) and squamous cell carcinoma (SCC) | Unsure | Visium | Non-applicable | ArrayExpress E-MTAB-11932 | https://www.ebi.ac.uk/biostudies/arrayexpress/studies/E-MTAB-11932?query=spatial%20transcriptomics |
| *Coutant et al. Spatial transcriptomics reveal pitfalls and opportunities for the detection of rare high-plasticity breast cancer subtypes. Laboratory Investigation. 2023* | 15 | FALSE | 15 | Non-applicable | 3 putative claudin-low (CL) tumors (CL-like) and 4 non-CL, genomically unstable TNBC samples 4 MpBCs | 652 (tar) | Visium | Non-applicable | GEO GSE213688 | https://www.ncbi.nlm.nih.gov/geo/query/acc.cgi?acc=GSE213688 |
| *Tokura et al. Single-Cell Transcriptome Profiling Reveals Intratumoral Heterogeneity and Molecular Features of Ductal Carcinoma In Situ. Cancer Research. 2022* | 1 | FALSE | 1 | N/A | DCIS | 150 Mb | Visium | Non-applicable | GEO GSE196208 | https://www.ncbi.nlm.nih.gov/geo/query/acc.cgi |
| *Anna Lyubetskaya et al. Assessment of spatial transcriptomics for oncology discovery. Cell reports methods. 2022. (focus on human portion only)* | 3 | FALSE | 3 | Non-applicable | pancreatic ductal adenocarcinomas | 5 GB(zipped) | Visium | Non-applicable | GEO GSE211895 | https://www.ncbi.nlm.nih.gov/geo/query/acc.cgi?acc=GSE211895 |
| *Andrwe L Ji. Multimodal Analysis of Composition and Spatial Architecture in Human Squamous Cell Carcinoma. Cell 2020.* | 4 | FALSE | 12 | Non-applicable | Squamous Cell Carcinoma | 164 MB(.gz) | Visium | Non-applicable | GEO GSE144240 | https://www.ncbi.nlm.nih.gov/geo/query/acc.cgi?acc=GSE144240 |
| *Moeyersoms AHM, Gallo RA, Zhang MG, et al. Spatial Transcriptomics Identifies Expression Signatures Specific to Lacrimal Gland Adenoid Cystic Carcinoma Cells. Cancers (Basel). 2023;15(12):3211. Published 2023 Jun 16. doi:10.3390/cancers15123211* | 1 | FALSE | 1 | Non-applicable | Lacrimal gland adenoid cystic carcinoma (LGACC post treatment) | 262.3 (.tar) | Visium | Non-applicable | GEO GSE228685 | https://0-www-ncbi-nlm-nih-gov.brum.beds.ac.uk/geo/query/acc.cgi?acc=GSE228685 |
| *Guo C, Qu X, Tang X, et al. Spatiotemporally deciphering the mysterious mechanism of persistent HPV-induced malignant transition and immune remodelling from HPV-infected normal cervix, precancer to cervical cancer: Integrating single-cell RNA-sequencing and spatial transcriptome. Clin Transl Med. 2023* | 9 | FALSE | 4 | Non-applicable | Normal Cervix, HPV infection, high-grade squamous intraepithelial lesions (HSIL) + HPV and cervical cancer + HPV | 222.0 Mb (tar) | Visium | Non-applicable | GEO GSE208654 | https://www.ncbi.nlm.nih.gov/geo/query/acc.cgi?acc=GSE208654 |
| *Fu R, Norris GA, Willard N, et al. Spatial transcriptomic analysis delineates epithelial and mesenchymal subpopulations and transition stages in childhood ependymoma. Neuro Oncol. 2023* | 14 | FALSE | 14 | Non-applicable | ependymoma posterior fossa subgroup A (PFA) (11 primary; 3 matched recurrence) | 4.2 Gb (.tar) | Visium | Non-applicable | GEO GSE195661 | https://www.ncbi.nlm.nih.gov/geo/query/acc.cgi?acc=GSE195661 |
| *Cheng HY, Hsieh CH, Lin PH, Chen YT, Hsu DS, Tai SK, Chu PY, Yang MH. Snail-regulated exosomal microRNA-21 suppresses NLRP3 inflammasome activity to enhance cisplatin resistance. J Immunother Cancer. 2022* | 2 | TRUE | 8 | Non-applicable | Human head and neck squamous cell carcinoma (HNSCC) | 343.8 Mb | Visium | Non-applicable | GEO GSE181300 | https://www.ncbi.nlm.nih.gov/geo/query/acc.cgi?acc=GSE181300 |
| *Charting the Heterogeneity of Colorectal Cancer Consensus Molecular Subtypes using Spatial Transcriptomics, View ORCID ProfileAlberto Valdeolivas et. al. (Preprint)* | 7 | FALSE | 14 | Non-applicable | colorectal cancer | 2.1 Gb (zip) | Visium | Non-applicable | Zenodo | https://zenodo.org/records/7760264 |
| *Defining the immunophenotype within the clear cell renal cell carcinoma tumour microenvironment to guide immune checkpoint blockade Nicholas Matigan et. al.* | 9 | FALSE | 9 | Non-applicable | clear cell renal cell carcinoma (ccRCC) | ~3 Gb (zip) | Visium | Non-applicable | ArrayExpress E-MTAB-12767 | https://www.ebi.ac.uk/biostudies/arrayexpress/studies/E-MTAB-12767?query=Defining%20the%20immunophenotype%20within%20the%20clear%20cell%20renal%20cell%20carcino |
| *Fine mapping of cell states across anatomical sites in healthy human skin and basal cell carcinoma - Spatial Transcriptomics - 10X Visium* | 8 (BCC), rest normal | N/A | 22 | Non-applicable | basal cell carcinoma (BCC) and healthy [ Has healthy samples to be removed hard to tell which is which ] | Unsure | Visium | Non-applicable | ArrayExpress E-MTAB-13085 | https://www.ebi.ac.uk/biostudies/arrayexpress/studies/E-MTAB-13085?query=Multi-scale%20spatial%20mapping%20of%20cell%20populations%20across%20anatomical%20sites%20in%20healthy%20h[illegible] |

**Supplementary Data 1 Visium 10X**

| Dataset | Num of Samples | Disease Type | Stage | Extra Info | Data Size | Link | File Name |
|---|---|---|---|---|---|---|---|
| *Human Brain Cancer, 11 mm Capture Area (FFPE)* | 1 | Glioblastoma multiforme | N/A | N/A | ~250 Mb | https://www.10xgenomics.com/resources/datasets/human-brain-cancer-11-mm-capture-area-ffpe-2-standard | 10X_HumanBrainCancer.tar.gz |
| *Human Ovarian Cancer, 11 mm Capture Area (FFPE)* | 1 | Ovarian Serous Carcinoma, High Grade | N/A | N/A | ~250 Mb | https://www.10xgenomics.com/resources/datasets/human-ovarian-cancer-11-mm-capture-area-ffpe-2-standard | 10X_HumanOvarianCancer.tar.gz |
| *Human Breast Cancer: Visium Fresh Frozen, Whole Transcriptome* | 1 | Breast Cancer | AJCC/UICC Stage T2N0M0, ER positive, PR negative, | N/A | ~250 Mb | https://www.10xgenomics.com/resources/datasets/fresh-frozen-visium-on-cytassist-human-breast-cancer-probe-based-whole-transcriptome-profiling-2-standard | 10X_HumanBreastCancer.tar.gz |
| *Human Prostate Cancer, Acinar Cell Carcinoma (FFPE)* | 1 | nvasive Acinar Cell Carcinoma | Stage IV | Total Gleason score: 7 | ~250 Mb | https://www.10xgenomics.com/resources/datasets/human-prostate-cancer-acinar-cell-carcinoma-ffpe-1-standard | 10X_HumanProstateCancer.tar.gz |
| *Human Ovarian Cancer: Whole Transcriptome Analysis. Stains: DAPI, Anti-PanCK, Anti-CD45* | 1 | Endometrial Adenocarcinoma of the ovary tissue | N/A | Samples were stained with antibodies and DAPI | ~250 Mb | https://www.10xgenomics.com/resources/datasets/human-ovarian-cancer-whole-transcriptome-analysis-stains-dapi-anti-pan-ck-anti-cd-45-1-standard-1-2-0 | 10X_HumanEndometrialAdenocarcinoma.tar.gz |
| *Human Glioblastoma: Whole Transcriptome Analysis* | 1 | glioblastoma multiforme | N/A | N/A | ~250 Mb | https://www.10xgenomics.com/resources/datasets/human-glioblastoma-whole-transcriptome-analysis-1-standard-1-2-0 | 10X_HumanGlioblastoma.tar.gz |
| *Human Breast Cancer: Whole Transcriptome Analysis* | 1 | Invasive Lobular Carcinoma breast tissue | Stage Group 1 | AJCC/UICC Stage Group I, ER positive, PR positive, HER2 negative. | ~250 Mb | https://www.10xgenomics.com/resources/datasets/human-breast-cancer-whole-transcriptome-analysis-1-standard-1-2-0 | 10X_HumanLobularCarcinoma.tar.gz |
| *Human Lung Cancer, 11 mm Capture Area (FFPE)* | 1 | Lung Cancer, Neuroendocrine Carcinoma | (AJCC): IB | TNM system: T2aN0MX | ~250 Mb | https://www.10xgenomics.com/resources/datasets/human-lung-cancer-11-mm-capture-area-ffpe-2-standard | /ST.compendium/Extra_Visium_Sets/CytAssist_11mm_FFPE_Human_Lung_Cancer |
| *Human Lung Cancer (FFPE)* | 1 | Lung, Squamous Cell Carcinoma | N/A | N/A | ~250 Mb | https://www.10xgenomics.com/resources/datasets/human-lung-cancer-ffpe-2-standard | /ST.compendium/Extra_Visium_Sets/CytAssist_FFPE_Human_Lung_Squamous_Cell_Carcinoma |
| *Human Ovarian Cancer (FFPE)* | 1 | serous papillary carcinoma of human ovarian. | N/A | N/A | ~250 Mb | https://www.10xgenomics.com/resources/datasets/human-ovarian-cancer-1-standard | /ST.compendium/Extra_Visium_Sets/Visium_FFPE_Human_Ovarian_Cancer_ |
| *Human Colorectal Cancer, 11 mm Capture Area (FFPE)* | 1 | Colorectal cancer, Adenocarcinoma | (AJCC): II-A | TNM system: T3 | ~250 Mb | https://www.10xgenomics.com/resources/datasets/human-colorectal-cancer-11-mm-capture-area-ffpe-2-standard | /ST.compendium/Extra_Visium_Sets/CytAssist_11mm_FFPE_Human_Colorectal_Cancer |
| *Human Melanoma, IF Stained (FFPE)* | 1 | Skin, Malignant Melanoma | N/A | IF stained | ~200 Mb | https://www.10xgenomics.com/resources/datasets/human-melanoma-if-stained-ffpe-2-standard | /ST.compendium/Extra_Visium_Sets/Visium_FFPE_Human_Cervical_Cancer |
| *Human Cervical Cancer (FFPE)* | 1 | squamous cell carcinoma of human cervical cancer | N/A | N/A | ~250 Mb | https://www.10xgenomics.com/resources/datasets/human-cervical-cancer-1-standard | /ST.compendium/Extra_Visium_Sets/CytAssist_FFPE_Human_Skin_Melanoma |

| | | | | | | | |
|---|---|---|---|---|---|---|---|
| *Human Breast Cancer: Ductal Carcinoma In Situ, Invasive Carcinoma (FFPE)* | 1 | Ductal Carcinoma In Situ, Invasive Carcinoma | N/A | Asian Female | ~250 Mb | https://www.10xgenomics.com/resources/datasets/human-breast-cancer-ductal-carcinoma-in-situ-invasive-carcinoma-ffpe-1-standard-1-3-0 | /ST.compendium/Extra_Visium_Sets/Visium_FFPE_Human_Breast_Cancer |
| *Human Colorectal Cancer: Whole Transcriptome Analysis* | 1 | Invasive Adenocarcinoma of the large intestine | N/A | N/A | ~250 Mb | https://www.10xgenomics.com/resources/datasets/human-colorectal-cancer-whole-transcriptome-analysis-1-standard-1-2-0 | /ST.compendium/Extra_Visium_Sets/Parent_Visium_Human_ColorectalCancer_ |
| *Human Breast Cancer (Block A Section 2)* | 1 | Invasive Ductal Carcinoma breast tissue | Stage II | AJCC/UICC Stage Group IIA, ER positive, PR negative, Her2 positive | ~250 Mb | https://www.10xgenomics.com/resources/datasets/human-breast-cancer-block-a-section-2-1-standard-1-1-0 | /ST.compendium/Extra_Visium_Sets/V1_Breast_Cancer_Block_A_Section_2 |
| *Visium CytAssist Gene and Protein Expression Library of Human Breast Cancer, IF, 6.5mm (FFPE)* | 1 | Human Breast Cancer | N/A | N/A | ~500 Mb | https://www.10xgenomics.com/resources/datasets/gene-and-protein-expression-library-of-human-breast-cancer-cytassist-ffpe-2-standard | /project/ST.compendium/10X_CytAssist_FFPE_Protein_Expression_Human_Breast_Cancer |
| *Fresh Frozen Visium on CytAssist: Human Breast Cancer, Probe-Based Whole Transcriptome Profiling* | 1 | Breast Cancer | N/A | N/A | ~250 Mb | https://www.10xgenomics.com/resources/datasets/fresh-frozen-visium-on-cytassist-human-breast-cancer-probe-based-whole-transcriptome-profiling-2-standard | /project/ST.compendium/10X_CytAssist_Fresh_Frozen_Human_Breast_Cancer |
| *Human Intestine Cancer (FPPE)* | 1 | Human Intestinal Cancer | N/A | N/A | ~250 Mb | https://www.10xgenomics.com/resources/datasets/human-intestine-cancer-1-standard | /project/ST.compendium/Visium_FFPE_Human_Intestinal_Cancer_ |
| *Invasive Ductal Carcinoma Stained With Fluorescent CD3 Antibody* | 1 | Invasive ductal carcinoma | Tumor Grade – III | AJCC/UICC Stage - T2N0M0 AJCC/UICC Stage Group – IIA ER – Positive PR – | ~250 Mb | https://www.10xgenomics.com/resources/datasets/invasive-ductal-carcinoma-stained-with-fluorescent-cd-3-antibody-1-standard-1-2-0 | /project/ST.compendium/Visium_V1_Human_Invasive_Ductal_Carcinoma_ |
| *Human Breast Cancer (Block A Section 1)* | 1 | Invasive Ductal Carcinoma breast tissue | Stage II | AJCC/UICC Stage Group IIA, ER positive, PR negative, Her2 positive | ~250 Mb | https://www.10xgenomics.com/resources/datasets/human-breast-cancer-block-a-section-1-1-standard-1-1-0 | /project/ST.compendium/Visium_V1_Breast_Cancer_Block_A_Section_1_ |

**Supplementary Data 1 Other Platforms**

| Publication | Dataset | Disease | Platform | Number of samples |
|---|---|---|---|---|
| A multi-modal single-cell and spatial expression map of metastatic breast cancer biopsies across clinicopathological features. Nature Medicine | HTAPP | Breast Cancer | Slideseq | 15 |
| A spatial cell atlas of neuroblastoma reveals developmental, epigenetic and spatial axis of tumor heterogeneity. Biorxiv | HTAPP | Neuroblastoma | Slideseq | 19 |
| Biermann J, Melms JC, Amin AD, Wang Y et al. Dissecting the treatment-naive ecosystem of human melanoma brain metastasis. Cell | GSE20078 | Melanoma | Slideseq | 16 |
| Multimodal single-cell and whole-genome sequencing of small, frozen clinical specimens. *Nat Genet* 2023 Jan;55(1):19-25. | GSE216055 | Melanoma | Slideseq | 3 |
| Spatially exploring RNA biology in archival formalin-fixed paraffin-embedded tissues. Cell 2024 Nov 14;187(23):6760-6779.e24. PMID: 39353436 | GSE274641 | Lymphoma | Dbit-seq | 5 |
| Spatial genomics enables multi-modal study of clonal heterogeneity in tissues. Nature | Broad Institute SCP portal SCP1278 | Colon Cancer | Slideseq | 1 |

### Supplementary Data 1 (Summary)

| Samples | Studies | |
|---|---|---|
| 431 | 43 | Visium public domain |
| 21 | 1 | Visium 10X Genomics |
| 54 | 5 | Slideseq |
| 5 | 1 | Dbitseq |
