## Supplementary Data 2 for "CancerSTFormer enables multi-scale analysis of spot-resolution spatial transcriptomes and dissects gene and immune regulatory responses to targeted therapies"

**Supplementary Data 2. CancerSTFormer parameter tuning**
**Transfer learning hyperparameter tuning settings**
**Treatment sensitive and resistant gene predictions**

| | Gene-sets to train | Learning Rate | Weight Decay | Epochs | Trials | Training ST dataset | LR scheduler | Warmup steps | Seed | Tuner |
|---|---|---|---|---|---|---|---|---|---|---|
| 50um Local Model | PD-1 Sensitive | (1e-3, 1e-2) | (0.01, 0.05) | 1 | 60 | TNBC dataset | (linear, cosine, polynomial) | (5, 50) | (0, 100) | Ray |
| 50um Local Model | PD-1 Resistant | (1e-3, 1e-2) | (0.01, 0.05) | 1 | 60 | TNBC dataset | (linear, cosine, polynomial) | (5, 50) | (0, 100) | Ray |
| 250um Extended Model | PD-1 Sensitive | (1e-3, 1e-2) | (0.01, 0.05) | 1 | 60 | TNBC dataset | (linear, cosine, polynomial) | (5, 50) | (100, 1000) | Ray |
| 250um Extended Model | PD-1 Resistant | (1e-3, 1e-2) | (0.01, 0.05) | 1 | 60 | TNBC dataset | (linear, cosine, polynomial) | (5, 50) | (100, 1000) | Ray |
| 50um Local Model | IGF1R Sensitive | (1e-3, 1e-2) | (0.01, 0.05) | 1 | 60 | TNBC dataset | (linear, cosine, polynomial) | (5, 50) | (0, 100) | Ray |
| 50um Local Model | IGF1R Resistant | (1e-3, 1e-2) | (0.01, 0.05) | 1 | 60 | TNBC dataset | (linear, cosine, polynomial) | (5, 50) | (0, 100) | Ray |
| 250um Extended Model | IGF1R Sensitive | (1e-3, 1e-2) | (0.01, 0.05) | 1 | 60 | TNBC dataset | (linear, cosine, polynomial) | (5, 50) | (100, 1000) | Ray |
| 250um Extended Model | IGF1R Resistant | (1e-3, 1e-2) | (0.01, 0.05) | 1 | 60 | TNBC dataset | (linear, cosine, polynomial) | (5, 50) | (100, 1000) | Ray |
| 50um Local Model | ANG1-Sensitive | (1e-3, 1e-2) | (0.01, 0.05) | 1 | 60 | TNBC dataset | (linear, cosine, polynomial) | (5, 50) | (0, 100) | Ray |
| 50um Local Model | ANG1-Resistant | (1e-3, 1e-2) | (0.01, 0.05) | 1 | 60 | TNBC dataset | (linear, cosine, polynomial) | (5, 50) | (0, 100) | Ray |
| 250um Extended Model | ANG1-Sensitive | (1e-3, 1e-2) | (0.01, 0.05) | 1 | 60 | TNBC dataset | (linear, cosine, polynomial) | (5, 50) | (100, 1000) | Ray |
| 250um Extended Model | ANG1-Resistant | (1e-3, 1e-2) | (0.01, 0.05) | 1 | 60 | TNBC dataset | (linear, cosine, polynomial) | (5, 50) | (100, 1000) | Ray |

**Tissue type spot-level predictions**

| | Tissue types to train | Learning Rate | Weight Decay | Epochs | Trials | Training dataset | LR scheduler | Warmup Ratio | Seed | Tuner |
|---|---|---|---|---|---|---|---|---|---|---|
| 50um Local Model | 9 tissue types | (1e-5, 1e-3) | (0, 0.3) | 1 | 32 | Balanced multi-tissue ST dataset, 4 samples per tissue, FFPE+Fresh Frozen samples represented | cosine | (0, 0.3) | 42 | Ray |

**Metastasis gene prediction**

| | Gene-sets to train | Learning Rate | Weight Decay | Epochs | Trials | Training ST dataset | LR scheduler | Warmup steps | Seed | Tuner |
|---|---|---|---|---|---|---|---|---|---|---|
| 50um Local Model | Lung-met specific genes | 1.00E-05 | 0.35 | 30 | 30 | Entire compendium | polynomial | 500 | 100 | None, single-setting |
| 50um Local Model | Brain-met specific genes | (1e-6, 1e-5) | (0.15, 0.35) | 10 | 30 | Entire compendium | (linear, cosine, polynomial) | (500, 2000) | (0, 100) | Ray |
| 50um Local Model | Bone-met specific genes | (3e-6, 3e-4) | (0.02, 0.35) | 10 | 10 | Entire compendium | (linear, cosine, polynomial) | (500, 2000) | (0, 100) | Ray |
| 250um Extended Model | Lung-met specific genes | (1e-3, 1e-2) | (0.01, 0.05) | 1 | 30 | Entire compendium | (linear, cosine, polynomial) | (5, 50) | (0, 100) | Ray |
| 250um Extended Model | Brain-met specific genes | (1e-3, 1e-2) | (0.01, 0.05) | 1 | 30 | Entire compendium | (linear, cosine, polynomial) | (5, 50) | (0, 100) | Ray |
| 250um Extended Model | Bone-met specific genes | (1e-3, 1e-2) | (0.01, 0.05) | 1 | 30 | Entire compendium | (linear, cosine, polynomial) | (5, 50) | (0, 100) | Ray |

**Responder vs nonresponder prediction**

| | Types to predict | Learning Rate | Weight Decay | Epochs | Trials | Training dataset | LR scheduler | Warmup steps | Seed | Tuner |
|---|---|---|---|---|---|---|---|---|---|---|
| 50um Local Model | Responder ST samples versus Nonresponder ST samples | (0.0001, 0.001) | (0.25, 0.35) | 3 | 10 | GSE238264 | (linear, cosine) | (100, 1000) | (0, 100) | Ray |

**Race prediction**

| | Types to predict | Learning Rate | Weight Decay | Epochs | Trials | Training dataset | LR scheduler | Warmup steps | Seed | Tuner |
|---|---|---|---|---|---|---|---|---|---|---|
| 50um Local Model | AA vs EA | (0.0002, 0.001) | (0.25, 0.35) | 5 | 10 | GSE210616 | (linear, cosine) | (400, 1000) | (10, 100) | Ray |
