## Supplementary Data 3 for "CancerSTFormer enables multi-scale analysis of spot-resolution spatial transcriptomes and dissects gene and immune regulatory responses to targeted therapies"

**Supplementary Data 3 Metastasis genes (Lung)**

**Proteins/genes involved in breast cancer lung metastasis, and experimental evidences**

| protein/gene name | cancer type | species | paper title |
|---|---|---|---|
| BRMS1 | lung metastasis | human | Breast cancer metastasis suppressor 1 coordinately regulates metastasis-associated microRNA expression |
| NF-κB | lung metastasis | human | The role of NF-κB in breast cancer initiation, growth, metastasis, and resistance to chemotherapy |
| Lin28B | lung metastasis | human | Lin28B-high breast cancer cells promote immune suppression in the lung pre-metastatic niche via exosomes and support cancer progression |
| CECR2 | lung metastasis | human and mouse | CECR2 drives breast cancer metastasis by promoting NF-κB signaling and macrophage-mediated immune suppression |
| TGF-β2 | lung metastasis | human and mouse | Lung-resident alveolar macrophages regulate the timing of breast cancer metastasis |
| Nm23-H1 (NME1) | lung metastasis | human | Activation of Nm23-H1 to suppress breast cancer metastasis via redox regulation |
| RARRES3 | lung metastasis | human | *RARRES3* suppresses breast cancer lung metastasis by regulating adhesion and differentiation |
| Stat3, TGF-β, FoxP3 | lung metastasis | mouse (4T1 breast cancer model) | Tumor-Evoked Regulatory B Cells Promote Breast Cancer Metastasis by Converting Resting $CD4^+$ T Cells to T-Regulatory Cells |
| STAT3 | lung metastasis | human | STAT3 as a potential therapeutic target in triple negative breast cancer: a systematic review |
| BC069792, KCNQ4, JAK2, p-AKT | lung metastasis | human and mouse | LncRNA-BC069792 suppresses tumor progression by targeting KCNQ4 in breast cancer |
| FOXP2, miR-199a | lung metastasis | human | MSC-Regulated MicroRNAs Converge on the Transcription Factor FOXP2 and Promote Breast Cancer Metastasis |
| ATP11B, PTDSS2, BRCA1 | lung metastasis | human | ATP11B inhibits breast cancer metastasis in a mouse model by suppressing externalization of nonapoptotic phosphatidylserine |
| CXCR3, CXCR4, CCR7 | lung metastasis | Human | Regulation of Tumor and Metastasis Initiation by Chemokine Receptors |
| SDPR | lung metastasis | Human and Mouse (NOD/SCID mouse model used) | SDPR functions as a metastasis suppressor in breast cancer by promoting apoptosis |
| CYR61 | lung metastasis | human | The matricellular protein CYR61 promotes breast cancer lung metastasis by facilitating tumor cell extravasation and suppressing anoikis |
| NR2F1 | lung metastasis | human and mouse | An NR2F1-specific agonist suppresses metastasis by inducing cancer cell dormancy |
| WDR5 | lung metastasis | human | Human WDR5 promotes breast cancer growth and metastasis via KMT2-independent translation regulation |
| IMP1 (IGF2BP1), PTGS2, GDF15, IGF-2 | lung metastasis | Human and Mouse (Mouse xenograft model used) | IMP1 suppresses breast tumor growth and metastasis through the regulation of its target mRNAs |
| IL13Rα2, INHBA, Smad2, Smad3, Akt | lung metastasis | Human and Mouse (In vivo lung metastasis model used) | Activin A Signaling Regulates IL13Rα2 Expression to Promote Breast Cancer Metastasis |
| PTGS2 | lung metastasis | mouse | Immunosuppressive reprogramming of neutrophils by lung mesenchymal cells promotes breast cancer metastasis |
| ULK1, ATG4b | lung metastasis | Human and Mouse (MDA-MB-231 xenograft model) | Selective Reversible Inhibition of Autophagy in Hypoxic Breast Cancer Cells Promotes Pulmonary Metastasis |
| HOXB9, VEGF, bFGF, IL-8, ANGPTL-2, ErbB, TGF-β | lung metastasis | Human and Mouse (in vivo lung metastasis model used) | *HOXB9* , a gene overexpressed in breast cancer, promotes tumorigenicity and lung metastasis |
| AMPKα1, PI3K, HER2, ΔNp63α, E-cadherin | lung metastasis | Human and Mouse (in vivo metastasis model used) | Transcriptional suppression of AMPKα1 promotes breast cancer metastasis upon oncogene activation |
| GALNT14, KRAS, PI3K, c-JUN, BMPs, FGFs | lung metastasis | human | GALNT14 promotes lung-specific breast cancer metastasis by modulating self-renewal and interaction with the lung microenvironment. |
| MYC, PIK3CA, TP53 | lung metastasis | human | The Metabolic Mechanisms of Breast Cancer Metastasis |
| RASAL2, RAC1, ARHGAP24, miR-203 | lung metastasis | human | RASAL2 activates RAC1 to promote triple-negative breast cancer progression |

**Supplementary Data 3 Metastasis genes (Brain)**

**Proteins/genes involved in breast cancer brain metastasis, and experimental evidences**

| protein/gene name | cancer type | species | paper title |
|---|---|---|---|
| HER2, STAT3, NF-κB, PD-L1, CD74, MIF, PI3K-Akt, xCT | brain metastasis | human | Molecular and cellular mechanisms underlying brain metastasis of breast cancer |
| TGLI1, CD44, Nanog, Sox2, OCT4 | brain metastasis | human and mouse | TGLI1 transcription factor mediates breast cancer brain metastasis via activating metastasis-initiating cancer stem cells and astrocytes in the tumor microenvironment |
| SOX2, FSCN1, HBEGF | brain metastasis | human (with possible | SOX2 Promotes Brain Metastasis of Breast Cancer by Upregulating the Expression of FSCN1 and HBEGF |
| Cx31, FAK, NF-κB, CX43, LAMA4, α3 integrin | brain metastasis | human and mouse | Connexins orchestrate progression of breast cancer metastasis to the brain by promoting FAK activation |
| AXL, Tenascin C | brain metastasis | human and mouse | Distinct tumor architectures and microenvironments for the initiation of breast cancer metastasis in the brain |
| HER2, EGFR, VEGF | brain metastasis | mouse | Metastasis of Breast Tumor Cells to Brain Is Suppressed by Phenethyl Isothiocyanate in a Novel *In Vivo* Metastasis Model |
| Extracellular vesicle-microRNAs (miRNAs) | brain metastasis | human | Crosstalk between breast cancer-derived microRNAs and brain microenvironmental cells in breast cancer brain metastasis |
| CACNA1H, CAMKII, p38 MAPK, β-catenin, HMGA2, miR-1246 | brain metastasis | Human and Mouse ( | Ca2+ and CACNA1H mediate targeted suppression of breast cancer brain metastasis by AM RF EMF |
| EZH2, Src, c-JUN, G-CSF, Arg1, PD-L1 | brain metastasis | human and mouse | Brain metastases recruit suppressive immune cells via novel pathway to facilitate their growth |
| Src, EGFR, HER2 | brain metastasis | human and mouse | Src Family Kinases as Novel Therapeutic Targets to Treat Breast Cancer Brain Metastases |
| miR-128, MTDH (AEG-1, Lyric) | brain metastasis | human | MiR-128 suppresses metastatic capacity by targeting metadherin in breast cancer cells |
| KISS1, CXCL12, MIR345, ATG5, ATG7, MMP9, IL8 | brain metastasis | human and mouse | Astrocytes promote progression of breast cancer metastases to the brain via a KISS1-mediated autophagy |
| STAT3, ERK1/2, p53, PD-L1 | brain metastasis | Human (MDA-MB-23 | Brain Metastases Are Regulated by Immuno-inflammatory Signaling Pathways Governed by STAT3, MAPK and Tumor Suppressor p53 Status: Possible Therapeutic Targets |
| AKT, HSF1 | brain metastasis | Human and Mouse ( | Combined inhibition of AKT and HSF1 suppresses breast cancer stem cells and tumor growth |
| XIST, c-Met, MSN, miRNA-503 | brain metastasis | human and mouse | Loss of XIST in Breast Cancer Activates MSN-c-Met and Reprograms Microglia via Exosomal miRNA to Promote Brain Metastasis |
| STAT3, ERK1/2, p53, PD-L1 | brain metastasis | Human (MDA-MB-23 | Brain Metastases Are Regulated by Immuno-inflammatory Signaling Pathways Governed by STAT3, MAPK and Tumor Suppressor p53 Status: Possible Therapeutic Targets |
| Gabra3, AKT | brain metastasis | human and mouse | Gene previously observed only in brain is important driver of metastatic breast cancer |

**Supplementary Data 3 Metastasis genes (Bone)**

**Proteins/genes involved in breast cancer bone metastasis, and experimental evidences**

| protein/gene name | cancer type | paper title |
|---|---|---|
| RKIP | bone metastasis | RKIP Suppresses Breast Cancer Metastasis to the Bone by Regulating Stroma-Associated Genes |
| BMP4 | bone metastasis | Activation of Canonical BMP4-SMAD7 Signaling Suppresses Breast Cancer Metastasis |
| RANKL | bone metastasis | Breast cancer metastasis to the bone: mechanisms of bone loss |
| IL-1B | bone metastasis | The role of IL-1B in breast cancer bone metastasis |
| DLC1 | bone metastasis | DLC1-dependent parathyroid hormone–like hormone inhibition suppresses breast cancer bone metastasis |
| Brachyury (Bry), SOX5 | bone metastasis | Transactivation of *SOX5* by Brachyury promotes breast cancer bone metastasis |
| RUNX2 | bone metastasis | Role of transcription factors in metastasis of breast cancer |
| VCAM1, TGF-β, Jagged1 | bone metastasis | Dissecting Tumor-Stromal Interactions in Breast Cancer Bone Metastasis |
| Cx43 (Connexin 43) | bone metastasis | Antibody-activation of connexin hemichannels in bone osteocytes with ATP release suppresses breast cancer and osteosarcoma malignancy |
| RANKL, RANK, β2AR | bone metastasis | Stimulation of Host Bone Marrow Stromal Cells by Sympathetic Nerves Promotes Breast Cancer Bone Metastasis in Mice |
| β-catenin, PPARγ | bone metastasis | Lipid Osteoclastokines Regulate Breast Cancer Bone Metastasis |
| RANKL, NF-κB | bone metastasis | Gold clusters prevent breast cancer bone metastasis by suppressing tumor-induced osteoclastogenesis |
| PD-1, TIGIT, IL1β | bone metastasis | Combining TIGIT Blockade with MDSC Inhibition Hinders Breast Cancer Bone Metastasis by Activating Antitumor Immunity |
| RANKL, PTHrP | bone metastasis | Molecular Regulation of Bone Metastasis Pathogenesis |
| EZH2, ITGB1, FAK, TGFβRI, TGFβRII | bone metastasis | EZH2 engages TGFβ signaling to promote breast cancer bone metastasis via integrin β1-FAK activation |
| CapG, STC-1, PRMT5, HGF, Met, VEGF, PI3K | bone metastasis | Novel mediators of breast cancer bone metastasis—insights from studies of gene-regulation and the global proteome |
| RANKL, MAPK | bone metastasis | Ononin Inhibits Tumor Bone Metastasis and Osteoclastogenesis By Targeting Mitogen-Activated Protein Kinase Pathway in Breast Cancer |
| ZEB1, FST, NOG, CHRDL1 | bone metastasis | The EMT-activator ZEB1 induces bone metastasis associated genes including BMP-inhibitors. |
| interleukin-11 and CTGF TGFβ | bone metastasis | A multigenic program mediating breast cancer metastasis to bone |
| *IL-11* | bone metastasis | Breast cancer bone metastasis mediated by the Smad tumor suppressor pathway |
| MAF PTHrP | bone metastasis | Enhanced MAF Oncogene Expression and Breast Cancer Bone Metastasis |
| RKIP | bone metastasis | RKIP Suppresses Breast Cancer Metastasis to the Bone by Regulating Stroma-Associated Genes |
| RUNX2 | bone metastasis | Breast tumor stiffness instructs bone metastasis via maintenance of mechanical conditioning |
| CapG and GIPC1 | bone metastasis | Novel mediators of breast cancer bone metastasis—insights from studies of gene-regulation and the global proteome |
| DDX3 | bone metastasis | Targeting RNA helicase DDX3X with a small molecule inhibitor for breast cancer bone metastasis treatment |
| IRF7 | bone metastasis | An Innate Immune Pathway Regulates Breast Cancer Metastasis |
| ET-1 DKK1 Wnt-5a, β-catenin, AXIN2, and LRP5 | bone metastasis | Wnt/β-Catenin Signaling Pathway Regulates Osteogenesis for Breast Cancer Bone Metastasis: Experiments in an *In Vitro* Nanoclay Scaffold Cancer Testbed |
| ABCC5 | bone metastasis | ABCC5 supports osteoclast formation and promotes breast cancer metastasis to bone |
| PTH1R | bone metastasis | Parathyroid hormone 1 receptor signaling mediates breast cancer metastasis to bone in mice |
| Osteoactivin matrix metalloproteinase-3 | bone metastasis | Osteoactivin Promotes Breast Cancer Metastasis to Bone |
| KDM1A MAF | bone metastasis | *MAF* amplification licenses ERα through epigenetic remodelling to drive breast cancer metastasis |
| ITGBL1 RUNX2 | bone metastasis | ITGBL1 Is a Runx2 Transcriptional Target and Promotes Breast Cancer Bone Metastasis by Activating the TGFβ Signaling Pathway |
| ITGA5 | bone metastasis | Integrin alpha5 in human breast cancer is a mediator of bone metastasis and a therapeutic target for the treatment of osteolytic lesions |
| NGF | bone metastasis | Nerve Growth Factor in Breast Cancer Cells Promotes Axonal Growth and Expression of Calcitonin Gene-related Peptide in a Rat Model of Spinal Metastasis |
| EZH2 | bone metastasis | EZH2 Engages TGFβ Signaling to Promote Breast Cancer Bone Metastasis via Integrin β1-FAK Activation |
| alpha v integrin | bone metastasis | Genetic depletion and pharmacological targeting of alpha v integrin in breast cancer cells impairs metastasis in zebrafish and mouse xenograft models |
| ABL kinases, TRAIL, STAT5, MMP1, IL-6, TAZ | bone metastasis | ABL kinases promote breast cancer osteolytic metastasis by modulating tumor-bone interactions through TAZ and STAT5 signaling |
| Siglec-15 | bone metastasis | Siglec-15/sialic acid axis as a central glyco-immune checkpoint in breast cancer bone metastasis |

Supplementary Data 3 Metastasis Genes Summary

**Proteins from previous worksheets (lung, brain, bone) were converted to genes, as shown below**

| **Lung metastasis genes** | **Brain metastasis genes** | **Bone metastasis genes** |
|---|---|---|
| BRMS1 | ERBB2 | PEBP1 |
| NFKB1 | STAT3 | BMP4 |
| RELA | NFKB1 | TNFSF11 |
| LIN28B | RELA | IL1B |
| CECR2 | CD274 | DLC1 |
| TGFB2 | CD74 | TBXT |
| NME1 | MIF | SOX5 |
| RARRES3 | PIK3CA | RUNX2 |
| STAT3 | AKT1 | VCAM1 |
| TGFB1 | SLC7A11 | TGFB1 |
| FOXP3 | TGLI1 | JAG1 |
| KCNQ4 | CD44 | GJA1 |
| JAK2 | NANOG | TNFSF11 |
| AKT1 | SOX2 | TNFSF11A |
| FOXP2 | POU5F1 | ADRB2 |
| MIR199A | SOX2 | CTNNB1 |
| MIR199A1 | FSCN1 | PPARG |
| MIR199A2 | HBEGF | NFKB1 |
| ATP11B | GJB3 | PDCD1 |
| PTDSS2 | PTK2 | TIGIT |
| BRCA1 | GJA1 | IL1B |
| CXCR3 | LAMA4 | PTHLM |
| CXCR4 | ITGA3 | EZH2 |
| CCR7 | AXL | ITGB1 |
| SPDR | TNC | PTK2 |
| CYR61 | EGFR | TGFBR1 |
| NR2F1 | VEGFA | TGFBR2 |
| WDR5 | CACNA1H | CAPG |
| IGF2BP1 | CAMK2A | STC1 |
| PTGS2 | MAPK11 | PRMT5 |
| GDF15 | MAPK14 | HGF |
| IGF2BP1 | CTNNB1 | MET |
| IL13RA2 | HMGA2 | VEGFA |
| INHBA | MIR1246 | PIK3CA |
| SMAD2 | EZH2 | ZEB1 |
| SMAD3 | SRC | FST |
| AKT1 | JUN | NOG |
| PTGS2 | CSF3 | CHRDL1 |
| ULK1 | ARG1 | IL11 |
| ATG4B | CD274 | CTGF |
| HOXB9 | EGFR | MAF |
| VEGFA | MIR128 | PTH1R |
| FGF2 | MTDH | DDX3X |
| CXCL8 | KISS1 | IRF7 |
| ANGPTL2 | CXCL12 | EDN1 |
| ERBB2 | MIR345 | DKK1 |
| TGFB1 | ATG5 | WNT5A |
| PRKAA1 | ATG7 | AXIN2 |
| PIK3CA | MMP9 | LRP5 |
| ERBB2 | CXCL8 | ABCC5 |
| TP63 | STAT3 | GPNMB |
| CDH1 | MAPK3 | MMP3 |
| GALNT14 | MAPK1 | KDM1A |
| KRAS | TP53 | ITGBL1 |
| BMP2 | CD274 | ITGA5 |
| BMP4 | AKT1 | NGF |
| FGF2 | HSF1 | ITGAV |
| FGF4 | XIST | ABL1 |
| MYC | MET | ABL2 |
| TP53 | MSN | TNFSF10 |
| RASAL2 | MIR503 | STAT5A |
| RAC1 | GABRA3 | STAT5B |
| ARHGAP24 | | MMP1 |
| MIR203 | | IL6 |
| | | WWTR1 |
| | | SIGLEC15 |
