## Supplementary Tables 1 & 2 for "CancerSTFormer enables multi-scale analysis of spot-resolution spatial transcriptomes and dissects gene and immune regulatory responses to targeted therapies"

**Supplementary Table 1: *In silico* gene perturbations (*CD274*, *CTLA4*, and *PDCD1* deletion) with the CancerSTFormer-50μm Local model**

| Affected Gene | Supporting Literature | Biomarker study | Mechanistic study | Cell Type | Support (+), or suppress (-) ICB |
|---|---|---|---|---|---|
| ***CD274* deletion** | | | | | |
| *ACAD9* | ACADL suppresses PD-L1 expression to prevent cancer immune evasion by targeting Hippo/YAP signaling in lung adenocarcinoma.[1]<br>Fatty acid oxidation fuels glioblastoma radioresistance with CD47-mediated immune evasion.[2] | | ● | | + |
| *PODNL1* | PODNL1 Methylation Serves as a Prognostic Biomarker and Associates with Immune Cell Infiltration and Immune Checkpoint Blockade Response in Lower-Grade Glioma.[3] | ● | | | - |
| *DOCK11* | Systemic Inflammation and Normocytic Anemia in DOCK11 Deficiency.[4]<br>The immunosenescence-related factor DOCK11 is involved in secondary immune responses of B cells.[5] | ● | ● | B-cell | + |
| *TSEN15* | Pan-cancer prognostic model and immune microenvironment analysis of natural killer cell-related genes.[6] | ● | | NK | - |
| *UBXN2A* | Role of Post-Translational Modifications in Colorectal Cancer Metastasis.[7] | ● | | | + |
| *GIMAP6* | A human case of GIMAP6 deficiency: a novel primary immune deficiency.[8]<br>GIMAP6 is required for T cell maintenance and efficient autophagy in mice.[9]<br>Investigation and verification of GIMAP6 as a robust biomarker for prognosis and tumor immunity in lung adenocarcinoma.[10] | ● | ● | T-cell | + |
| *SYNGAP1* | Antisense oligonucleotide modulation of non-productive alternative splicing upregulates gene expression.[11] | | ● | | +/- |
| ***CTLA4* deletion** | | | | | |
| *MYO6* | Single-Cell Sequencing Reveals PD-L1-Mediated Immune Escape Signaling in Lung Adenocarcinoma.[12]<br>MYO6 regulates Spatial Organization of Signaling Endosomes Driving AKT Activation and Actin Dynamics.[13] | | ● | | - |
| *FSTL3* | FSTL3 promotes tumor immune evasion and attenuates response to anti-PD1 therapy by stabilizing c-Myc in colorectal cancer.[14] | | ● | Treg | - |
| *LTA4H* | Radiotherapy-exposed CD8+ and CD4+ neoantigens enhance tumor control.[15]<br>LTA4H improves the tumor microenvironment and prevents HCC progression via targeting the HNRNPA1/LTBP1/TGF-β axis.[16]<br>Blocking $LTB_4$ signaling-mediated TAMs recruitment by *Rhizoma Coptidis* sensitizes lung cancer to immunotherapy.[17] | | ● | TAM | - |
| *MDM2* | MDM2-Mediated Ubiquitination Facilitates Forkhead Box P3 Stability and Positively Modulates Human Regulatory T Cell Function.[18]<br>MDM2 inhibition enhances immune checkpoint inhibitor efficacy by increasing IL-15 and MHC class II production.[19]<br>Association between MDM2/MDM4 amplification and PD-1/PD-L1 inhibitors-related hyperprogressive disease: A pan-cancer analysis.[20] | ● | ● | Treg | - |
| *ACAA1* | ACAA1 Is a Predictive Factor of Survival and Is Correlated With T Cell Infiltration in Non-Small Cell Lung Cancer.[21] | ● | | | + |
| *PINK1* | Mitochondrial and cytosolic roles of PINK1 shape induced regulatory T cell development and function.[22] | | ● | Treg | - |
| *PITPNM3* | Blocking recruitment of naive CD4+ T cells reverses immunosuppression in breast cancer.[23] | | ● | CD4.T | - |
| *SMARCD2* | Genome-wide CRISPR screens of T cell exhaustion identify chromatin remodeling factors that limit T cell persistence.[24] | | ● | CD8.T | - |
| *FAM216A* | Germline modifiers of the tumor immune microenvironment implicate drivers of cancer risk and immunotherapy response.[25] | ● | | | +/- |

| | | | | | |
|---|---|---|---|---|---|
| ***PDCD1* deletion** | | | | | |
| *GLIS3* | The gene regulatory molecule GLIS3 in gastric cancer as a prognostic marker and be involved in the immune infiltration mechanism.[26]<br>A costimulatory molecule-related signature in regards to evaluation of prognosis and immune features for clear cell renal cell carcinoma.[27] | ● | | | - |
| *KCNMB1* | Exploring of bladder cancer immune-related genes and potential therapeutic targets based on transcriptomic data and Mendelian randomization analysis.[28] | ● | | B-cell | + |
| *TMC5* | Prediction of response to immune checkpoint blockade in patients with metastatic colorectal cancer with microsatellite instability.[29]<br>Comprehensive analysis of the prognosis and immune infiltration of TMC family members in renal clear cell carcinoma.[30] | ● | | | - |
| *TTC19* | An integrated single-cell transcriptomic dataset for non-small cell lung cancer.[31] | ● | | Tnaive | - |
| *NSUN5* | Knocking down NSUN5 inhibits the development of clear cell renal cell carcinoma by inhibiting the p53 pathway.[32]<br>Single-cell and spatial transcriptomics reveal m5C RNA methylation regulators immunologically reprograms tumor microenvironment characterizations, immunotherapy response and precision treatment of clear cell renal cell carcinoma.[33] | ● | ● | Treg, Mac | - |
| *PROM2* | The prognostic significance of a novel ferroptosis-related gene model in breast cancer.[34] | ● | | | - |

Abbreviations:

ICB: Immune-checkpoint blockade

+/-: Unclear due to context-dependence or lack of report of direction

NK: Natural killer; Treg: Regulatory T cell; Mac: Macrophage; TAM: Tumor-associated macrophage; Tnaive: Naïve T cell.

**Supplementary Table 2: *In silico* gene perturbations (*CD274*, *CTLA4*, and *PDCD1* deletion) with the CancerSTFormer-250µm Extended model**

| Affected Gene | Supporting Literature | Biomarker study | Mechanistic study | Cell Type | Support (+), or Suppress (-) ICB |
|---|---|---|---|---|---|
| ***CD274* deletion** | | | | | |
| *CCL18* | BCL2A1 and CCL18 Are Predictive Biomarkers of Cisplatin Chemotherapy and Immunotherapy in Colon Cancer Patients<br>Targeting macrophages in cancer immunotherapy.[35] | ● | | Macro | - |
| *MGP* | MGP promotes CD8+ T cell exhaustion by activating the NF-κB pathway leading to liver metastasis of colorectal cancer.[36]<br>MGP+ and IDO1+ tumor-associated macrophages facilitate immunoresistance in breast cancer revealed by single-cell RNA sequencing.[37] | | ● | Fibro, Macro | - |
| *S100A9* | S100A9+CD14+ monocytes contribute to anti-PD-1 immunotherapy resistance in advanced hepatocellular carcinoma by attenuating T cell-mediated antitumor function.[38]<br>S100A9 and HMGB1 orchestrate MDSC-mediated immunosuppression in melanoma through TLR4 signaling.[39] | | ● | MDSC | - |
| *SPP1* | Single-cell and spatial analysis reveal interaction of FAP+ fibroblasts and SPP1+ macrophages in colorectal cancer.[40]<br>Evolution of myeloid-mediated immunotherapy resistance in prostate cancer.[41] | ● | ● | Fibro, Macro | - |
| *WARS* | Tryptophan potentiates CD8+ T cells against cancer cells by TRIP12 tryptophanylation and surface PD-1 downregulation.[42] | | ● | | + |
| ***PDCD1* deletion** | | | | | |
| *APOE* | Apolipoprotein E Promotes Immune Suppression in Pancreatic Cancer through NF-κB–Mediated Production of CXCL1.[43] | | ● | Fibro, Macro | - |
| *MUCL1* | MUC1-C induces PD-L1 and immune evasion in triple-negative breast cancer.[44]<br>MUC1-C integrates PD-L1 induction with repression of immune effectors in non-small-cell lung cancer.[45] | | ● | Fibro | - |
| *FN1* | Spatial transcriptomics reveals prognostically LYZ+ fibroblasts and colocalization with FN1+ macrophages in diffuse large B-cell lymphoma.[46]<br>Single-cell RNA sequencing identifies a subtype of FN1 + tumor-associated macrophages associated with glioma recurrence and as a biomarker for immunotherapy.[47] | ● | | Macro, Fibro | - |
| *SOWAHC* | Methylation-driven genes PMPCAP1, SOWAHC and ZNF454 as potential prognostic biomarkers in lung squamous cell carcinoma.[48] | ● | | | +/- |
| ***CTLA4* deletion** | | | | | |
| *YBX1* | Targeting Phosphorylation of Y-Box-Binding Protein YBX1 by TAS0612 and Everolimus in Overcoming Antiestrogen Resistance.[49] | | ● | | - |
| *CD74* | Autoimmune antibodies correlate with immune checkpoint therapy-induced toxicities.[50]<br>MIF and CD74 as Emerging Biomarkers for Immune Checkpoint Blockade Therapy.[51] | ● | | | +/- |
| *CCL19* | CCL19-producing fibroblasts promote tertiary lymphoid structure formation enhancing anti-tumor IgG response in colorectal cancer liver metastasis.[52]<br>Local injection of CCL19-expressing mesenchymal stem cells augments the therapeutic efficacy of anti-PD-L1 antibody by promoting infiltration of immune cells.[53] | | ● | Fibro, MSC | + |
| *ISG15* | Cohort-based pan-cancer analysis and experimental studies reveal ISG15 gene as a novel biomarker for prognosis and immunotherapy efficacy prediction.[54] | ● | | | - |
| *SNHG25* | Systematic analysis reveals a pan-cancer SNHG family signature predicting prognosis and immunotherapy response.[55] | ● | | | +/- |

| | | | | | |
|---|---|---|---|---|---|
| *COL1A2* | CTLA-4 blockade and interferon-α induce proinflammatory transcriptional changes in the tumor immune landscape that correlate with pathologic response in melanoma.[56] | ● | ● | ECM, Fibro | - |

Abbreviations:

ICB: Immune-checkpoint blockade

+/-: Unclear due to context-dependence, or lack of report of direction

Fibro: Fibroblast; Mac: Macrophage; MSC: Mesenchymal stem cell; MDSC: Myeloid-derived suppressor cell; ECM: Extracellular matrix (stromal cell).

### References


1. Li, L., Wang, L.-L., Wang, T.-L. & Zheng, F.-M. ACADL suppresses PD-L1 expression to prevent cancer immune evasion by targeting Hippo/YAP signaling in lung adenocarcinoma. *Medical Oncology* **40**, 118 (2023).
2. Jiang, N. *et al.* Fatty acid oxidation fuels glioblastoma radioresistance with CD47-mediated immune evasion. *Nat Commun* **13**, 1511 (2022).
3. Noor, H., Zaman, A., Teo, C. & Sughrue, M. E. PODNL1 Methylation Serves as a Prognostic Biomarker and Associates with Immune Cell Infiltration and Immune Checkpoint Blockade Response in Lower-Grade Glioma. *Int J Mol Sci* **22**, 12572 (2021).
4. Block, J. *et al.* Systemic Inflammation and Normocytic Anemia in DOCK11 Deficiency. *New England Journal of Medicine* **389**, 527–539 (2023).
5. Sugiyama, Y. *et al.* The immunosenescence-related factor DOCK11 is involved in secondary immune responses of B cells. *Immunity & Ageing* **19**, 2 (2022).
6. Li, C. *et al.* Pan-cancer prognostic model and immune microenvironment analysis of natural killer cell-related genes. *Transl Cancer Res* **13**, 1936–1953 (2024).
7. Peng, N. *et al.* Role of Post-Translational Modifications in Colorectal Cancer Metastasis. *Cancers (Basel)* **16**, 652 (2024).
8. Shadur, B. *et al.* A human case of GIMAP6 deficiency: a novel primary immune deficiency. *European Journal of Human Genetics* **29**, 657–662 (2021).
9. Pascall, J. C. *et al.* GIMAP6 is required for T cell maintenance and efficient autophagy in mice. *PLoS One* **13**, e0196504 (2018).
10. Chen, X. *et al.* Investigation and verification of GIMAP6 as a robust biomarker for prognosis and tumor immunity in lung adenocarcinoma. *J Cancer Res Clin Oncol* **149**, 11041–11055 (2023).
11. Lim, K. H. *et al.* Antisense oligonucleotide modulation of non-productive alternative splicing upregulates gene expression. *Nat Commun* **11**, 3501 (2020).
12. Zhang, A. *et al.* Single-Cell Sequencing Reveals PD-L1-Mediated Immune Escape Signaling in Lung Adenocarcinoma. *J Cancer* **16**, 1438–1450 (2025).
13. Masters, T. A., Tumbarello, D. A., Chibalina, M. V. & Buss, F. MYO6 Regulates Spatial Organization of Signaling Endosomes Driving AKT Activation and Actin Dynamics. *Cell Rep* **19**, 2088–2101 (2017).
14. Li, H. *et al.* FSTL3 promotes tumor immune evasion and attenuates response to anti-PD1 therapy by stabilizing c-Myc in colorectal cancer. *Cell Death Dis* **15**, 107 (2024).
15. Lhuillier, C. *et al.* Radiotherapy-exposed CD8+ and CD4+ neoantigens enhance tumor control. *Journal of Clinical Investigation* **131**, (2021).
16. Yang, S. *et al.* LTA4H improves the tumor microenvironment and prevents HCC progression via targeting the HNRNPA1/LTBP1/TGF-β axis. *Cell Rep Med* **6**, 102000 (2025).
17. Yan, J. *et al.* Blocking LTB4 signaling-mediated TAMs recruitment by Rhizoma Coptidis sensitizes lung cancer to immunotherapy. *Phytomedicine* **119**, 154968 (2023).
18. Wang, A. *et al.* Mouse Double Minute 2 Homolog-Mediated Ubiquitination Facilitates Forkhead Box P3 Stability and Positively Modulates Human Regulatory T Cell Function. *Front Immunol* **11**, (2020).
19. Langenbach, M. *et al.* MDM2 Inhibition Enhances Immune Checkpoint Inhibitor Efficacy by Increasing IL15 and MHC Class II Production. *Molecular Cancer Research* **21**, 849–864 (2023).

20. Ju, W. *et al.* Association between MDM2/MDM4 amplification and PD-1/PD-L1 inhibitors-related hyperprogressive disease: A pan-cancer analysis. *Journal of Clinical Oncology* **37**, 2557–2557 (2019).
21. Feng, H. & Shen, W. ACAA1 Is a Predictive Factor of Survival and Is Correlated With T Cell Infiltration in Non-Small Cell Lung Cancer. *Front Oncol* **10**, (2020).
22. Ellis, G. I., Zhi, L., Akundi, R., Büeler, H. & Marti, F. Mitochondrial and cytosolic roles of PINK1 shape induced regulatory T-cell development and function. *Eur J Immunol* **43**, 3355–3360 (2013).
23. Su, S. *et al.* Blocking the recruitment of naive CD4+ T cells reverses immunosuppression in breast cancer. *Cell Res* **27**, 461–482 (2017).
24. Belk, J. A. *et al.* Genome-wide CRISPR screens of T cell exhaustion identify chromatin remodeling factors that limit T cell persistence. *Cancer Cell* **40**, 768-786.e7 (2022).
25. Pagadala, M. *et al.* Germline modifiers of the tumor immune microenvironment implicate drivers of cancer risk and immunotherapy response. *Nat Commun* **14**, 2744 (2023).
26. Ding, Y. *et al.* The gene regulatory molecule GLIS3 in gastric cancer as a prognostic marker and be involved in the immune infiltration mechanism. *Front Oncol* **13**, (2023).
27. Hua, X. *et al.* A costimulatory molecule-related signature in regard to evaluation of prognosis and immune features for clear cell renal cell carcinoma. *Cell Death Discov* **7**, 252 (2021).
28. Xu, Z. *et al.* Exploring of bladder cancer immune-related genes and potential therapeutic targets based on transcriptomic data and Mendelian randomization analysis. *Front Immunol* **16**, (2025).
29. Ratovomanana, T. *et al.* Prediction of response to immune checkpoint blockade in patients with metastatic colorectal cancer with microsatellite instability. *Annals of Oncology* **34**, 703–713 (2023).
30. Tang, W. *et al.* Comprehensive analysis of the prognosis and immune infiltration of TMC family members in renal clear cell carcinoma. *Sci Rep* **13**, 11668 (2023).
31. Prazanowska, K. H. & Lim, S. Bin. An integrated single-cell transcriptomic dataset for non-small cell lung cancer. *Sci Data* **10**, 167 (2023).
32. Li, L., Li, M., Zheng, J., Li, Z. & Chen, X. Knocking down NSUN5 inhibits the development of clear cell renal cell carcinoma by inhibiting the p53 pathway. *Aging* https://doi.org/10.18632/aging.204761 (2023) doi:10.18632/aging.204761.
33. Gui, C.-P. *et al.* Single-cell and spatial transcriptomics reveal 5-methylcytosine RNA methylation regulators immunologically reprograms tumor microenvironment characterizations, immunotherapy response and precision treatment of clear cell renal cell carcinoma. *Transl Oncol* **35**, 101726 (2023).
34. Lu, Y.-J., Gong, Y., Li, W.-J., Zhao, C.-Y. & Guo, F. The prognostic significance of a novel ferroptosis-related gene model in breast cancer. *Ann Transl Med* **10**, 184–184 (2022).
35. Yue, T. *et al.* BCL2A1 and CCL18 Are Predictive Biomarkers of Cisplatin Chemotherapy and Immunotherapy in Colon Cancer Patients. *Front Cell Dev Biol* **9**, (2022).
36. Rong, D. *et al.* MGP promotes CD8 $^+$ T cell exhaustion by activating the NF-κB pathway leading to liver metastasis of colorectal cancer. *Int J Biol Sci* **18**, 2345–2361 (2022).
37. Chang, K. *et al.* MGP+ and IDO1+ tumor-associated macrophages facilitate immunoresistance in breast cancer revealed by single-cell RNA sequencing. *Int Immunopharmacol* **131**, 111818 (2024).
38. Tu, X. *et al.* S100A9+CD14+ monocytes contribute to anti-PD-1 immunotherapy resistance in advanced hepatocellular carcinoma by attenuating T cell-mediated antitumor function. *Journal of Experimental & Clinical Cancer Research* **43**, 72 (2024).
39. Özbay Kurt, F. G. *et al.* S100A9 and HMGB1 orchestrate MDSC-mediated immunosuppression in melanoma through TLR4 signaling. *J Immunother Cancer* **12**, e009552 (2024).
40. Qi, J. *et al.* Single-cell and spatial analysis reveal interaction of FAP+ fibroblasts and SPP1+ macrophages in colorectal cancer. *Nat Commun* **13**, 1742 (2022).
41. Lyu, A. *et al.* Evolution of myeloid-mediated immunotherapy resistance in prostate cancer. *Nature* **637**, 1207–1217 (2025).
42. Qin, R. *et al.* Tryptophan potentiates CD8 $^+$ T cells against cancer cells by TRIP12 tryptophanylation and surface PD-1 downregulation. *J Immunother Cancer* **9**, e002840 (2021).

43. Kemp, S. B. *et al.* Apolipoprotein E Promotes Immune Suppression in Pancreatic Cancer through NF-κB–Mediated Production of CXCL1. *Cancer Res* **81**, 4305–4318 (2021).
44. Maeda, T. *et al.* MUC1-C Induces PD-L1 and Immune Evasion in Triple-Negative Breast Cancer. *Cancer Res* **78**, 205–215 (2018).
45. Bouillez, A. *et al.* MUC1-C integrates PD-L1 induction with repression of immune effectors in non-small-cell lung cancer. *Oncogene* **36**, 4037–4046 (2017).
46. Dai, L. *et al.* Spatial transcriptomics reveals prognostically LYZ+ fibroblasts and colocalization with FN1+ macrophages in diffuse large B-cell lymphoma. *Cancer Immunology, Immunotherapy* **74**, 123 (2025).
47. Xu, H. *et al.* Single-cell RNA sequencing identifies a subtype of FN1 + tumor-associated macrophages associated with glioma recurrence and as a biomarker for immunotherapy. *Biomark Res* **12**, 114 (2024).
48. Zhu, Q. *et al.* Methylation-driven genes PMPCAP1, SOWAHC and ZNF454 as potential prognostic biomarkers in lung squamous cell carcinoma. *Mol Med Rep* https://doi.org/10.3892/mmr.2020.10933 (2020) doi:10.3892/mmr.2020.10933.
49. Shibata, T. *et al.* Targeting Phosphorylation of Y-Box–Binding Protein YBX1 by TAS0612 and Everolimus in Overcoming Antiestrogen Resistance. *Mol Cancer Ther* **19**, 882–894 (2020).
50. Tahir, S. A. *et al.* Autoimmune antibodies correlate with immune checkpoint therapy-induced toxicities. *Proceedings of the National Academy of Sciences* **116**, 22246–22251 (2019).
51. Fey, R. M., Nichols, R. A., Tran, T. T., Vandenbark, A. A. & Kulkarni, R. P. MIF and CD74 as Emerging Biomarkers for Immune Checkpoint Blockade Therapy. *Cancers (Basel)* **16**, 1773 (2024).
52. Zhang, Y. *et al.* CCL19-producing fibroblasts promote tertiary lymphoid structure formation enhancing anti-tumor IgG response in colorectal cancer liver metastasis. *Cancer Cell* **42**, 1370-1385.e9 (2024).
53. Iida, Y. *et al.* Local injection of CCL19-expressing mesenchymal stem cells augments the therapeutic efficacy of anti-PD-L1 antibody by promoting infiltration of immune cells. *J Immunother Cancer* **8**, e000582 (2020).
54. Wei, J. *et al.* Cohort-based pan-cancer analysis and experimental studies reveal ISG15 gene as a novel biomarker for prognosis and immunotherapy efficacy prediction. *Cancer Immunology, Immunotherapy* **74**, 168 (2025).
55. Zheng, H. *et al.* Systematic analysis reveals a pan-cancer SNHG family signature predicting prognosis and immunotherapy response. *iScience* **26**, 108055 (2023).
56. Khunger, A. *et al.* CTLA-4 blockade and interferon-α induce proinflammatory transcriptional changes in the tumor immune landscape that correlate with pathologic response in melanoma. *PLoS One* **16**, e0245287 (2021).
