## Supplementary Figure 1 for "CancerSTFormer enables multi-scale analysis of spot-resolution spatial transcriptomes and dissects gene and immune regulatory responses to targeted therapies"

**Supplementary Fig 1**

**a**

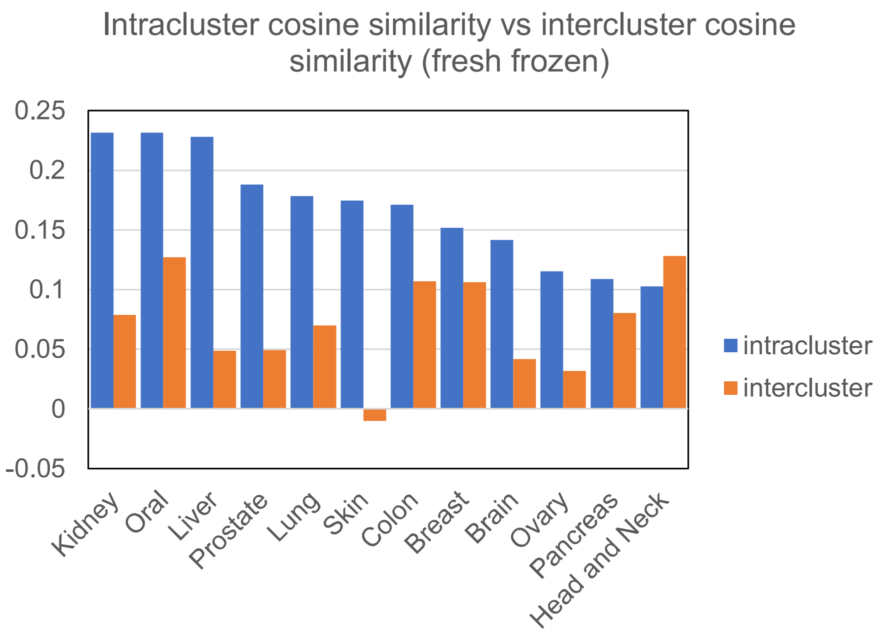


**b**

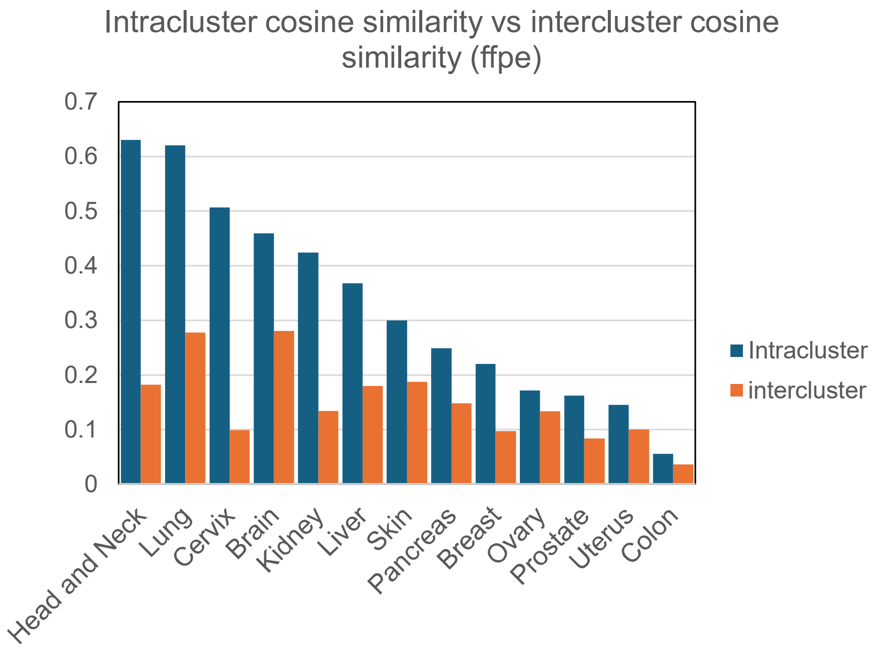


Evaluation of pretrained embedding's cluster coherence by measuring intracluster vs intercluster cosine similarity. For ST samples marked with each cancer type (x-axis), we measured the samples' intracluster vs intercluster cosine similarity (to the closest cluster), and plot the similarity values. Higher intracluster cosine similarities than intercluster similarities indicate better cluster coherence. **a.** Limited to fresh-frozen ST samples only. Head and Neck carcinoma has a higher intercluster score, as it is indistinguishable from oral carcinoma, both of which are of squamous origin. **b**. Limited to FFPE samples only.
