## Supplementary Figure 2 for "CancerSTFormer enables multi-scale analysis of spot-resolution spatial transcriptomes and dissects gene and immune regulatory responses to targeted therapies"

**Supplementary Fig 2**

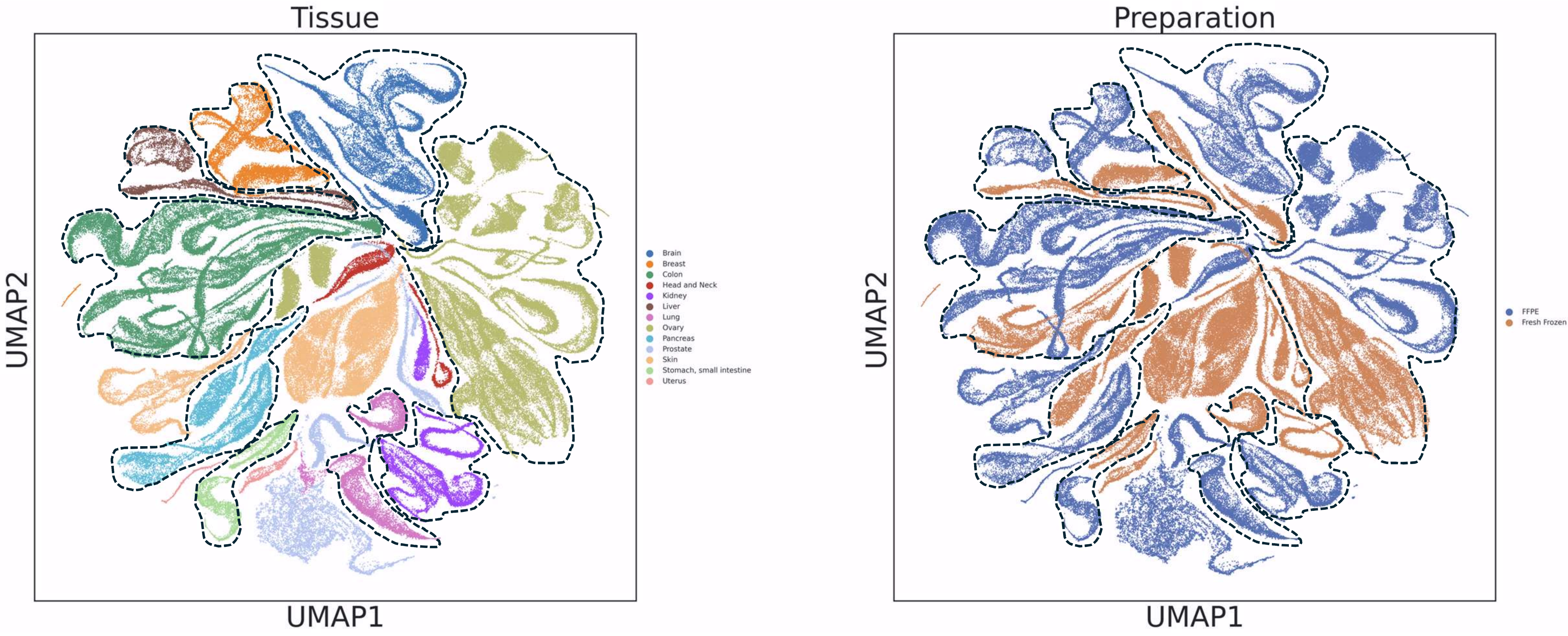


Application of Combat on pretrained embeddings removes sample preparation related batch effects. Post-correction results shown. Left: post-correction embedding, colored by tissue types. Right: same embedding colored by sample preparation (FFPE vs fresh frozen). Dashed lines delineate tissue type clusters (see left) which cluster together regardless of sample preparation (see right).
