## Supplementary Figure 3 for "CancerSTFormer enables multi-scale analysis of spot-resolution spatial transcriptomes and dissects gene and immune regulatory responses to targeted therapies"

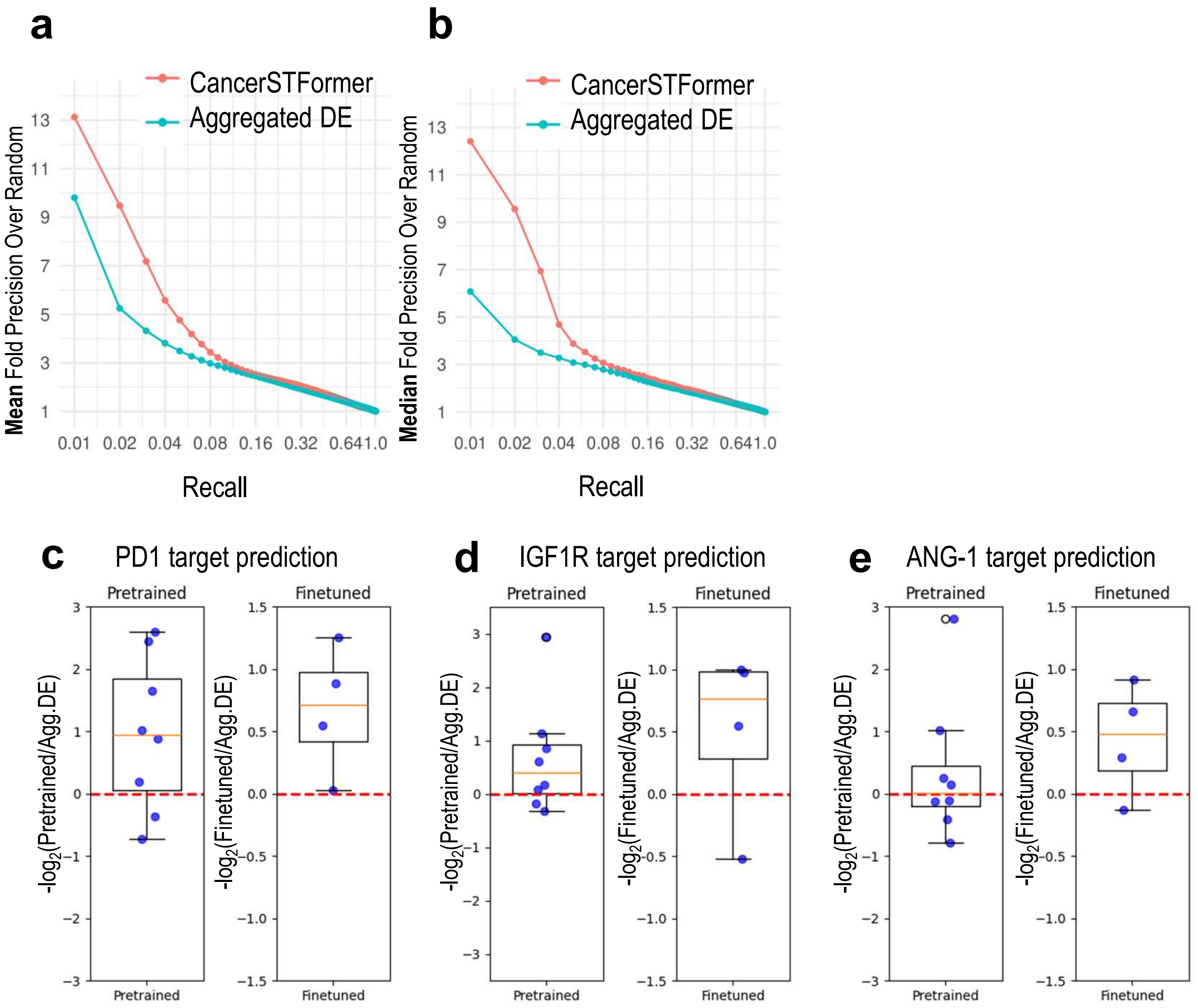


Comparison between CancerSTFormer and a baseline method. Aggregated DE merges ligand-hi vs ligand-lo niche-DE genes across many ST samples using Stouffer's P-value aggregation. CancerSTFormer was same as used in Figure 3. Ligand-target evaluation presented on the metrics the mean (**a.**) and the median (**b.**) fold-precision-over-random. **c-e.** Comparisons expressed as log2 fold-change in performance (FPOR@0.01) between pretrained model and aggregated DE, or finetuned model and aggregated DE in all the treatment prediction results presented in Figure 5.
