## Supplementary Figure 4 for "CancerSTFormer enables multi-scale analysis of spot-resolution spatial transcriptomes and dissects gene and immune regulatory responses to targeted therapies"

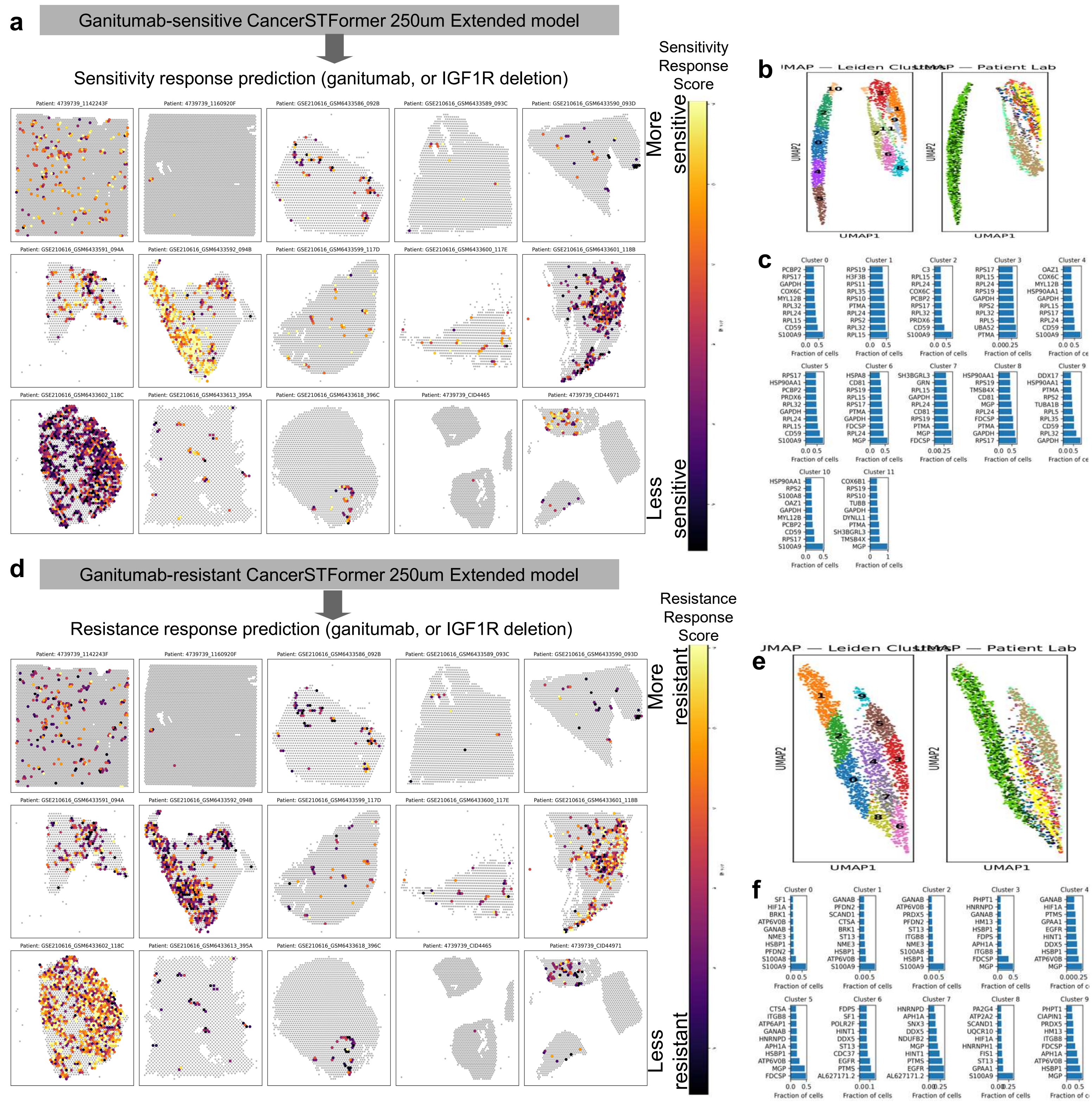


Spatial visualization of ganitumab treatment responses using CancerSTFormer 250μm Extended model. **a.** Prioritization of the TNBC ST samples based on sensitivity signatures to ganitumab therapy (IGF1R-targeting). Sensitivity score is calculated using the finetuned 250um-Extended model that was trained to recognize ganitumab sensitivity. **b.** UMAP of ganitumab responses exhibited by all IGF1R expressing spots. 12 spot clusters were derived based on UMAP. **c.** Top sensitive signatures returned by the model per spot cluster. X-axis: sensitivity score. **d.** Prioritization of TNBC ST samples based on ganitumab resistance. **e.** UMAP plot, overlayed with cluster labels and patient labels.10 spot clusters were derived. **f.** Top resistance markers per cluster. X-axis: resistance score. Results show that 1142243F and GSM6433592 are highly sensitive while GSM6433601 and GSM6433602 are highly resistant.
